# Mammalian Proteome Profiling Reveals Readers and Antireaders of Strand-Symmetric and -Asymmetric 5-Hydroxymethylcytosine-Modifications in DNA

**DOI:** 10.1101/2025.06.27.661915

**Authors:** Lena Engelhard, Zeyneb Vildan Cakil, Simone Eppmann, Tye Gonzalez, Rasmus Linser, Petra Janning, Daniel Summerer

## Abstract

The cytosine (C) modifications 5-methylcytosine (mC) and 5-hydroxymethylcytosine (hmC) are central regulatory elements of mammalian genomes. Both marks occur in dinucleotide contexts of double stranded DNA in either strand-symmetric or -asymmetric fashion, but it is still poorly understood how this symmetry information is selectively read out by the nuclear proteome as basis of potential symmetry-dependent regulation mechanisms. We report comparative enrichment/proteomics studies with promoter probes being strand-symmetrically or asymmetrically modified with C, mC and hmC, enabling a direct assessment of their reader profiles in the same sequence, tissue, and experimental contexts. We identify in human and mouse nuclear lysates a high number of tissue-specific readers for hmC-modified sequences that fall into distinct, probe-specific sub-groups, including members of important transcription factor classes and chromatin regulators. Among them, we discover the master regulators MYC and MAX that play central roles in cell (de)differentiation and cancer progression to read hmC in a sequence-dependent manner. We also find RFX5, a transcription factor critically involved in primary MHC class II deficiency to discriminate between specific hmC symmetries in CpG dyads. Our findings provide further support for the hypothesis that hmC symmetry information can provide distinct regulatory outputs, and provide a resource for studying the molecular mechanisms triggered by symmetric and asymmetric hmC modifications in chromatin regulation during development and disease.

## 1. Introduction

The epigenetic DNA modification 5-methylcytosine (mC) is a central regulator of mammalian gene expression with crucial roles in development, differentiation, and cancer formation^1^. mC is written and maintained by DNA methyltransferases (DNMTs) predominantly within palindromic CpG dyads, of which 60-80 % are methylated in somatic cells^2^. mC can be recognized and converted into transcriptionally repressive outputs by methyl-CpG-binding domain (MBD) reader proteins^3^, but can also recruit or repel transcription factors^4^. CpG methylation can further be actively reversed by the action of ten-eleven translocation (TET) dioxygenases, which catalyze the step-wise oxidation of mC to 5-hydroxymethylcytosine (hmC), 5-formylcytosine (fC) and 5-carboxycytosine (caC) (**Fig. 1a**). fC and caC can subsequently be restored to C via the thymine-DNA glycosylase (TDG)-initiated base excision repair (BER) pathway^5,6^. The oxidized mC derivatives differ significantly from mC in their chemical properties, genomic distributions and dynamic global changes during development, and thus have potential to differently regulate chromatin-associated processes via selective recruitment or repulsion of regulatory proteins^5^. In particular, hmC can exhibit high stability^7^, and occurs at high levels in embryonic stem cells (ESC) and neurons, where it is enriched in promoters and gene bodies^8–10^.

**Figure 1.**
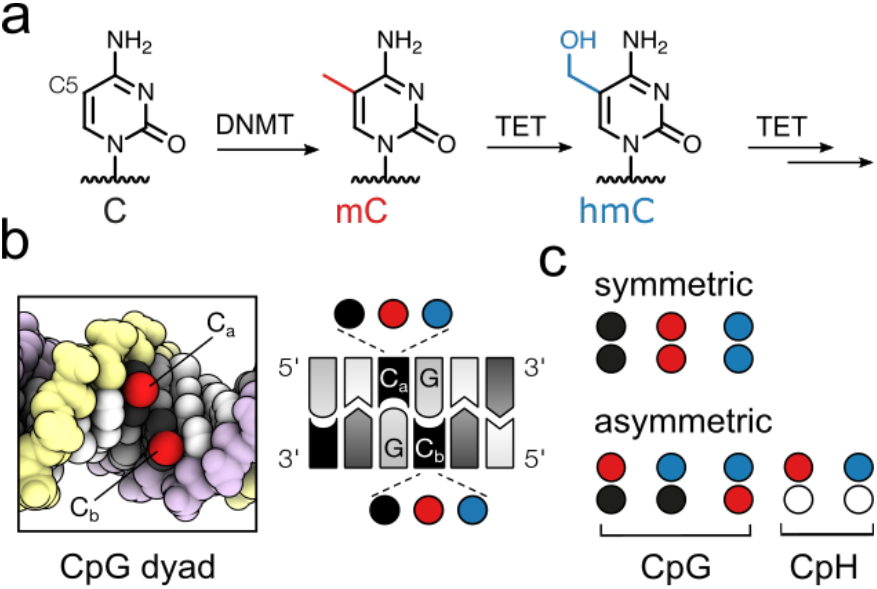
Mammalian cytosine modifications in doublestranded DNA. (a) The cytosine modifications mC and hmC are written by DNMT and TET enzymes. (b) In CpG dyads, both cytosines (C_a_, C_b_) can exist as C, mC or hmC. (c) Strand-symmetric and -asymmetric versions of CpG and CpH dinucleotides containing C, mC and hmC (white circle: T, A, or G).

Importantly, hmC can co-exist in symmetric and asymmetric combinations with C, mC and itself in the two strands of CpG dyads, giving rise to six combinatorial marks that are presented in the DNA major groove^11^, a preferred interaction surface of DNA-binding proteins (**Fig. 1b, c**)^5^. Previous mapping studies found that the dyad hmC/mC (for top and bottom-strand, respectively) accounts for the majority of all hmC-containing CpG dyads (>70% in mouse cerebellum)^12,13^, whereas hmC/hmC and hmC/C dyads occur 4-6-fold less frequently, respectively. In addition, mC can occur at higher levels in inherently asymmetric CpH dinucleotides, which are found to be further oxidized to hmCpH to smaller, but not negligible levels in a tissue-specific manner (**Fig. 1c**)^14,15^.

Previous studies with individual chromatin factors such as UHRF1/2^16,17^, MBDs^18,19^, SALL1/4^20^, and others have revealed that they interact differently with hmC and mC in vitro (reviewed in^21,22^), partially being able to differentiate between different hmC CpG symmetries. Similarly, specific preferences for symmetrically or hemi-modified hmC CpG dyads have been observed for a collection of recombinant transcription factor constructs via in vitro selections^23^. These studies provide insights into direct hmC interactions of individual recombinant proteins, providing biochemical support for the hypothesis that hmC may regulate chromatin processes in a symmetry-specific manner. A particularly powerful approach for the discovery of (anti)readers and potential new functions of hmC are enrichment/proteomics studies with natural nuclear extracts. Such studies allow system-level analyses and, unlike in vitro binding studies with recombinant proteins, take into account the complexity of mammalian chromatin, in which protein-DNA interactions are controlled by diverse additional protein or RNA complex partners, ligands and post-translational modifications in a complex and dynamic manner. In this direction, previous proteomics studies using symmetric dsDNA enrichment probes containing symmetrically modified CpG dyads (e.g., hmC/hmC) have afforded unique reader profiles for each C modification^20,24–26^.

To extend these efforts and to systematically study the role of hmC-modification symmetry in DNA-proteome interactions, we here report proteomics screens covering symmetric and asymmetric hmC-modifications to compare their interactomes in the same sequence, tissue, and experimental context. Employing human and mouse nuclear extracts and DNA promoter probes that are modified with C, mC or hmC in a strand-symmetric or -asymmetric fashion, we identify comparable numbers of readers for all three hmC modified-probes (hmC/C, hmC/mC and hmC/hmC) as for C/C and mC/mC probes combined (containing the frequent, canonical C/C or mC/mC CpGs). The hmC readers thereby fell into distinct subgroups for each of the three hmC probes, with the majority of human readers binding to asymmmetric hmC/C and hmC/mC probes, and not hmC/hmC probes (as previously used). By further validation studies, we identify diverse readers with distinct hmC probe symmetry preferences, among them members of the RFX proteins, FOX proteins, and the master regulators MYC and MAX. This provides further support for the view that hmC (a)symmetry plays a role in mammalian protein-DNA interactions, with possible functions in chromatin regulation.

## 2. Results and Discussion

### Generation of asymmetric DNA probes and establishment of enrichment protocol

For the discovery of readers of (a)symmetric C modifications, we established an enrichment protocol employing 5’-biotinylated, double-stranded DNA probes containing the target modifications immobilized to streptavidin-coated magnetic beads.

Briefly, the bead-bound probes were incubated with nuclear extracts, washed and subjected to western blots or were processed for nanoHPLC-MS/MS using an Ultimate 3000 RSLC nano-HPLC system and a Hybrid-Orbitrap mass spectrometer (**Fig. 2a**).

**Figure 2.**
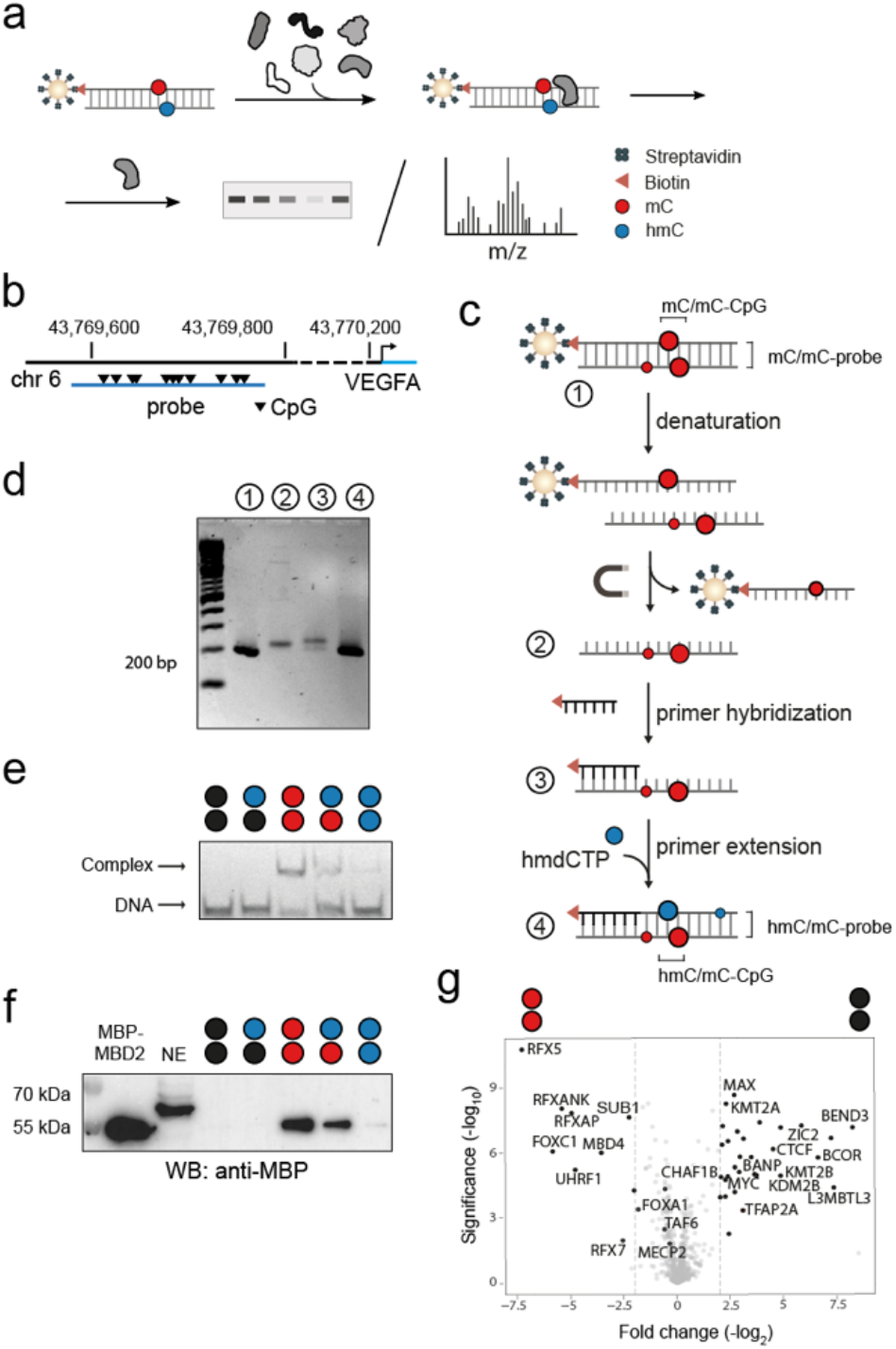
Asymmetric DNA probe generation and validation of enrichment protocol with nuclear extracts from HEK293T cells. (a) Identification of (anti)readers of symmetric and asymmetric C-modifications by enrichment and blots or LC-MS/MS analysis. (b) Genomic location of the 201 nt human VEGFA DNA promoter probe containing 11 CpGs. (c) Workflow of asymmetric probe generation exemplified for hmC/mC probes. (d) Agarose gel validation of VEGFA probe generation steps shown in **Fig. 2c**. (e) EMSA analysis of MBP-MBD2 binding to synthetic oligonucleotide probes containing a single one of the indicated CpG dyads (67 nM protein, 2 nM DNA) in an oligo-dA context. (f) Anti-MBP western blot of enrichment eluates from HEK293T nuclear extracts (NE) containing MBP-MBD2 using human VEGFA probes with the indicated strand modifications generated as shown in **Fig. 2c** (note that a higher molecular weight off-target protein is bound by this antibody). (g) Volcano plot (five technical replicates) comparing enriched proteins (p-value < 0.025 and log2-fold change >1.5) between mC/mC and C/C probes.

We considered multiple parameters in the design of our probes that crucially determine the enrichment results. For example, to offer enrichment of a reasonable diversity of sequence-specific proteins, a probe needs to be of sufficient length and sequence complexity (the use of probes with random sequence contexts around CpGs for enriching a maximal number of proteins is less informative, see below and **Fig. S1-4**). Ideally, the probe sequence occurs in the genome to provide a natural spatial arrangement of protein binding sites, and it is involved in important, mC-dependent regulatory processes. Whereas the practical use of (highly hmC-modified) synthetic oligonucleotide probes is cost-prohibitive and restricted by length limitations^24^, PCR-generated probes can fulfill all aforementioned criteria and are thus the current standard of proteomics studies^20,25,26^. PCR probes thereby contain modifications in CpG and CpH contexts^20,25,26^ and offer the discovery of readers for both marks (this needs to be taken into account during data interpretation^20,25,26^). We designed a probe covering ~200 nt of the human vascular endothelial growth factor (VEGF) A promoter region (**Fig. 2b**) containing a high number of transcription factor binding sites (see **Table S3** for JASPAR motif analysis). Both promoter CpG methylation and TET3 are involved in the regulation of VEGFA expression, suggesting a role of hmC-containing CpG dyads^27^. Moreover, VEGFA is a pivotal regulator of angiogenesis^28^, and aberrant VEGFA promoter methylation has been associated with pathological outcomes, such as various types of cancer and inflammation^29–31^. To obtain strand-asymmetrically modified DNA probes, we first generated probes by PCR using a 5’-biotinylated primer and either dCTP, dmCTP or dhmCPT in the dNTP mix, leading to C/C, mC/mC and hmC/hmC probes (**Fig. 2c**). We immobilized the PCR product to streptavidin-coated magnetic beads and recovered the non-biotinylated reverse single strand via alkaline denaturation with 20 mM NaOH (**Fig. 2c** and **Fig. S5**). We hybridized a 5’-biotinylated primer to the reverse strand, and conducted a primer extension employing dATP, dTTP, dGTP and either dCTP, dmCTP or dhmCTP, leading to biotinylated hmC/hmC, hmC/mC and hmC/C probes bearing the desired symmetric and asymmetric hmC CpG dyads (**Fig. 2c**; **Fig. 2d** shows agarose gel analyses of the individual steps). We further confirmed that the final probes are fully double-stranded (**Fig. S6-7, 9**) and verified their sequence integrity and quantitative modification (**Fig. S8**).

To validate the function of our probes, we conducted control enrichments with human methyl-CpG binding domain protein 2 (MBD2) as a validated mC/mC CpG reader. MBD2 does not bind C/C and hmC/C CpGs, but exhibits increasing affinities in the order hmC/hmC << hmC/mC < mC/mC)^18,19^, making it ideally suited to evaluate, if our enrichment protocol is able to reveal subtle CpG dyad preferences of candidate proteins (**Fig. 2e**). For the enrichments, we added 1 nM of MBD2 bearing an N-terminal maltose binding protein (MBP) tag to nuclear extracts (NE) obtained from human embryonic kidney (HEK) 293T cells, and incubated this mix with the VEGFA DNA probes immobilized on streptavidin-coated magnetic beads. After washing steps, we eluted and analyzed the probe-bound protein by anti-MBP western blots. In agreement with the EMSA data, MBP-MBD2 was enriched only for the mC/mC and to a lesser extent the hmC/mC probes, indicating that our enrichment protocol is capable of profiling protein DNA interactions with sufficient selectivity for differentiating chemically similar CpG dyads (**Fig. 2f**).

### Establishment of enrichment/MS-based proteomics for discovering (anti)reader profiles of cytosine modifications

We next combined enrichments with quantitative HPLC-MS/MS to identify (anti)reader proteins. We conducted enrichments with HEK293T nuclear extracts in two independent biological experiments (a third biological experiment was conducted for a subset of probes under technically slightly different conditions; **Table S1** and **Fig. S10-12**). These experiments provide insights into how (anti)reading events may vary biologically (e.g., due to changes in expression levels, modification, localization etc.). For statistical significance analyses, we conducted five technical replicates for each biological experiment, allowing for robust identification of (anti)readers.

We first evaluated our setup by comparative proteome enrichments with symmetric C/C and mC/mC probes containing the frequent, canonical C/C and mC/mC CpGs, which we expected to afford a larger number of known (anti)readers as a reference. Indeed, we observed a significant enrichment (p-value < 0.025 and log2-fold change >1.5) of many known mC readers for the mC/mC probe, most notably MBD proteins^3^ (MBD2, MeCP2, MBD4), UHRF1, and RFX proteins^32^ (such as RFX5, RFX7, RFXANK and RXFAP (**Fig. 2g, Fig. S10** and **Table S2a**). These proteins partially overlap with enriched proteins in previous proteomics studies^20,24–26^, although the use of different tissues and/or probe sequences in these studies allow only limited comparisons. We also observed an enrichment of many proteins for the C/C over the mC/mC probe, among them CXXC-domain containing proteins like KMT2A and KDM2B (the latter together with other components of the noncanonical polycomb repressive complex PRC1.1, such as PCGF1 and BCOR), and other C/C readers such as CTCF, MYC and MAX (**Fig. 2g, Table S2a**).

To explore the possibility of maximizing the number of enriched readers, we also designed probe pools containing CpG dyads in random sequence contexts (NNNNCGNNNN), covering a large number of CpG-containing target sequences. Probes with a central C/C or mC/mC CpG enriched only a smaller subset of readers compared to the VEGFA probe (**Fig. S1**). Since these two probe designs differ in the absolute number of CpGs, we also designed and tested random probes with additional repeats of the NNNNCGNNNN sequence (**Fig. S2**). A probe with three such repeats enriched more readers, but still less than the VEGFA probe, and the enrichment was mainly limited to non-sequence-specific readers such as MBD proteins (**Fig. S3-4**). This data suggests that the lower concentrations of individual probe sequences in random probe pools prevent effective enrichment of sequence-specific readers, confirming that probes with a single, complex sequence (such as our VEGFA probe) are the design of choice.

### Distribution of reader numbers among symmetrically and asymmetrically modified probes

We initially assessed the overall number of enriched readers for each probe, since a common observation in previous proteomics studies has been an overall low number of hmC readers as compared to C and mC readers^20,24–26^. However, given the exclusive use of probes containing symmetrically modified CpGs in these studies, this finding is only valid for the isolated case of hmC/hmC CpGs (and hmCpH). In our experiments, we identified 62, 42 and 16 proteins that were significantly enriched for C/C, mC/mC, and hmC/hmC probes in at least one biological experiment, respectively (**Fig. 3a**, see **Fig. S13** for distribution of individual biological experiments). Although this distribution somewhat differs from previous reports by a relatively higher number of mC/mC probe readers, the data show the same trend. Strikingly, we also identified a total of 42 and 15 readers for asymmetric hmC/C and hmC/mC probes that have not been employed in previous proteomics studies – many more than for the known symmetric hmC/hmC probes. Importantly, the hmC readers thereby fell into distinct, symmetry-specific sub-groups, which may reflect preferences of many proteins for specific hmC-CpG dyads (**Fig. 3a-b**).

**Figure 3:**
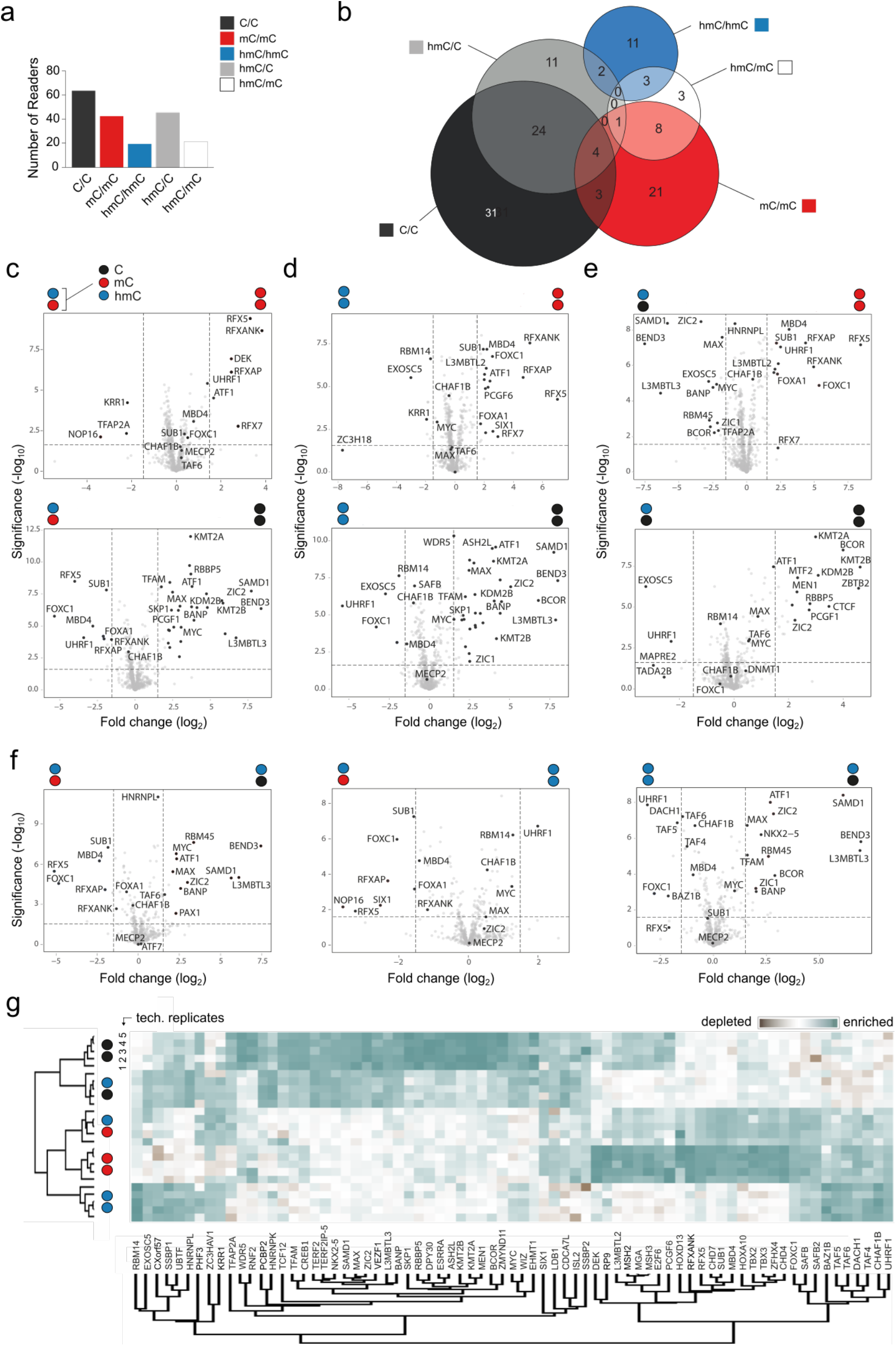
Discovery of readers and antireaders of symmetrically and asymmetrically hmC-modified DNA probes from human HEK293T cells by enrichment/proteomics. (a) Overall number of enriched reader proteins for indicated DNA probes. (b) Venn diagrams for significantly enriched proteins for indicated probes (note that color code refers to both DNA strands); see **Table S4** for protein names. (c-f) Volcano plots from pairwise probe comparisons for one exemplary biological experiment (p-value < 0.025 and log2-fold change >1.5; see **Fig. S10** and **Table S2** for additional experiments and **Fig. S11** for reproducibility data). (g) Heatmap of correlation-based clustering of the label-free quantification (LFQ) intensities after log2 transformation and normalization by row mean subtraction. Proteins included in the clustering significantly bind to at least one of the probes as determined by an ANOVA test (data from the biological experiment of **Fig. 3c-f**, see **Fig. S12** for heatmaps of other experiments).

### Readers and antireaders of hmC/mC probes

We next assessed the individual (anti)readers for our different hmC-modified probes including their overlaps. **Fig. 3b** shows Venn diagrams of readers for each probe identified in at least one of the biological experiments (see also **Table S4** and **Fig. S14** for other experiments). **Fig. 3c-f** show the volcano plots for pairwise comparisons of the indicated probes for one biological experiment, and **Fig. 3g** shows the respective heat map generated through hierarchical clustering of the mass spectral counts for the most significantly enriched or depleted proteins in pairwise comparisons (see **Fig. S10-12** and **Table S2** for additional biological experiments and full hit tables). To qualify as reader in the below discussion, a protein needed to be significantly enriched in at least four of five technical replicates in at least one of the biological experiments (i.e., some readers appear only in **Fig. S10**, not **Fig. 3**). Among the proteins that were enriched for mC/mC over hmC/mC probes (most of which being also enriched over C/C) were numerous RFX-family proteins (e.g., RFX1, RFX7, RFX5 and its interaction partner RFXANK), MBD2, and DEK (**Fig. 3c, Table S2b**). Conversely, we found KRR1, NOP16 and TFAP2 to be enriched for hmC/mC over mC/ mC probes. TFAP2, a transcription factor that previously displayed preferential binding to C and mC probes^23,26^, has been linked to enhanced transcriptional activation by interfering with NuRD^33^ and is an interactor of MYC^34^. Comparisons between hmC/mC and C/C probes afforded proteins strongly enriched for hmC/mC (**Fig. 3c** and **Table S2c**). All of them were also enriched for mC/mC over C/C probes (**Fig. 2g**), but only some of them were weakly enriched for mC/mC over hmC/mC probes, suggesting that most of these proteins act as dual readers of mC and hmC marks. This was the case for FOXC1/A1, UHRF1, as well as the MBDs MBD4 and MECP2 (**Fig. 3c**; note that MECP2 has been shown to also bind to hmCpA dyads, which are present in PCR/primer extension-generated probes^35,36^). Both FOXC1 and UHRF1 previously showed binding to symmetric mC and hmC CpGs^17,24^. Overall, our enrichments with hmC/mC probes suggest that many classical methylation readers bind with reduced affinity to hmC, whereas some are dual readers that strongly prefer both marks over non-modified C.

### Readers and antireaders of hmC/hmC probes

We obtained a somewhat related picture for enrichments with hmC/hmC probes. Few proteins were enriched for hmC/hmC over mC/mC or C/C probes, with proteins enriched over C/C being partially found also enriched for hmC/mC and mC/mC over C/C probes, and they thus seem to act as more universal mC and hmC readers. These included UHRF1 and MECP2 (both of which have been identified to bind to hmC/hmC CpGs in previous proteomics studies using mouse ESC nuclear extracts and (CGA)_n_ repeat probes^24^), as well as FOXC1, EXOSC5 and CHAF1A/B proteins (**Fig. 3d** and **Table S2d-e**). CHAF1B, a master epigenetic transcription regulator^37^ capable of gene silencing via the displacement of activators such as CE-BPα^38^ has not yet been demonstrated to have a preference for hmC. The proteins enriched for mC/mC over hmC/hmC probes overlapped with the ones enriched over hmC/mC (predominantly RFX proteins, compare **Fig. 3c-d**).

### Readers and antireaders of hmC/C probes

Experiments with hmC/C probes afforded proteins enriched over mC/mC probes that constituted a subset of the proteins enriched for C/C over mC/mC probes (**Fig. 3e**; e.g., BEND3, SMAD1, L3MBTL3, components of the PRC1.1 complex such as BCOR and KDM2B, as well as the master transcription factors MYC and MAX, **Table S2f**). Conversely, the proteins enriched for mC/mC over hmC/C probes largely overlapped with the ones enriched for mC/mC over C/C (compare **Fig. 3e** and **2g**). For hmC/C over C/C, we found only few proteins to be enriched that were also found in hmC/mC and hmC/hmC over C/C comparisons (MECP2, UHRF1 and EXOSC5, **Fig. 3e** and **Table S2g**). Conversely, many proteins were enriched for C/C over hmC/C probes that strongly overlapped with the ones enriched for C/C over mC/mC, suggesting that many readers of nonmodified Cs are repelled by (hydroxy)methylation of their target sequences. In comparison, fewer proteins were found enriched for C/C over hmC/mC or hmC/hmC probes. For the case of CpG dyads, this observation fits to a model in that repulsion of C readers is increased by additional modification of the dyad’s antisense C with either mC or hmC.

### Reader and antireader preferences among symmetric and asymmetric hmC probes

A striking observation from the enrichment comparisons is that the readers of C/C and mC/mC probes are well separated, and that the hmC/C and hmC/mC probe readers partially overlap with the C/C and mC/mC probes, respectively (**Fig. 3b**). In contrast, hmC/hmC probes show hardly any overlap, in agreement with the low number of hmC readers identified in previous studies employing symmetric hmC/hmC probes^24,25^. As a consequence, each of the hmC-containing probes attracts a very different reader group, which – in view of CpG dyads – fits to a model in that the (a)symmetry information of the different hmC dyads is differently interpreted by nuclear proteins as potential initiating event of dyad-specific regulatory processes.

### Discovery of (anti)readers of cytosine modifications from mouse brain tissue

To investigate (anti)readers of hmC (a)symmetry information in cells that show particularly high hmC levels, we conducted proteomics experiments using mouse brain tissue^6^. We designed a second enrichment probe covering 240 nt of the mouse Sp1 promoter, and containing 17 CpGs in different arrangement/sequence contexts as in the VEGFA probe (**Fig. 4a**, see **Table S3** for JASPAR motif analysis). Sp1 is a central transcription factor involved in tumorigenesis, proliferation and differentiation^39^ and is tightly regulated over the cell cycle, with TET1 overexpression leading to Sp1 upregulation^40^. We again confirmed the ability of this probe to report preferences of the mC/mC-CpG reader protein MBD2 among chemically similar CpG dyads (**Fig. 4b**, compare to **Fig. 2e-f**), and conducted enrichment/proteomics experiments with C/C and mC/mC probes (**Fig. 4c**).

**Figure 4.**
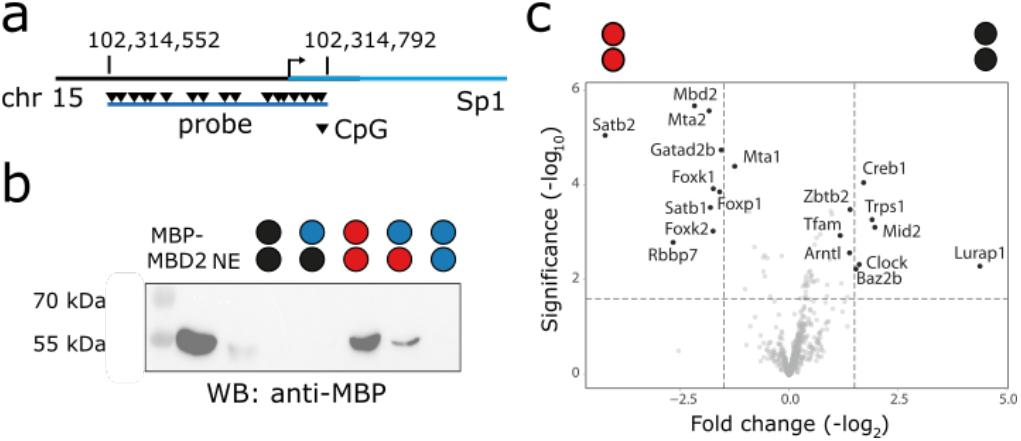
Mouse Sp1 promoter probe design and validation with nuclear extracts from mouse brain tissue. (a) Genomic location of the mouse Sp1 promoter probe. (b) Anti-MBP western blot of enrichment eluates from mouse brain nuclear extracts (NE) containing MBP-MBD2 using Sp1 probes with the indicated strand modifications generated as shown in **Fig. 2c**. (c) Volcano plot from enriched protein comparison (five technical replicates, p-value < 0.025 and log2-fold change >1.5) for mC/mC and C/C probes.

The experiment confirmed the enrichment of known mouse C and mC readers, with, e.g., Rbbp7, Mta2, endogenous Mbd2 as well as Forkhead proteins like Foxk1 and 2 being enriched for the mC/mC probe, and e.g., Creb1 being enriched for the C/C probe^24,25^ (**Fig. 4c**). We extended our proteomics experiments to the three hmC-modified versions of the Sp1 probe and observed a related overall reader distribution among the five probes as compared to the human VEGFA experiments (**Fig. 5a**, compare to **Fig. 3a** and **Fig. S13**). The largest number of readers was enriched for the C/C and mC/mC probes. Moreover, the overlap between the hmC-probes and C/C as well as mC/mC probes was similar, and again, the three hmC probes attracted different reader groups (**Fig. 5b**). Interestingly, despite this similar overall distribution, the specific identified reader candidates were largely different between human and mouse (**Fig. 5c-f, Table S2**). For example, the group being enriched for hmC/mC over other probes included factors like Srsf11, Lrrfp2 and Rbbp7 and the homeodomain transcription factor Satb2, all being proteins not identified in the human experiments (**Fig. 5c, f-g, Table S2b-c**). In the hmC/C group, we identified Mid2, Hnrnpk, Rplp1, Haus3 and the zinc finger transcription factor Trps1 (**Fig. 5e-g, Table S2f-i**). Some of these proteins overlapped with the ones found for hmC/hmC, whereas Uhrf2^16^ and Sfpq were uniquely enriched for this symmetric probe (**Fig. 5d, f-g**). Proteins that overlapped with the human experiments were mainly those enriched for the symmetric C/C or mC/mC probes containing the canonical C/C or mC/mC CpG dyads, such as the aforementioned orthologs of mC/mC reader Mbd2 and leucine zipper transcription factor Creb1, but also different members of the same families observed for human, such as forkhead domain transcription factors Foxp1 and Foxk1-2 (FoxA1/C1 in human), and the DNA binding BEN-domain containing chromatin factor Bend6 (Bend3 and BANP in human; **Fig. 5c-e**). BANP has been identified as a selective C/C reader^24,25^ that is inhibited by methylation^41^.

**Figure 5.**
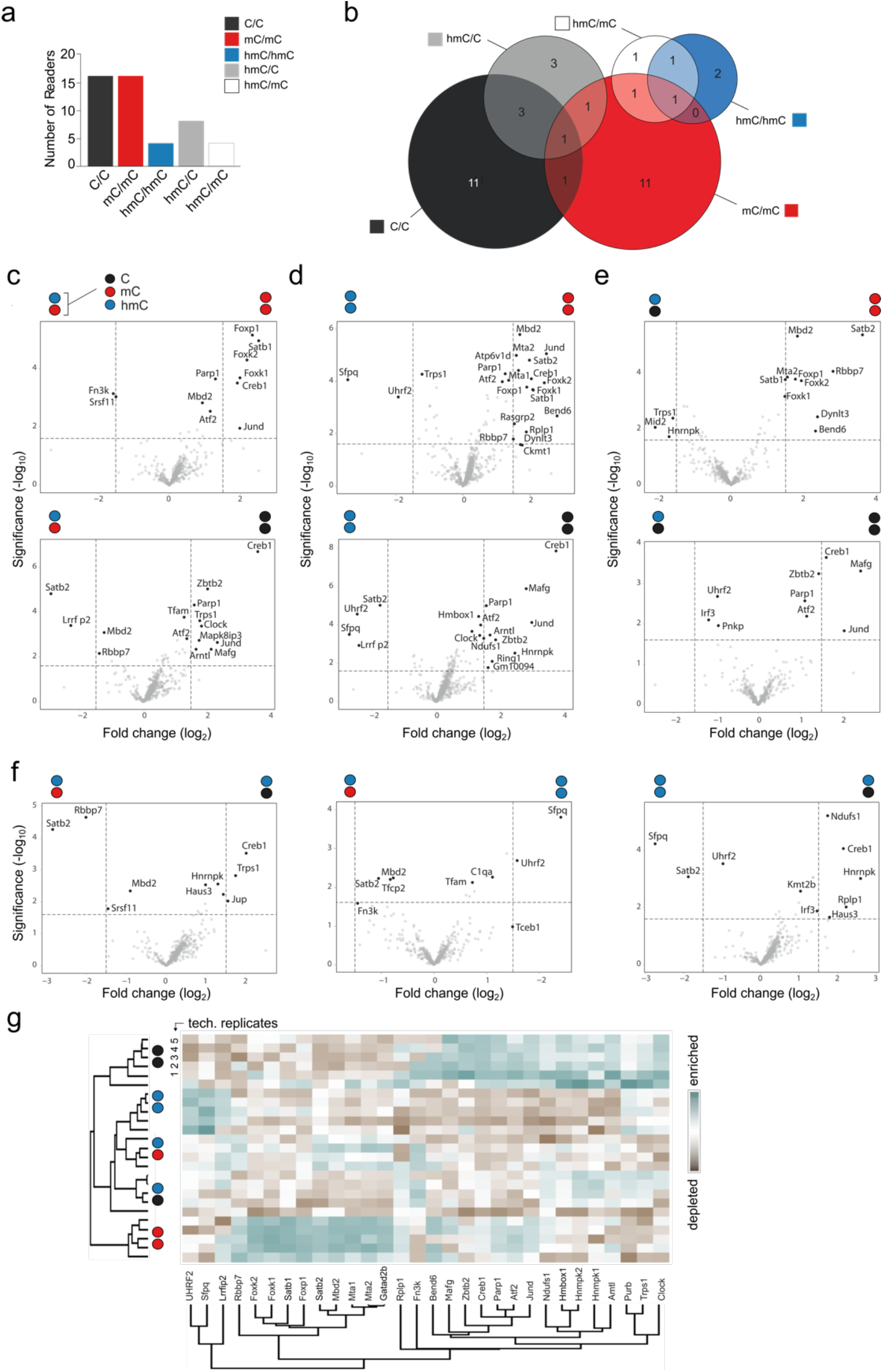
Discovery of readers and antireaders of symmetrically and asymmetrically hmC-modified DNA from mouse brain tissue by enrichment/proteomics. (a) Overall number of enriched reader proteins for indicated DNA probes. (b) Venn diagrams for significantly enriched proteins for indicated probes (note that color code refers to both DNA strands); see **Table S4** for protein names. (c-f) Volcano plots from pairwise probe comparisons (p-value < 0.025 and log2-fold change >1.5). (g) Heatmap of correlation-based clustering of the label-free quantification (LFQ) intensities after log2 transformation and normalization by row mean subtraction. Proteins included in the clustering significantly bind to at least one of the probes as determined by an ANOVA test.

Overall, this data indicates that readers of symmetric and asymmetric hmC marks are highly organism/tissue-dependent and/or vary greatly for different DNA sequences, and indeedm probe- and tissue-specificity has also been observed in previous enrichment studies^20,24–26^. Given that defined DNA enrichment probes cover only a tiny fraction of sequences actually present in the genome, this suggests that many hmC readers are yet to be discovered, and that the full regulatory potential of hmC in its specific symmetry settings remains to be fully explored.

### Validation of selected reader and antireader candidates

Our proteomics profiles indicate that many nuclear proteins have specific preferences for single or multiple symmetric or - asymmetric DNA probes. In particular, hmC/C, hmC/mC and hmC/hmC probes attracted different reader candidates that showed little overlap, a prerequisite for a model in that the (a)symmetry information of hmC marks enables unique regulatory functions. Besides inherently asymmetric CpH marks, the combinatorial nature of CpG dyads that can be differently modified in both dyad strands bears particularly rich symmetry information (**Fig. 1c**). Moreover, each hmC modified CpG mark is written by regulated enzymatic activities: mC/mC-CpGs are the initial substrates of TET dioxygenases, with hmC/mC-CpGs being the first and hmC/hmC-CpGs being a subsequent oxidation product. hmC/C-CpGs may arise from hmC-containing dyads by passive dilution or repair, or from direct TET oxidation of mC/C^5^. To study, if these different TET-associated CpG dyads would exhibit unique protein interactomes as a basis for potential regulatory functions, we evaluated the profiles of a series human (anti)reader candidates individually.

We cloned ten candidates covering different protein scaffolds and transcription factor classes from thyroid and prostate cDNA libraries as fusion constructs with an MBP tag, and expressed them in *E. coli* (**Fig. S15-16**). We applied different assays for pre-screening and in-depth analysis, including enrichments with our five VEGFA PCR probes using expression lysates followed by analyses by anti-MBP dot or western blots, as well as electromobility shift assays (EMSA) using purified proteins with removed MBP tag. EMSAs were performed either with the five VEGFA PCR probes, or with synthetic 57mer variants covering a central part (nt 83-139) of the VEGFA probe and being modified only at four CpGs. These combined experiments report direct DNA binding and modification selectivity of recombinantly expressed readers and complement our enrichment/proteomics results with mammalian nuclear extracts. It should however be noted that the DNA binding properties of a protein may depend on PTMs, interacting proteins, RNAs or ligands only present in mammalian nuclear lysates. Hence, assays with recombinant proteins complement the proteomics data, but do not reliably invalidate candidate readers if negative. In dot blot pre-tests, we identified only few candidates with low DNA binding affinity, indicating either non-functional folding/incompatibility with our assay or indeed indirect recruitment or dependence on other nuclear factors. This included the transcriptional coactivator SUB1^42^, the transcription factor TFAM and the histone methyl-lysine reader L3MBTL3 (**Fig. S17**). However, other readers showed DNA binding and/or interesting selectivities between the five probes in dot blots or initial EMSAs (such as the transcription regulators FOXA1, MYC, MAX, ATF2, RFX5, as well as other chromatin factors such as CHAF1B, **Fig. S17-18**). We studied a selection of candidate readers in more detail.

MYC and MAX, both known to bind nonmethylated DNA^43^, were enriched for C/C over mC/mC and hmC/mC probes, but not for C/C over hmC/C and hmC/hmC (**Fig. 3c-f** and **Table S2 and Fig. S10**). Dot blots indeed showed a trend for selective binding of C/C over mC/mC probes together with an unexpected binding of hmC/hmC for both proteins (**Fig. S18**), which we confirmed by dot and western blots with purified N-terminal MBP fusion proteins in the presence of nuclear extracts (**Fig. 6a** and **S19**). MYC and MAX are basic/helix-loop-helix/leucine zipper (bHLHLZ) transcription factors that typically act as heterodimers to globally amplify the transcription of active genes^44^ via binding to the E-box (CACGTG) and diverse sequence variants containing a central CpG or CpH (**Fig. 6b-c)**. We expressed full-length (fl) and bHLHLZ domain-only versions of MYC and MAX (**Fig. 6b** and **S16c-e**), established functional dimerization protocols (**Fig. S20**) and conducted EMSA assays using our five modified VEGFA probes and MAX_fl_/MYC_bHLHZ_ heterodimers as well as MYC and MAX homodimers (**Fig. S21**). All three dimers showed high affinity to hmC/hmC probes, establishing them as hmC readers (**Fig. 6d**; see **Fig. 6e** for EMSA titration experiments). In addition to MYC, MAX can also dimerize with other bHLHLZ partners (MAD, MNT, MGA and others) to cause different regulatory outputs, and crystal structures of different bHLHLZ dimers with E-box DNA show virtually identical DNA recognition modes with a conserved arginine h-bonding to the CpG guanosine (**Fig. 6b** and **Fig. S22-23**)^45–47^. We modelled the interactions of MYC/MAX with modified CpGs, suggesting that mC abrogates this interaction, whereas the hmC hydroxyl can compensate for this via an alternative h-bond (**Fig. 6f**, a related mode has been described for MAX_2_ with the rare nucleobase caC^43^). This combined data suggests that the observed hmC affinity could be a general property also of other bHLHLZ dimers, a central class of transcription factors that plays essential roles in cell (de)differentiation and cancer progression – processes involving significant changes of genomic hmC levels^44^.

**Figure 6.**
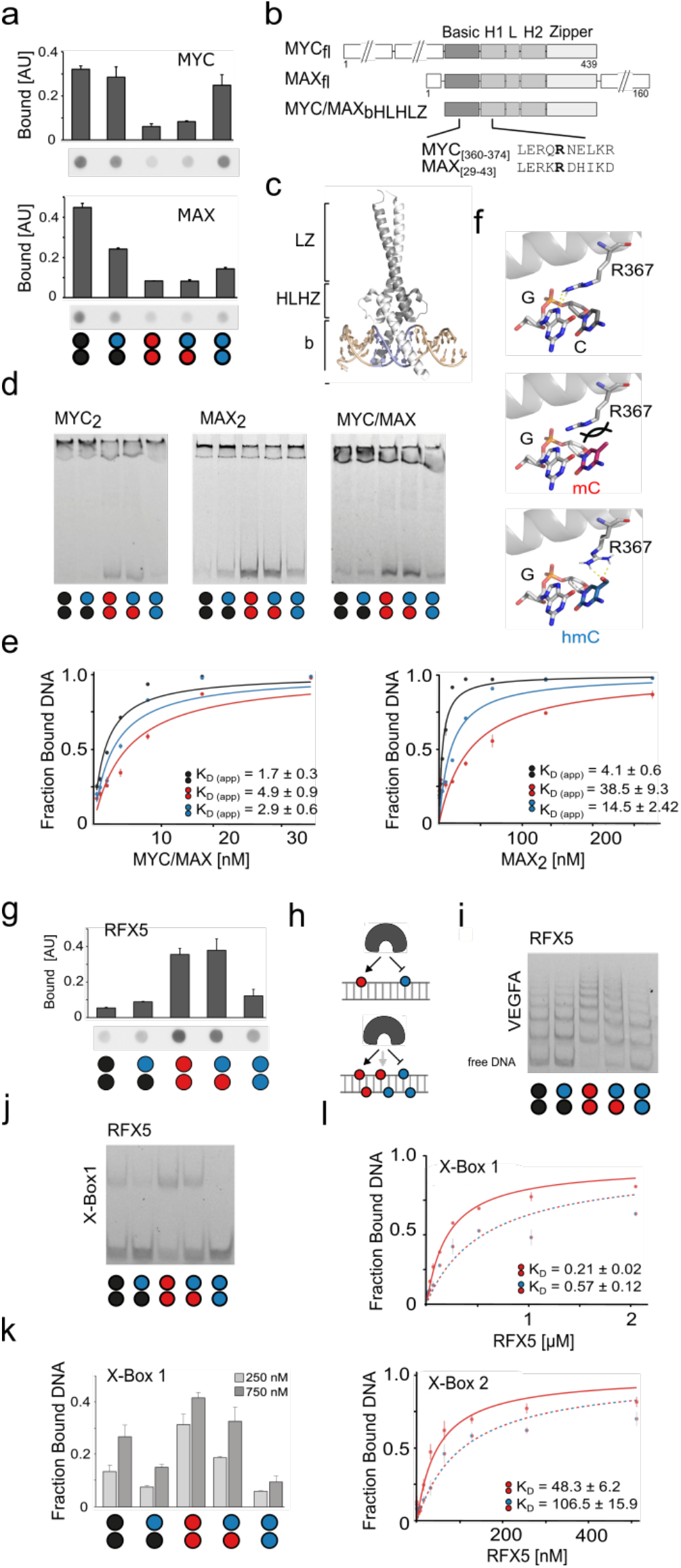
Validation of selected candidate (anti)readers of hmC symmetry. (a) Anti-MBP dot blots of enrichments using VEGFA PCR probes and *E. coli* expression extracts of MYC and MAX proteins fused to an N-terminal MBP tag. (b) Domain structure of full length and bHLHZ truncation of MYC and MAX as well as sequence alignment of DNA-binding bearing conserved arginine R367 (MYC). (c) Crystal structure of the MYC/MAX heterodimer (PDB 1NKP) bound to E-box DNA. Grey: MAX, white: MYC, blue: E-Box. (d) EMSA of MYC__fl_, MAX__bHLHLZ_ homodimers and the respective heterodimer using VEGFA PCR probes modified as shown. (e) EMSA titration experiments and apparent K_D_s with indicated dimers and VEGFA PCR probes modified as shown. (f) Interaction of MYC R367 in the MYC/MAX dimer with E-box G4 (top, PDB 1NKP) and corresponding models of MYC/MAX bound to mC/mC or hmC/hmC dyad, respectively (middle and right). (g) Anti-MBP dot blots of enrichments using VEGFA PCR probes and *E. coli* expression extracts of RFX5 protein fused to an N-terminal MBP tag. (h) Cartoon showing previous (top) and current (bottom) model of RFX5-DNA interaction. (i) EMSA of RFX5 and VEGFA PCR probes modified as shown. (j) EMSA of RFX5 and synthetic X-box 1 probe modified as shown. (k) EMSA profile of RFX5 (two concentrations) and synthetic X-box 1 probe modified as shown. Error bars from duplicate experiments. (l) EMSA titration experiments and K_D_s for RFX5 and synthetic X-Box1 and 2 probes modified as shown.

Given the breadth of their CpG and CpH-containing DNA target sequences and the complexity of the employed VEGFA probe, it is not yet clear, which sequences and dinucleotides may be read by MYC and MAX. JASPAR analysis predicts multiple alternative MYC/MAX target sequences in our probe (**Table S3**), and EMSA-based mapping for MBP-MAX_2_ binding sites in unmodified tiling probes revealed binding to at least three regions of the VEGFA probe (**Fig. S24**). Additional EMSA experiments with synthetic DNA probes containing the E-box (not contained in our VEGFA probe) revealed a C/C > hmC/C > mC/mC, hmC/mC, hmC/hmC CpG dyad preference (**Fig. S20f-g**), whereas for a synthetic VEGFA fragment probe modified only at CpG dyads, we observed a weak hmC/mC > C/C > hmC/C, mC/mC hmC/hmC selectivity (**Fig. S21e**). This data indicates a complex, sequence-dependent hmC-reading of MYC and MAX that we now start to explore by comprehensive in vitro profiling studies using hmC-SELEX and -array assays with a larger number of bHLHLZ dimers as well as ChIP-Seq studies.

We also studied RFX proteins, of which several members showed particular probe preferences in proteomics. These proteins share the conserved DNA-binding winged-helix (WH) domain^48^, and we observed in particular an enrichment of the transcription factor RFX5 and its binding partners RFXANK and RFXAP that together form the so-called RFX complex being involved in MHC-II transcription^49^, with RFX5 mutations being disease-causing for primary MHC class II deficiency^50^. Dot blot analyses revealed a high affinity for the mC/mC and a low affinity for the hmC/hmC probe, respectively, which is in agreement with previous data covering symmetric mC/mC and hmC/hmC dyads (**Fig. 6g**; RFXANK showed no DNA binding, in agreement with an indirect enrichment via its known partner RFX5, **Fig. S17**). This selectivity of RFX5 has been hypothesized to reside on an mC binding pocket that cannot accommodate hmC^24^, observed in a homology model built from a DNA-complex structure of RFX1 that recognizes only one DNA strand^51^. Our extended dataset covering asymmetric probes revealed a similar affinity for the symmetric mC/mC and the asymmetric hmC/mC probe, whereas other probes were bound far less (**Fig. 6g**). This is indeed in agreement with an mC-selective RFX5-DNA interaction that occurs via only one DNA strand (**Fig. 6h**). We studied this interaction in more detail by EMSA using our VEGFA PCR probe and the purified RFX5 DNA binding domain (DBD). RFX5 bound to about six sites in the probe, and as expected from dot blots, with highest affinity to the mC/mC probe (**Fig. 6i**). However, unexpectedly, we observed a lower affinity to the hmC/mC probe, contradicting the aforementioned pocket model. EMSA assays with synthetic DNA probes based on the RFX5 canonical binding motif (the X-box containing only a single RFX5 binding site with a defined CpG modification) afforded the same result, indicating that RFX5 does not recognize the mC nucleobase alone, but is indeed a symmetry-specific reader (with specific preferences among hmC CpG dyads, **Fig. 6j-k**). K_D_ measurements using EMSA with this probe design showed that RFX5 discriminates between the two dyads by a factor of ~2.5, and this discrimination was confirmed with a second synthetic probe containing the X-box target sequence in a different physiological context (**Fig. 4l and S25**)^51^. The data reveals a more complex CpG readout by RFX5 than suggested by previous studies involving only symmetric CpG dyads, and biochemically supports the view that symmetric and asymmetric hmC CpG dyads may cause distinct regulatory outputs from the same RFX5 target sequences.

## 3. Conclusion

We have combined a strategy for generating strand-asymmetric DNA enrichment probes with MS-based proteomics for systematically studying the role of symmetry in the reading of hmC-marks by mammalian nuclear proteomes. By employing promoter probes with five symmetric and asymmetric C, mC and hmC strand-modification combinations, we report the first enrichent/proteomics studies employing probes that cover asymmetric hmC CpG dyads, and in particular the dyad hmC/mC that represents the by far most frequent hmC dyad in tissues characterized so far^12,13^. In human and mouse nuclear extracts, we identify a similar number of readers for all hmC-modified probes as for symmetric probes containing the canonical C/C and mC/mC CpGs combined. This suggests, from a biochemical perspective, that hmC modifications are important protein interaction sites in the genome, with implications for their role in human chromatin regulation. Importantly, the reader groups between the three hmC probes differed greatly, with many more proteins interacting with asymmetric hmC/mC and hmC/C probes as compared to hmC/hmC probes. We confirmed for a series of human readers distinct hmC symmetry preferences. For example, we discover the master regulators MYC and MAX that play central roles in cell (de)differentiation and cancer progression to read hmC in a sequence-dependent manner. We also find RFX5, a transcription factor critically involved in primary MHC class II deficiency to selectively read out specific hmC symmetries in CpG dyads. Our findings collectively illustrate that different hmC symmetries – that are written onto genomes by defined and regulated enzymatic processes – have different nuclear interactomes, with potential to cause distinct regulatory outputs in chromatin. Previous proteomics studies using probes containing symmetrically modified CpG dyads have reported a high sequence- and tissue-dependence of reader profiles^20,24–26^.

We identified very different readers in our human and mouse experiments with asymmetric probes, as well. In addition, hmC levels (and thus presumably hmC symmetries) are generally tissue-dependent and significantly affected in hematopoietic cancers via TET2 and IDH1/2 mutations as well as altered metabolite concentrations affecting TET activity (2-oxoglutarate, 2-hydroxyglutarate, fumarate)^52^, making hmC an important cancer biomarker^53^. Our datasets thus provide a basis for other researchers to study the molecular mechanisms triggered by symmetric and asymmetric hmC modifications in chromatin regulation associated with development and disease.

## Methods

### Cell culture

HEK293T cells were cultivated in DMEM (Dulbeccos’s Modified Eagle Medium) supplemented with 10 % FBS, 2 mM L-glutamine, 100 U/ml penicillin, and 0.1 mg ml^-1^ streptomycin in a sterile humidified incubator (≥ 95 %) at 37 °C and 5 % CO_2_.

### Nuclear extraction

HEK293T cells were washed with 1× PBS solution and detached with trypsin for 5 min at 37°C. Cells were harvested by centrifugation at 300 × g for 5 min. The pellet was washed twice in ice-cold 1× PBS and resuspended in 7 volumes of ice-cold cell lysis buffer (10 mM HEPES pH 7.9, 10 mM KCl, 1.5 mM MgCl_2_, 0.1 mM EDTA, 0.5 mM DTT and 0.5 mM PMSF) containing protease inhibitor cocktail (Roche). After incubation on ice for 10 min, 0.5 % IGEPAL was added, the solution was vortexed and pipetted 50 times with a 200 µl tip. Afterwards centrifugation was carried out at 3000 × g for 20 min at 4°C. The pellet was washed twice with CLB. The washed pellet containing the nuclei was resuspended in nuclear extraction buffer (20 mM HEPES pH 7.9, 420 mM NaCl, 1.5 mM MgCl_2_, 0.2 mM EDTA, 0.5 mM DTT, 0.5 mM PMSF) containing protease inhibitor (Roche) corresponding to half of the cell lysis buffer volume. The solution was incubated at 4 °C and vortexed every 10 min for an overall duration of 50 min. Centrifugation was carried out at 16,000 x g for 20 min and the supernatant was aliquoted and stored at −80 °C. Protein concentrations were determined by BCA assay (ThermoFisher).

### Nuclear extraction from tissue

Mouse brain was cut on dry ice and rinsed in PBS. Cut tissue was added to a dounce homogenizer along with buffer A (10 mM HEPES pH 7.9, 10 mM KCl, 0.1 mM EDTA, 0.1 mM EGTA, 1mM DTT, 0.5 mM PMSF) containing protease inhibitor cocktail (Roche) (10 µl buffer A/mg tissue). The tissue was homogenized by 15 strokes with pestle A, followed by addition of 0.5 % IGEPAL and another 15 strokes with pestle A. After incubation on ice for 10 min, the homogenate was centrifuged at 3000 x g for 10 min at 4 °C. The pellet was washed twice with buffer A. The washed pellet containing the nuclei was resuspended in buffer B (20 mM HEPES pH 7.9, 400 mM NaCl, 1 mM EDTA, 1 mM EGTA, 1 mM DTT, 1 mM PMSF) containing protease inhibitor (Roche) corresponding to half of the buffer A volume. The solution was incubated at 4 °C and vortexed every 10 min for an overall duration of 50 min. Centrifugation was carried out at 16,000 x g for 20 min and the supernatant was aliquoted and stored at −80 °C. Protein concentrations were determined by BCA assay.

### DNA probe generation

The VEGFA probe (o5488, Table S5) was amplified via PCR using Dreamtaq DNA Polymerase (Thermo Fisher Scientific) with the biotinylated forward primer o4681 and the reverse primer o4666 (or Cy5 labeled o4746). 16 pmol of PCR product was immobilized on 20 µl Dynabeads™ MyOne™ Streptavidin T1 (ThermoFisher) in 1x Binding and washing (B & W) buffer (5 mM Tris-HCl pH 7.5, 0.5 mM EDTA, 2 M NaCl) on a turning wheel for 3 h at RT. The beads were washed twice with 1xB&W buffer and then resuspended in 50 µl 20 mM NaOH. After incubation on a turning wheel for 10 min at RT, the supernatant containing single-stranded DNA (ssDNA) was separated from the beads with a magnet and then added to a new tube containing a final concentration of 1xTE (10 mM Tris-HCl pH 7.5, 1 mM EDTA) and 20 mM HCl. The ssDNA was hybridized with 1.5 x molar excess of o4681 (or Cy3-labeled o6021) in 1x hybridization buffer (20 mM HEPES PH 7.3, 30 mM KCl, 1 mM EDTA, 1 mM (NH4)_2_SO_4_). The annealing mixture was heated to 95 °C and then gradually cooled down to RT in a Dewar filled with boiling water. The primer was extended with Klenow fragment in 1 x NEB2 buffer using 0.1 mM dNTPs including the respective modified dCTP for 3 h at 37 °C. The final product was purified with a PCR clean-up kit (Macherey-Nagel) and analyzed by agarose gel electrophoresis and ethidium bromide staining (or scanned using a Typhoon FLA-9500 laser scanner (GE Healthcare) with excitation/emission wavelengths of 550/570 nm for Cy3 and 649/670 nm for Cy5 for probes generated with labeled primers, see **Table S5 and Fig. S6**). Additionally, FAM-labeled probes were generated using a biotin- and FAM-containing forward primer (o4745) and analyzed by 8.3 M urea on an 8 % sequencing PAGE with 50 cm chamber length (Carl Roth). They were scanned using a Typhoon FLA-9500 laser scanner (GE Healthcare) with excitation/emission wavelengths of 490/520 nm. Unlabeled probes with different cytosine modification combinations were sequenced by Sanger sequencing (see **Fig. S8**). The Sp1 probe (p3546, **Table S5**) was amplified via PCR using Dreamtaq DNA Polymerase (Thermo Fisher Scientific) with the biotinylated forward primer o5232 and the reverse primer o5233. Probe generation was then carried out as described above.

### DNA pulldowns

5 µl of Dynabeads™ MyOne™ Streptavidin T1 (ThermoFisher Scientific, 65601) were incubated with 4 pmol biotinylated DNA probe (or more for NX pulldowns, compare **Fig. S1-4**, the amount of beads was upscaled accordingly) in 1x B&W buffer for 3 h at RT. The beads were then washed once with 1xB&W buffer followed by two washes with Protein binding buffer (PBB) (50 mM Tris-HCl pH 8, 150 mM NaCl, 1 mM DTT, 1mM PMSF, 0.25 % IGEPAL) containing protease inhibitor. They were then incubated with 50 µg nuclear extracts (3.6 µM) and 5 µg dA:dT (1.1 µM) competitor DNA in PBB in a total volume of 300 µl on a turning wheel overnight at 4°C (note that these reducing conditions are not compatible with O_2_-dependent TET activity, ensuring integrity of the probes during incubation). The beads were washed twice with 1xPBS and stored in 20 µl PBS until further processing.

### On-bead digestion

Pulldown samples stored in 20 µl PBS were denatured and reduced on the beads by adding 50 µl Denaturation/Reducing buffer (8 M urea, 50 mM Tris pH 7.5, 1mM DTT) for 30 min at 20 °C and 350 rpm before 5.55 µl alkylation solution (50 mM chloroacetamide in Denaturing/Reducing buffer) was added for 30 min at 20 °C and 350 rpm. Samples were on-bead digested by adding 1 µg Lys-C (NEB) dissolved in nuclease free water for 1 h at 37 °C and 350 rpm. The supernatant was transferred into a new tube. 1 µg Trypsin (NEB) dissolved in 165 µl 50 mM Tris-HCl was added to the beads and incubated for 1 h at 37 °C 350 rpm. The supernatants from both digestions were combined. To the combined supernatants 2 µg Trypsin were added. Incubation was carried out for overnight at 37 °C 350 rpm. The reaction was stopped by adding 20 µl 10 % TFA. The digested peptides were desalted using Pierce™ C18 Tips (ThermoFisher). The tips were activated by aspirating and dispensing 100 µl 50 % ACN in water twice and equilibrated with 100 µl 0.1 % TFA. The sample was aspirated and dispensed for 15 cycles. The tip was rinsed by aspirating 0.1 % TFA/5 % ACN twice. The samples were eluted by aspirating two times 20 µl of 0.1 % formic acid and dispensing into a new tube. The eluted peptides were dried in a vacuum concentrator for 1.5 h at 30 °C.

### Mass spectrometry

Following the Lys-C/tryptic digests, the dried peptides were dissolved in 10 - 20 µl of 0.1% TFA and analyzed by nanoHPLC-MS/MS using an Ultimate 3000 RSLC nano-HPLC system and a Hybrid-Orbitrap mass spectrometer Q Exactive Plus (ThermoFisher). 3 - 5 µl of the peptide solution were injected and enriched on a C18 PepMap 100 column (5 mm, 100 Å, 300 mm ID * 5 mm, Dionex) using 0.1% TFA, at a flow rate of 30 µL/min, for 5 min and separated on a C18 PepMap 100 column (3 mm, 100 Å, 75 mm ID * 50 cm) using a linear gradient (5-30% ACN/H2O + 0.1% formic acid over 90 min) with a flow rate of 300 nL/min. The nano-HPLC apparatus was coupled online using a nano emitter with 10 µm tip diameter. Signals in the mass range of m/z = 300 - 1650 were acquired at a resolution of 70,000 for full scan, followed by up to ten high-energy collision-dissociation (HCD) MS/MS scans of the most intense at least doubly charged ions at a resolution of 17,500. Proteins were relatively quantified by using MaxQuant^54^ v.2.2.0.0, including the Andromeda search algorithm and searching in parallel the Homo sapiens reference proteome of the UniProt database and a contaminants database implemented in MaxQuant. Briefly, an MS/MS ion search was performed for enzymatic trypsin cleavage, allowing two missed cleavages. Carbamidomethylation was set as a fixed protein modification, and oxidation of methionine and acetylation of the N-terminus were set as variable modifications. The mass accuracy was set to 20 parts per million (ppm) for the first search, and to 4.5 ppm for the second search. The false discovery rates for peptide and protein identification were set to 0.01. Only proteins for which at least two peptides were quantified were chosen for further validation. Relative quantification of proteins was performed by using the label-free quantification algorithm implemented in MaxQuant.

Statistical data analysis of pulldown samples was performed using Perseus^55^ v.2.0.7.0 and 2.0.11.0 including proteins which were identified in at least four out of five technical replicates in at least one of the two compared conditions. Label-free quantification (LFQ) intensities were log-transformed (log2); replicate samples were grouped together. Missing values were imputed using small normally distributed values, and a two-sided t-test was performed. Volcano plots were generated using the VolcaNoseR web app^56^. Proteins with log2-fold changes < 1.5 and −log p <0.025 were considered as statistically significant enriched.

### Western blotting

Pulldown samples were eluted in 15 µl PBS and boiled with 5 µl 4x Laemmli sample buffer at 95 °C for 5 min and subsequently run on 12 % SDS Gels. Proteins were transferred to polyvinylidene fluoride (PVDF) membranes using a Trans-Blot Turbo system (BioRad, 1.9 A, 25 V, 7 min). The membranes were blocked for at least 1 h using 5 % (m/v)skimmed milk in TBS-T (TBS + 0.1 % v/v Tween-20). MBP-antibody (NEB E8032) was incubated 1:10,000 in 1.25 % skimmed milk in TBS-T overnight at 4 °C. The secondary antimouse antibody coupled to horseradish peroxidase (Sigma, GENA931) was incubated in 2.5 % skimmed milk in TBS-T for 1 h at RT. The chemiluminescent reaction was initiated using a Clarity Western ECL substrate (Bio-Rad) and images were acquired on a Bio-Rad ChemiDocTM imaging system.

### Recombinant protein expression for DNA interaction assays

The coding sequences of MYC (aa1-439, [AAA36340.1]), MAX (aa1-160, [NP_001394023.1]), RFX5 (aa1-616, [NP_000440.1]), RFXANK (aa1-260, [NP_001357162.1]), L3MBTL3 (aa1-780, [NP_115814.1]), SUB1 (aa1-127, [NP_006704.3]), FOXA1 (aa1-472, [NP_004487.2]), ATF1 (aa1-505, [NP_001243019.1]), TFAM (aa1-246, [NP_003192.1]), and CHAF1b (aa1-559, [NP_005432.1]) were amplified from thyroid (HD-503, Zyagen) or prostate (10108-A, Biocat) cDNA libraries (see Table S5 for primer sequences) and cloned into an *XhoI*-digested pET-21d(+) vector containing a N-terminal MBP tag and a C-terminal His tag by Gibson assembly (Fig. S4a). Expression plasmids were transformed into *E. coli* BL21-Gold(DE3) (Agilent). Fresh overnight cultures of single clones were then diluted to an optical density (OD600) of 0.05 in 30 ml LB-Miller broth supplemented with 50 µg/mL carbenicillin, 1 mM MgCl_2_ and 1 mM ZnSO_4_ and grown at 37 °C until they reached an OD600 of 0.5 – 0.6. Expression was induced with 1 mM IPTG and cultures were incubated overnight at 30 °C shaking at 150 rpm. Cells were harvested and washed once by resuspension in ice-cold 20 mM Tris-HCl (pH = 8.0) before they were lysed by sonication (Branson Digital Sonifier 450 Cell Disruptor, 3 min, amplitude 15 %, 4 s on, 2 s off) in 2 ml lysis buffer (50 mM Tris-HCl pH 8.0, 20 % sucrose, 1 mM EDTA, 0.5 mM PMSF, 1 mM DTT, 0.1 mg/ml lysozyme, 0.1 % Triton X-100). After centrifugation at 14,000 × g for 20 min at 4 °C, the cleared lysates were retained, snap-frozen in liquid nitrogen and stored at −80°C. Enrichments were performed using 1.67 µl of Dynabeads™ MyOne™ Streptavidin T1 (ThermoFisher) to immobilize 1.33 pmol of VEGFA DNA probe which was subsequently incubated with 0.01 or 0.1 µl of *E. coli* lysates in the presence of 1.67 µg dA:dT competitor DNA. Pulldown samples were boiled in 15 µl PBS and 5 µl 4x SDS sample buffer (250 mM Tris-HCl pH 6.8, 500 mM DTT and 0.1 % SDS) at 95 °C for 5 min. 10 µl of these samples were mixed with 40 µl PBS and then applied to a nitrocellulose membrane (Amersham™ Protran®) using gravity filtration in a Bio-Dot® apparatus (Bio-Rad). The wells were rinsed twice with TBS-T using vacuum until dry. The blotted membrane was taken out from the apparatus and blocked for at least 1 h using 5 % (m/v)-skimmed milk in TBS-T. Afterwards the membrane was treated in the same way as described for western blots.

### Cloning and purification of recombinant proteins for electrophoretic mobility shift assays

Full-length (fl) MAX was obtained by removing MBP from MAX (aa1-160) using inverse PCR with primers o5938 and o5939. The basic helix-loop-helix and Leucine zipper domain (bHLHZ) of MYC (aa 354-434) was cloned by first removing the N-terminal part of MYC (aa1-439) with primers o5980 and o5981, followed by removal of the C-terminal part using the primers o5983 and o5984. For the MBP fused RFX5 DNA-binding domain (DBD) (aa75-168), the C-terminal part of RFX5 (aa1-616) was first removed using primers o6035 and o6036 and the N-terminal part was subsequently removed using the primers o6037 and o6038. In each case, ligation was carried out with T4 ligase at 16 °C overnight (see **Table S6** for primer sequences). Expression was carried out as described above. For purification of MAX fl, RFX5 DBD, MBP-PCGF6 and MBP-RNF2, harvested cells were resuspended in 2 ml binding buffer (20 mM Tris-HCl, 250 mM NaCl, 10% glycerol, adjusted to pH = 8.0, supplemented with 10 mM 2-mercaptoethanol, 5 mM imidazole and 0.1% Triton X-100) before 1 mM PMSF was added. Resuspended cells were sonicated (Branson Digital Sonifier 450 Cell Disruptor, 3 min, amplitude 15 %, 4 s on, 2 s off) and subsequently incubated with 1 mg/ml lysozyme and 10 U DNAseI overnight at 4 °C. After centrifugation at 14,000 × g for 20 min at 4 °C, the cleared supernatants were retained and incubated at 16 °C for 20 min shaking at 700 rpm with 450 µL 50% Ni-nitriloacetic acid (NTA) HisPur agarose resin (ThermoFisher) that were prior equilibrated in binding buffer. The resins were washed 2 x with 1 mL binding buffer containing 90 mM imidazole for 5 min at 16 °C shaking at 700 rpm and the fusion protein was eluted in 2 × 0.2 mL and 1 × 0.4 mL binding buffer with 500 mM imidazole for 5 min at 16 °C shaking at 700 rpm. Fractions with sufficient purity judged by SDS PAGE were combined and dialyzed against 3 × 1 L 20 mM HEPES, 100 mM NaCl, 10% glycerol, adjusted to pH = 7.3, and 0.1% Triton X-100 in a tube with 12-14 kDa MWCO (Roth). For purification of MYC bHLHZ, the cell pellet was resuspended in 2 ml urea buffer (PBS with 1 % Triton X-100, 1 mM PMSF, 5 mM imidazole and 7 M urea) similarly as described previously^47^. Cell lysis was achieved by sonication (Branson Digital Sonifier 450 Cell Disruptor, 3 min, amplitude 15 %, 4 s on, 2 s off) prior to centrifugation for 20 min at 16,000 g at 4 °C. The supernatant was adjusted to a final concentration of 750 mM NaCl. The adjusted supernatant was incubated at 4 °C for 1 h shaking at 700 rpm with 450 µL 50% Ni-nitriloacetic acid (NTA) HisPur agarose resin (ThermoFisher) that were prior equilibrated in urea buffer. The resins were washed with 5 mL wash buffer (10 mM Tris pH 7.8, 500 mM NaCl, 1 % Triton X-100, 1 mM PMSF, 10 mM imidazole and 7 M urea). A second wash step was carried out using the same buffer with 100 mM NaCl. The protein was eluted by stepwise elution with wash buffer (5x 1 ml) with 100 mM NaCl and 250 mM imidazole. Fractions with sufficient purity judged by SDS PAGE were combined and dialyzed against 3 × 500 ml 10 mM Tris HCl pH 7.8, 100 mM NaCl, 1 % Triton X-100, 0.1 mM PMSF and 7 M urea in a tube with 12-14 kDa MWCO (Roth). The protein concentration was determined with a BCA assay (ThermoFisher) and the proteins were snap-frozen in liquid nitrogen and stored at −80 °C.

### Electrophoretic mobility shift assay

Differentially modified FAM labeled DNA probes (**Table S5**) were obtained either by asymmetric probe generation as described above using a FAM labeled forward primer and unlabeled reverse primer (**Table S5**) or by hybridization of 1.5 µM labeled forward strand and

1.8 µM unlabeled reverse strand oligos (**Table S5**) in IDT buffer (30 mM HEPES, 100 mM KOAc) by heating to 95 °C and then gradually cooling to RT in a Dewar flask filled with boiling water. MBP-fusion proteins were treated with 0.25 µM TEV-protease overnight at 4 °C to remove the MBP tag. For cytosine modification binding studies, varying concentrations of protein were incubated with 2 nM labeled DNA probe carrying the respective modification in EMSA buffer (20 mM HEPES, 30 mM KCl, 1 mM EDTA, 1 mM (NH4)2SO4, 0.2 % Tween 20) in presence of 5 µM dA:dT competitor and 1 mM DTT in a total volume of 15 µl for 20 min at 21 °C. For binding affinity studies, a dilution series of protein was prepared, ranging from 0.5 – 256 nM for MAX fl, 0.5 – 32 nM for MAX fl/ MYC bHLHZ dimer and 1 – 2048 nM for RFX5 DBD. Proteins were incubated with 2 nM labeled DNA probe under the conditions described above. Heterodimerization of MAX fl and MYC bHLHZ was achieved by incubating the proteins in a 1:300 MAX fl:MYC bHLHZ ratio for 30 min at 21 °C before adding the putative dimer to the DNA probe. Samples were mixed with 3 µl 6x EMSA loading buffer (1.5 x TBE, 40 % glycerol) and loaded to a non-denaturing polyacrylamide gel (12 % for VEGFA probes, 15 % for E-box and X-box probes) that was pre-run at 200 V for 2 h. Loaded gels were run at 240 V at 4 °C for 60 - 75 min, depending on DNA probe length. Visualization was performed using the 510 LP filter of the Typhoon FLA-9500 laser scanner (GE Healthcare) at 473 nm with 800 V-1000 V amplification. Bound and unbound DNA probes were quantified using ImageQuant TL v8.1 1D Gel Analysis (GE Healthcare).

### Binding affinity analysis

Band intensities from EMSA assays were quantified as described above. The bound fraction of DNA was calculated for each protein concentration and analyzed with R v4.1.2. Two independent replicates were averaged and the standard error of the mean (SEM) was determined. To estimate the equilibrium dissociation constant (kD), a one-site binding model was applied to the data. This model assumes 1:1 interaction of protein and DNA, which is described by the equation f=[P]/(K_D_+[P]), where f is the bound DNA fraction and [P] is the protein concentration. Nonlinear least-squares regression was used to fit the model to the data. K_D_ estimates and their standard errors were obtained from the regression. The goodness of the fit was evaluated using R^2^ and the residual standard error (RSE). Binding curves were visualized using the ggplot2 package.

### Modelling

Modification of the canonical bases to either mC or hmC was done via the CHARMM-GUI web server^57^ in order to generate topology and input files for energy minimization of the resulting structures in GROMACS^58^. Modelling of the newly formed interactions with hmC was pursued through manual iteration of the dihedral angles of rotatable bonds, c_1_ to c_4_, in the arginine side chain under careful consideration of spatial restraints within the major groove. Figures were composed in PyMol^59^.

## ACKNOWLEDGMENT

We thank the Faculty of Chemistry and Chemical Biology of the TU Dortmund University and the International Max-Planck Research School for Living Matter for continuous support. We thank the members of the group for support. We thank Dr. Cristina Cadenas for mouse brain. Funded by the Deutsche Forschungsgemeinschaft (SU726/9-1 and SU726/10-1) and the CANTAR program “Netzwerke 2021”, an initiative of the Ministry of Culture and Science of the State of Northrhine Westphalia. Funded/Co-funded by the European Union (ERC, Grant 101082494, bypassNMR). Views and opinions expressed are those of the author(s) only and do not necessarily reflect those of the European Union or the European Research Council. Neither the European Union nor the granting authority can be held responsible for them.

## AUTHOR CONTRIBUTIONS

L.E. developed methods, conducted wet lab experiments, and analyzed data. Z.C. developed methods, conducted wet lab experiments, and analyzed data. S.E. cloned plasmids and expressed and purified proteins. T.G. and R.L. conducted modelling experiments. P.J. supervised MS experiments and analyzed MS data. D.S and L.E. wrote the manuscript with the help of all other authors. D.S. analyzed data and conceived and supervised the study.

## ADDITIONAL INFORMATION

Associated content 1: Table S1 (Excel file); 2: Table S2 (Excel file); 3: Table S3 (Excel file), 4: Tables S4-S7 and, Figures S1-S45 (PDF).

