## Supplementary information for "Mammalian Proteome Profiling Reveals Readers and Antireaders of Strand-Symmetric and -Asymmetric 5-Hydroxymethylcytosine-Modifications in DNA"

This PDF file includes:
Supplementary Table 4-7
Supplementary Fig. 1 to 25
Uncropped Gel Fig. S26-S45

**Table S4**

**a.** Significantly enriched proteins (p-value < 0.025, FC > 1.5) for C/C, hmC/C, mC/mC, hmC/mC and hmC/hmC modifications in at least one out of two biological experiments (experiments 1 and 2) associated with the Venn diagram **Fig. 3b**.

| CpG dyad | # proteins | Protein names |
|---|---|---|
| C/C, hmC/C, hmC/mC, mC/mC | 2 | RFX5, SSBP2 |
| hmC/C, hmC/hmC, hmC/mC, mC/mC | 2 | UHRF1, MECP2 |
| C/C, hmC/C, mC/mC | 4 | ATF1, BANP, LDB1, CREB1 |
| C/C, hmC/C, hmC/mC | 1 | TFAP2A |
| C/C, hmC/mC, mC/mC | 1 | RFX1 |
| hmC/C, hmC/mC, mC/mC | 1 | SIX1 |
| hmC/hmC, hmC/mC, mC/mC | 1 | FOXC1 |
| C/C, hmC/C | 24 | ARNT, ASH2L, BEND3, TFPT, NKX2-5, ZIC1, MYC, BCOR, MAX, RBBP5, TFAM, CENPB, NKX2-1, SAMD1, PCGF1, ZIC2, INO80C, NKX2-3, RBM45, L3MBTL3, KDM2B, AHR, RUNX2, FLYWCH1 |
| C/C, mC/mC | 3 | PCGF6, E2F6, MGA |
| hmC/C, hmC/hmC | 2 | EXOSC5, SF3A3 |
| hmC/mC, mC/mC | 8 | SUB1, FOXF1, MBD4, RFXANK, ZFHX3, PTBP3, FOXA1, RFXAP |
| hmC/hmC, mC/mC | 1 | DACH1 |
| hmC/hmC, hmC/mC | 3 | CELSR1, KRR1, SYF2 |
| C/C | 31 | SMAD1, DPY30, MTF2, WDR5, KMT2B, VEZF1, ZMYND11, MEN1, ZBTB2, CBFB, NFIC, ESRRA, HCFC2, KDM2A, TBRG1, EHMT1, CTCF, MSANTD3, RUVBL1, ZBTB10, MAZ, NFIX, KMT2A, ACTR5, SKP1, HEY1, PBX1, RING1, TFAP4, TIMM8A, RNF2 |
| hmC/C | 11 | CDCA7L, CCAR2, CSTA, SRP68, PRPF4B, AIFM1, SYNCRIP, PAX1, TERF2IP, PHF3, MAPRE2 |
| mC/mC | 21 | DEK, TBX21, CFL1, BUB3, PPIA, ZFHX2, MTA2, ATF2, MDH2, CBX3, RFX7, TBX2, RBBP7, L3MBTL2, MBD2, SUMO2, HMGB2, EIF5A, ZBTB44, CDX1, CREB5 |
| hmC/mC | 3 | ZC3HAV1, BAG2, NOP16 |
| hmC/hmC | 11 | CHAF1A, SRFBP1, DNAJB3, RADX, CHAF1B, HNRNPLL, BAZ1B, RPP25L, KLHL7, RBM14, TOX |

**b.** Significantly enriched proteins (p-value < 0.025, FC > 1.5) for C/C, hmC/C, mC/mC, hmC/mC and hmC/hmC modifications in the mouse brain proteomics study associated with the Venn diagram **Fig. 5b**.

| CpG dyad | # proteins | Protein names |
|---|---|---|
| C/C, hmC/C, mC/mC | 1 | Creb1 |
| hmC/hmC, hmC/mC, mC/mC | 1 | Satb2 |
| C/C, hmC/C | 3 | Mid2, Hnrnpk, Trps1 |
| C/C, mC/mC | 1 | Jund |
| hmC/C, mC/mC | 1 | Rplp1 |
| hmC/mC, mC/mC | 1 | Rbbp7 |
| hmC/hmC, hmC/mC | 1 | Lrrfip2 |
| C/C | 11 | Mtmr10, Arntl, Lurap1, Zbtb2, Baz2b, Clock, Mafg, Gm10094, Parp1, Mapk8ip3, Ring1 |
| hmC/C | 3 | Jup, Ndufs1, Haus3 |
| mC/mC | 11 | Atp6v1d, Dynlt3, Mta2, Foxp1, Bend6, Rasgrp2, Mbd2, Gatad2b, Satb1, Foxk1, Foxk2 |

| | | |
|---|---|---|
| hmC/mC | 1 | Fn3k |
| hmC/hmC | 2 | Sfpq, Uhrf2 |

**c.** Significantly enriched proteins (p-value < 0.025, FC > 1.5) for C/C, hmC/C, mC/mC, hmC/mC and hmC/hmC modifications in experiment 1 associated with the Venn diagram **Fig. S14 (left)**.

| CpG dyad | # proteins | Protein names |
|---|---|---|
| hmC/C, hmC/hmC, hmC/mC, mC/mC | 1 | UHRF1 |
| C/C, hmC/C, mC/mC | 2 | ATF1, LDB1 |
| C/C, hmC/C, hmC/mC | 1 | TFAP2A |
| hmC/C, hmC/mC, mC/mC | 1 | SIX1 |
| hmC/hmC, hmC/mC, mC/mC | 1 | FOXC1 |
| C/C, hmC/C | 15 | SAMD1, ASH2L, BEND3, ZIC2, BANP, NKX2-5, RBM45, L3MBTL3, ZIC1, MYC, BCOR, MAX, RBBP5, FLYWCH1, TFAM |
| C/C, mC/mC | 2 | E2F6, PCGF6 |
| hmC/C, hmC/hmC | 2 | EXOSC5, SF3A3 |
| hmC/mC, mC/mC | 6 | MBD4, RFXANK, SUB1, RFX5, FOXA1, RFXAP |
| hmC/hmC, mC/mC | 1 | DACH1 |
| hmC/hmC, hmC/mC | 2 | KRR1, CELSR1 |
| C/C | 24 | ARNT, CENPB, DPY30, MTF2, PCGF1, EHMT1, CTCF, WDR5, MSANTD3, KMT2B, RUVBL1, MAZ, NFIX, ZMYND11, SSBP2, MEN1, ZBTB2, KMT2A, SKP1, KDM2B, AHR, ESRRA, HCFC2, RNF2 |
| hmC/C | 8 | CSTA, SRP68, PRPF4B, AIFM1, PHF3, SYNCRIP, MAPRE2, PAX1 |
| mC/mC | 10 | DEK, TBX21, ZFHX2, MTA2, CBX3, RFX7, TBX2, MAG, CDX1, L3MBTL2 |
| hmC/mC | 2 | ZC3HAV1, NOP16 |
| hmC/hmC | 6 | BAZ1B, SRFBP1, KLHL7, RBM14, DNAJB3, RADX |

**d.** Significantly enriched proteins (p-value < 0.025, FC > 1.5) for C/C, hmC/C, mC/mC, hmC/mC and hmC/hmC modifications in experiment 2 associated with the Venn diagram **Fig. S14 (middle).**

| CpG dyad | # proteins | Protein names |
|---|---|---|
| C/C, hmC/C, hmC/mC, mC/mC | 1 | RFX5 |
| hmC/C, hmC/hmC, hmC/mC, mC/mC | 2 | MECP2, UHRF1 |
| C/C, hmC/C, mC/mC | 2 | BANP, CREB1 |
| C/C, hmC/mC, mC/mC | 1 | RFX1 |
| hmC/C, hmC/mC, mC/mC | 1 | SSBP2 |
| C/C, hmC/C | 16 | ARNT, CENPB, NKX2-1, SAMD1, PCGF1, BEND3, INO80C, NKX2-3, TFPT, L3MBTL3, KDM2B, MYC, BCOR, AHR, RUNX2, MAX |
| hmC/mC, mC/mC | 8 | FOXF1, FOXC1, RFXANK, ZFHX3, PTBP3, SUB1, FOXA1, RFXAP |
| hmC/hmC, hmC/mC | 1 | SYF2 |
| C/C | 27 | SMAD1, DPY30, TBRG1, ASH2L, ZIC2, VEZF1, ZBTB10, NFIX, MEN1, ZIC1, ZBTB2, CBFB, KMT2A, ACTR5, HEY1, NFIC, PBX1, MAG, RING1, TFAP4, ESRRA, RBBP5, HCFC2, FLYWCH1, TFAM, TIMM8A, KDM2A |
| hmC/C | 4 | CDCA7L, CCAR2, TERF2IP, RBM45 |
| mC/mC | 14 | CFL1, MBD2, BUB3, SUMO2, PPIA, MTA2, ATF2, MDH2, HMGB2, EIF5A, RFX7, ZBTB44, CREB5, RBBP7 |

| | | |
|---|---|---|
| hmC/mC | 1 | BAG2 |
| hmC/hmC | 6 | CHAF1B, HNRNPLL, CHAF1A, RPP25L, DACH1, TOX |

**e.** Significantly enriched proteins (p-value < 0.025, FC > 1.5) for C/C, hmC/C, mC/mC, hmC/mC and hmC/hmC modifications in experiment 3 associated with the Venn diagram **Fig. S14 (right)**.

| CpG dyad | # proteins | Protein names |
|---|---|---|
| C/C, hmC/mC | 5 | ZIC2, BEND7, MYC, MAX, C14orf93 |
| hmC/mC, mC/mC | 16 | KHDRBS2, FOXF1, FOXC1, YLPM1, MECP2, NCOA5, MBD4, RFXANK, ZFHX3, RBM12B, UHRF1, RFX5, RFX7, SAFB2, SAFB, RFXAP |
| C/C | 67 | ARNT, CENPB, TCF12, DPY30, SAMD1, CCAR2, MXD4, MTF2, TERF1, PCGF1, CTCF, SUZ12, ASH2L, GLI1, BEND3, INO80C, KMT2B, VEZF1, EED, DNMT1, BANP, ZNF639, ZBTB10, MAZ, NFIX, TFAP2C, NKX2-5, MITF, RUNX1, ATF3, RBM45, ZMYND11, DYNLL1, L3MBTL3, ZIC1, CXXC5, ESRRG, ZBTB2, KMT2A, ZBTB25, CEBPB, ZNF148, SKP1, NFIC, XBP1, KDM2B, BCOR, ESRRA, AHR, TFAP4, MXI1, RUNX2, RBBP5, HCFC2, FLYWCH1, TFAM, PATZ1, S100A9, EZH2, ZNF740, NFIA, BCL11A, E2F2, KDM2A, KLF12, CREB1, HAND1 |
| mC/mC | 17 | PRPF40A, ACIN1, ZNF326, MBD2, RBM3, ZFHX2, ATF2, MBD1, SUB1, RBM25, FOXK1, ZBTB44, MTA1, RFX2, GATAD2A, RFX1, KDM6A |
| hmC/mC | 10 | CHAF1B, SFPQ, RBM4, PAX3, TBP, RBM14, RAI14, FBXO11, NONO, ZNF638 |

**Table S5**

Oligonucleotides used for probe generation and protein interaction assays.

| Name | Description | Purpose | Sequence 5' -> 3' |
|---|---|---|---|
| o4681 | VEGFA fw | PCR primer | [BtnTg]TTTCCAAAGCCCATTCCCT |
| o4666 | VEGFA rv | PCR primer | AGTGACCCCTGGCCT |
| o5488 | VEGFA template | Template | TTTCCAAAGCCCATTCCCTCTTTAGCCAGAGCCGGGGTGTGCAGACGGCAGTC ACTAGGGGGCGCTCGGCCACCACAGGGAAGCTGGGTGAATGGAGCGAGCAGCG TCTTCGAGAGTGAGGACGTGTGTGTCTGTGTGGGTGAGTGAGTGTGTGCGTGT GGGGTTGAGGGCGTTGGAGCGGGGAGAAGGCCAGGGGTCACT |
| o4745 | VEGFA FAM fw | PCR primer | [BtnTg]TTTCCAAAGCCCA[FAM]TCCCT |
| o4746 | VEGFA Cy5 rv | PCR primer | [Cy5]AGTGACCCCTGGCCT |
| o6021 | VEGFA Cy3 fw | PCR primer | [Cy3]TTTCCAAAGCCCATTCCCT |
| o5232 | SP1 fw | PCR primer | [BtnTg]CTGGGCGGAAACCAAGTACG |
| o5233 | SP1 rv | PCR primer | GGAAAAACGCGGACGCTGAC |
| p3546 | SP1 template (plasmid) | Template | CTGGGCGGAAACCAAGTACGCAACTTGCTCTTACACGCCTCAGCGAGAGAGCG AGTCCTACCATTGGGTAGGCAGCCCGCCTTTCCTCTGCAAGGCCCTCCTTTCA CCCTCCCTCATTGGGCGGGGCAGTAGATAAGGGGCGGGGATTAGGCCGGGCTT GTGGTGCGCTGCTCCCTCCTCCTTACCCCCCCCTCCCTGTCCGGTCCGGGTTC GCTTGCCTCGTCAGCGTCCGCGTTTTTCC |
| o5241 | 8NX template | Template | TCACCCTTTCATTCATTCCCTCCACACAACAACCATTCTTNNNNCGNNNNAAT GTGAGGAGGGTGTTATAGAATTGAGTAGTAAAGGAGAAG |
| o5242 | NX fw | PCR primer | [BTNTG]TCACCCTTTCATTCATTCCCTCCACACAACAACCATTC |
| o5243 | NX rv | PCR primer | CTTCTCCTTTACTACTCAATTCTATAACACCCTCCTCAC |
| o5246 | 12NX template | Template | TCACCCTTTCATTCATTCCCTCCACACAACAACCATTCTTNNNNCGNNNNCGN NNNAATGTGAGGAGGGTGTTATAGAATTGAGTAGTAAAGGAGAAG |
| o5247 | 16NX template | Template | TCACCCTTTCATTCATTCCCTCCACACAACAACCATTCTTNNNNCGNNNNCGN NNNCGNNNNAATGTGAGGAGGGTGTTATAGAATTGAGTAGTAAAGGAGAAG |
| o5688 | VEGFA split 1 fw | EMSA probe | [6-FAM]TTTCCAAAGCCCATTCCCTCTTTAGCCAGAGCCGGGGT |
| o5689 | VEGFA split 1 rv | EMSA probe | ACCCCGGCTCTGGCTAAAGAGGGAATGGGCTTTGGAAA |
| o5690 | VEGFA split 2 fw | EMSA probe | [6-FAM]GAGCCGGGGTGTGCAGACGGCAGTCACTAGGGGGCGCT |
| o5691 | VEGFA split 2 rv | EMSA probe | AGCGCCCCCTAGTGACTGCCGTCTGCACACCCCGGCTC |
| o5692 | VEGFA split 3 fw | EMSA probe | [6-FAM]AGGGGGCGCTCGGCCACCACAGGGAAGCTGGGTGAATG |
| o5693 | VEGFA split 3 rv | EMSA probe | CATTCACCCAGCTTCCCTGTGGTGGCCGAGCGCCCCCT |
| o5694 | VEGFA split 4 fw | EMSA probe | [6-FAM]TGGGTGAATGGAGCGAGCAGCGTCTTCGAGAGTGAGGA |
| o5695 | VEGFA split 4 rv | EMSA probe | TCCTCACTCTCGAAGACGCTGCTCGCTCCATTCACCCA |
| o5696 | VEGFA split 5 fw | EMSA probe | [6-FAM]CGAGAGTGAGGACGTGTGTGTCTGTGTGGGTGAGTGAG |
| o5697 | VEGFA split 5 rv | EMSA probe | CTCACTCACCCACACAGACACACACGTCCTCACTCTCG |
| o5698 | VEGFA split 6 fw | EMSA probe | [6-FAM]GGTGAGTGAGTGTGTGCGTGTGGGGTTGAGGGCGTTGG |
| o5699 | VEGFA split 6 rv | EMSA probe | CCAACGCCCTCAACCCCACACGCACACACTCACTCACC |
| o5700 | VEGFA split 7 fw | EMSA probe | [6-FAM]TTGAGGGCGTTGGAGCGGGGAGAAGGCCAGGGGTCACT |
| o5701 | VEGFA split 7 rv | EMSA probe | AGTGACCCCTGGCCTTCTCCCCGCTCCAACGCCCTCAA |

| | | | |
|---|---|---|---|
| o5917 | E-box fw | EMSA probe | [6-FAM]TAGGCCAXGTGGGAGG, X = C, mC or hmC |
| o5918 | E-box rv | EMSA probe | CCTCCCAXGTGGCCTA, X = C, mC or hmC |
| o6051 | X-box 2 (physiological) fw | EMSA probe | [6-FAM]CCTAGTTGCCXGGCAACCCCC, X = C, mC or hmC |
| o6052 | X-box 2 (physiological) rv | EMSA probe | GGGGGTTGCXGGGCAACTAGG, X = C, mC or hmC |
| o6053 | X-box 1 fw | EMSA probe | [6-FAM]CCTGATGAXGACGTACCG, X = C, mC or hmC |
| o6054 | X-box 1 rv | EMSA probe | CGGTACGTXGTCATCAGG, X = C, mC or hmC |
| o6107 | VEGFA synthetic fragment fw | EMSA probe | [6FAM]GCTGGGTGAATGGAGXGAGCAGXGTCTTXGAGAGTGAGGAXGTGTGT GTCTGTGTGG, X=C |
| o6108 | VEGFA synthetic fragment rv | EMSA probe | CCACACAGACACACACAXGTCCTCACTCTXGAAGAXGCTGCTXGCTCCATTCACC CAGC, X=C |
| o6109 | VEGFA synthetic fragment fw | EMSA probe | [6FAM]GCTGGGTGAATGGAGXGAGCAGXGTCTTXGAGAGTGAGGAXGTGTGT GTCTGTGTGG, X=mC |
| o6110 | VEGFA synthetic fragment rv | EMSA probe | CCACACAGACACACACAXGTCCTCACTCTXGAAGAXGCTGCTXGCTCCATTCACC CAGC, X=mC |
| o6111 | VEGFA synthetic fragment fw | EMSA probe | [6FAM]GCTGGGTGAATGGAGXGAGCAGXGTCTTXGAGAGTGAGGAXGTGTGT GTCTGTGTGG, X=hmC |
| o6112 | VEGFA synthetic fragment rv | EMSA probe | CCACACAGACACACACAXGTCCTCACTCTXGAAGAXGCTGCTXGCTCCATTCACC CAGC, X=hmC |

**Table S6**
Oligonucleotides used for CDS amplification of full-length proteins and Gibson assembly. Gibson overhangs aligning with the backbone p1379 (see **Fig. S15**) are underlined.

| Name | Description | Sequence 5' -> 3' |
|---|---|---|
| o5529 | ATF2 fw | GAAAATCTTTATTTTCAGTCTCTCATGAAATTCAAGTTACATGTGAATTCTG |
| o5530 | ATF2 rv | CAGTGGTGGTGGTGGTGGTGCTCACTTCCTGAGGGCTGTG |
| o5539 | CHAF1B fw | GAAAATCTTTATTTTCAGTCTCTCATGAAAGTCATCACTTGTGAAATAGC |
| o5540 | CHAF1B rv | CAGTGGTGGTGGTGGTGGTGCTCAGGGTCCAGACTTTCCG |
| o5545 | FOXA1 fw | GAAAATCTTTATTTTCAGTCTCTCATGTTAGGAACTGTGAAGATGGAAG |
| o5546 | FOXA1 rv | CAGTGGTGGTGGTGGTGGTGCTCGGAAGTGTTTAGGACGGGTC |
| o5555 | L3MBTL3 fw | GAAAATCTTTATTTTCAGTCTCTCATGACTGAATCTGCCTCTAGC |
| o5556 | L3MBTL3 rv | CAGTGGTGGTGGTGGTGGTGCTCAAGTTCATTGTGAGAATTCTTCTCTG |
| o5557 | MAX fw | GAAAATCTTTATTTTCAGTCTCTCATGAGCGATAACGATGACATCG |
| o5558 | MAX rv | CAGTGGTGGTGGTGGTGGTGCTCGCTGGCCTCCATCCG |
| o5561 | MYC fw | GAAAATCTTTATTTTCAGTCTCTCATGCCCCTCAACGTTAGC |
| o5562 | MYC rv | CAGTGGTGGTGGTGGTGGTGCTCCGCACAAGAGTTCCGTAG |
| o5567 | RFX5 fw | GAAAATCTTTATTTTCAGTCTCTCATGGCAGAAGATGAGCCTG |
| o5568 | RFX5 rv | CAGTGGTGGTGGTGGTGGTGCTCTGGGGGTGTTGCTTTTGG |
| o5571 | RFXANK fw | GAAAATCTTTATTTTCAGTCTCTCATGGAGCTTACCCAGCC |
| o5572 | RFXANK rv | CAGTGGTGGTGGTGGTGGTGCTCCTCAGGGTCAGCGGG |
| o5579 | SUB1 fw | GAAAATCTTTATTTTCAGTCTCTCATGCCTAAATCAAAGGAACTTGTTTC |

| | | |
|---|---|---|
| o5580 | SUB1 rv | CAGTGGTGGTGGTGGTGGTGCTCCAGTTTTCTTACTGCATCATCAATG |
| o5591 | TFAM fw | GAAAATCTTTATTTTCAGTCTCTCATGGCGTTTCTCCGAAGC |
| o5592 | TFAM rv | CAGTGGTGGTGGTGGTGGTGCTCACACTCCTCAGCACCATATTTTC |

**Table S7**
Oligonucleotides used for cloning of MAX fl, MYC bHLHZ and MBP-RFX5 DBD.

| Name | Description | Sequence 5’ -> 3’ |
|---|---|---|
| o5938 | MAX fl fw | ATCGAGGGAAGGCTCGAAAATC |
| o5939 | MAX fl rv | CGCCAGCGCCACTTTATC |
| o5980 | MYC bHLHZ fw 1 | ATGGTCAAGAGGCGAACACACAAC |
| o5981 | MYC bHLHZ rv 1 | GAGCCTTCCCTCGATCGC |
| o5983 | MYC bHLHZ fw 2 | GAGCACCACCACCACCACCACT |
| o5984 | MYC bHLHZ rv 2 | TAGCTGTTCAAGTTTGTGTTTCAACTGTTCTCG |
| o6035 | MBP-RFX5_DBD fw 1 | GAGCACCACCACCACCACCACTG |
| o6036 | MBP-RFX5_DBD rv 1 | GGTCTTCCTCCTTATGCCACTGTAGC |
| o6037 | MBP-RFX5_DBD fw 2 | GGAGACAAAAGCTCAGAGCCAAGT |
| o6038 | MBP-RFX5_DBD rv 2 | GAGAGACTGAAAATAAAGATTTTCGAGCCTTCC |

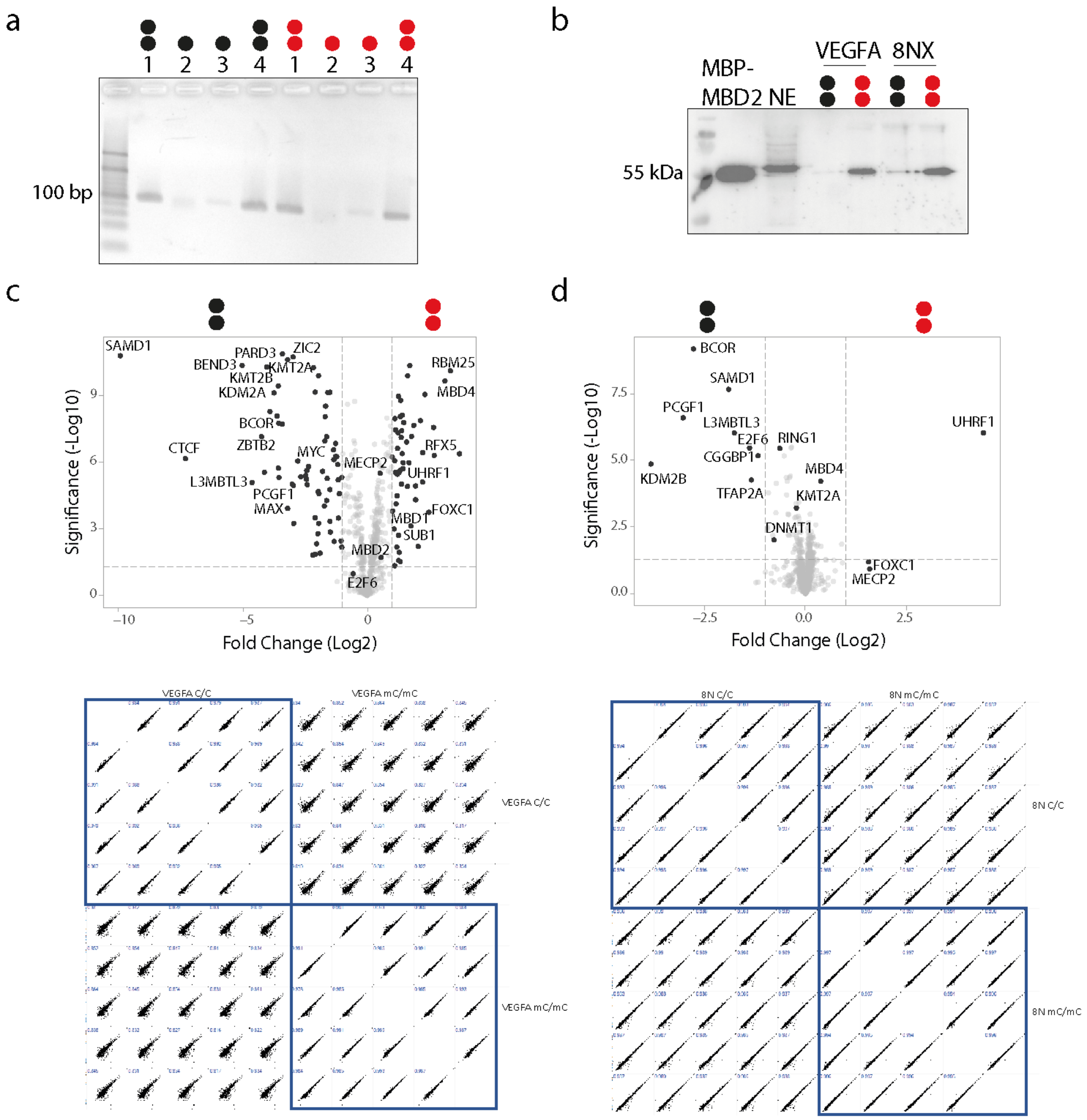


e

| probe | VEGFA | 8NX |
|---|---|---|
| concentration | 13 nM (4 pmol) | 160 nM (48 pmol) |
| Total # of CpGs | $2.89*10^{13}$ | |
| HEK293T nuclear etxracts | 3.6 µM (50 µg) | |
| Competitor dA:dT DNA | 1.1 µM (5 µg) | |
| MBP-MBD2 (only for Western blot pulldowns) | 1 nM (16 ng) | |

**Figure S1. Comparative pulldown experiments with VEGFA and random 8N probe and HEK293T nuclear extracts containing MBD2-MBP spike-in protein analyzed by anti-MBP western blots.** (a) Agarose gel validation of probe generation steps for probes containing a random NNNNCGNNNN target sequence ("8NX"; steps are identical to the ones shown in **Fig. 2c** of the main manuscript). Employed oligonucleotides were o5241 – o5243 (see **Table S5**). (b) Pulldowns were conducted with mixes shown in (e). Note that different absolute probe concentrations were used for VEGFA and 8NX probes to account for the different numbers of CpGs in the two probe sequences. As also visible in **Fig. 2c** of the main manuscript, a higher molecular weight off-target protein is bound by the antibody (lane NE). Both probes lead to comparable amounts of enriched MBD2-MBP protein and similar mC/mC versus C/C selectivity. (c) Volcano plot showing enriched proteins with the VEGFA probe (p-value < 0.05 and log2 fold change >1) between unmethylated (black circles) and methylated (red circles) CpG dyads. Reproducibility data are shown as multiscatter plots for the technical replicates. (d) As (c), but for the 8NX probe.

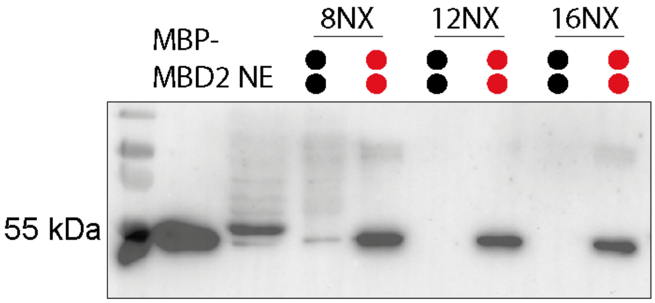


| probe | # seq. contexts | # CpGs | pmol used in pulldown | Seq. Coverage | Total CpGs |
|---|---|---|---|---|---|
| 8NX | 65536 | 1 | 48 | 446577758 | $2.89*10^{13}$ |
| 12NX | 16777216 | 2 | 24 | 972222 | |
| 16NX | 4294967296 | 3 | 16 | 2271 | |

**Figure S2. Comparative pulldown experiments with random probes and HEK293T nuclear extracts containing human MBP-MBD2 spike-in protein analyzed by anti-MBP Western blots**. Three different probes containing one to three NNNNCGNNNN random target sequences were employed ("8NX", "12NX" and "16NX"). Pulldowns were conducted as in **Fig. S1** with different absolute probe concentrations to account for the different numbers of CpGs in the probe sequences (compare data of table, column 4).

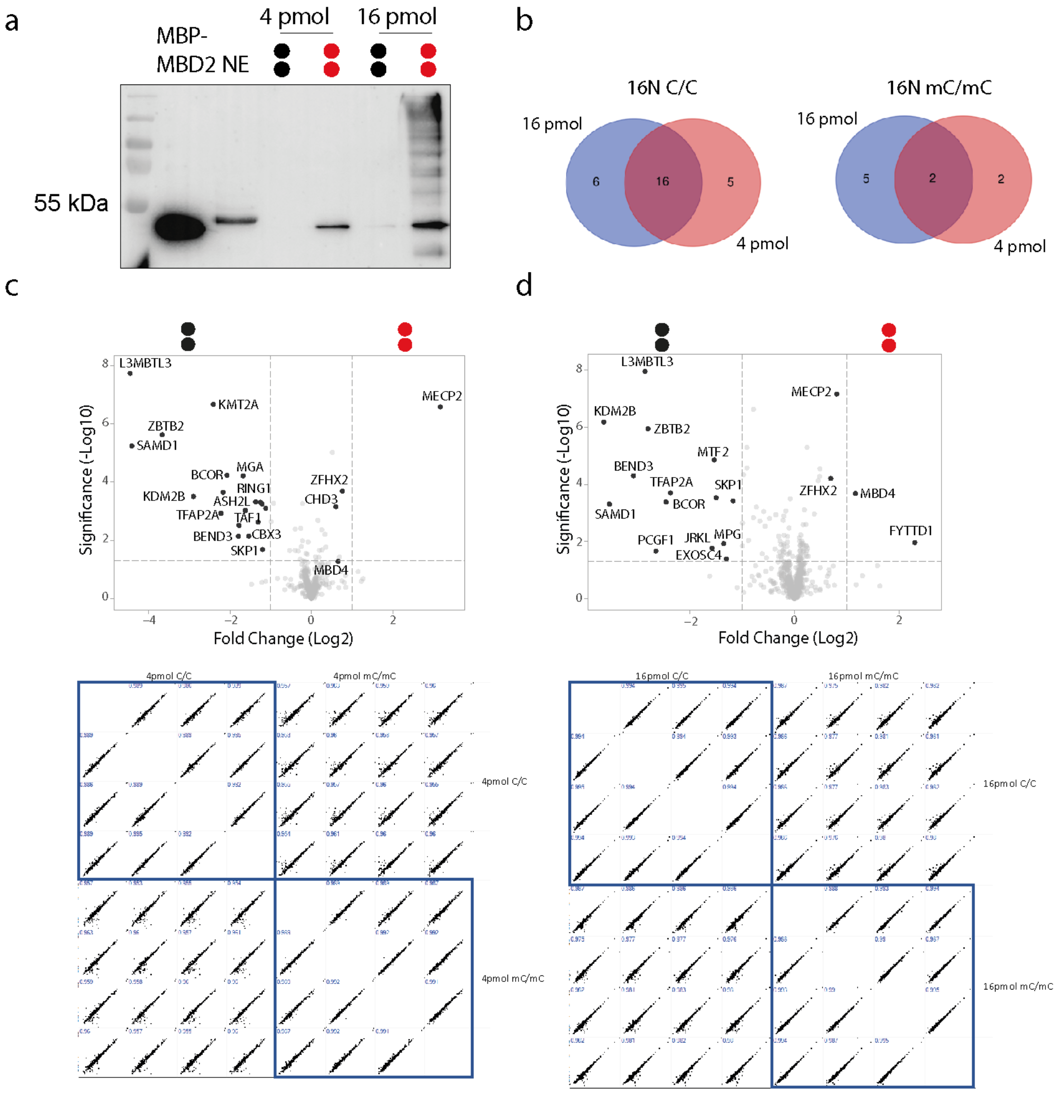

**Figure S3. Comparative pulldown experiments with the random 16NX probe and HEK293T nuclear extracts containing MBP-MBD2 spike-in protein.** The probe contains three NNNNCGNNNN random target sequences. Pulldowns were conducted as in **Fig. S1** with different absolute probe concentrations to account for the different numbers of CpGs in the probe sequences. (a) Anti-MBP Western blot showing higher enrichment of the MBD2-MBP protein with more probe concentration, however, off-target binding was enhanced as well. mC/mC and C/C selectivity were in the same range. (b) Venn diagrams showing number of enriched proteins with two different 16NX concentrations. (c) Volcano plot showing enriched proteins with a lower concentration of 16NX (4 pmol in total) (p-value $< 0.05$ and log2 fold change $>1$) between unmethylated (black circles) and methylated (red circles) CpG dyads. Reproducibility data are shown as multiscatter plots for the technical replicates. (d) As (c) but with 16 pmol of the 16NX probe.

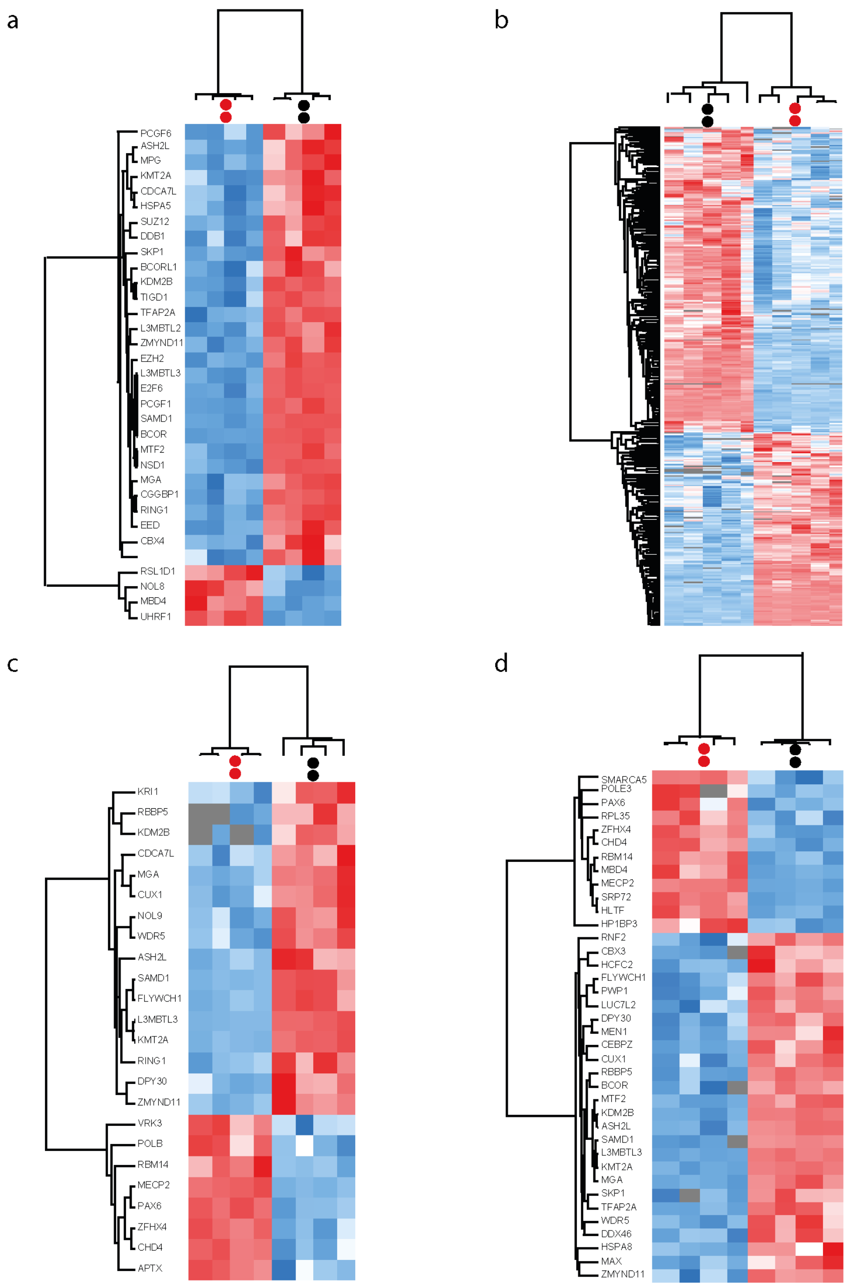


**Figure S4: Heatmaps of pulldowns corresponding to Fig. S1 and S3.** The heatmaps show correlation-based clustering of the LFQ intensities after log2 transformation and normalization by row mean subtraction. Proteins included in the clustering significantly bind unmethylated (black circles) or methylated (red circles) probe as determined by an ANOVA test (p-value $< 0.05$, $S0 = 0$). Blue indicates depletion, whereas enrichment is indicated in red. Heatmaps corresponding to pulldowns with (a) 8NX, (b) VEGFA, (c) 4 pmol of 16NX and (d) 16 pmol of 16NX. Note that gene names were omitted for the VEGFA enrichment due to large quantity.

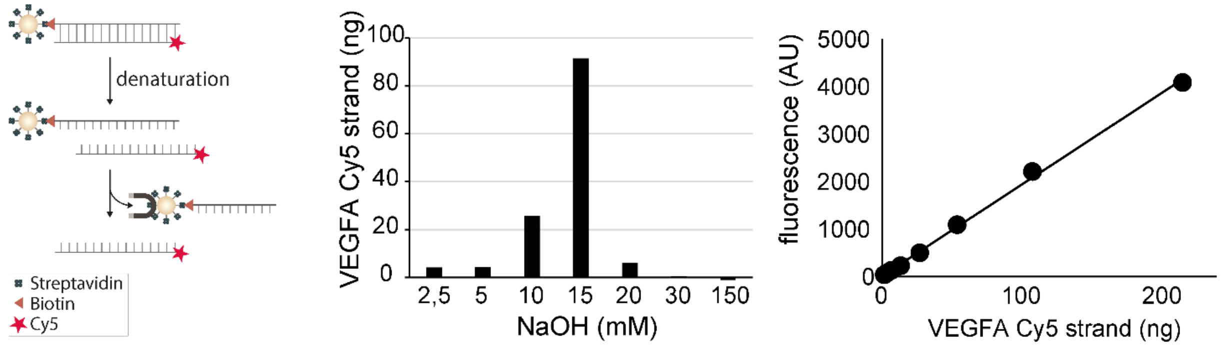


**Figure S5. Optimization/characterization of denaturation protocol for obtaining the antisense strand of asymmetric probes serving as template in the final primer extension step.** Left: scheme of denaturation protocol. Middle: Quantities of single strand obtained in individual, step-wise denaturing elution using increasing NaOH concentrations. A NaOH concentration of 20 mM was identified as optimal and used in later asymmetric probe generations. Right: calibration curve for quantification of obtained single strand via Cy5 fluorescence.

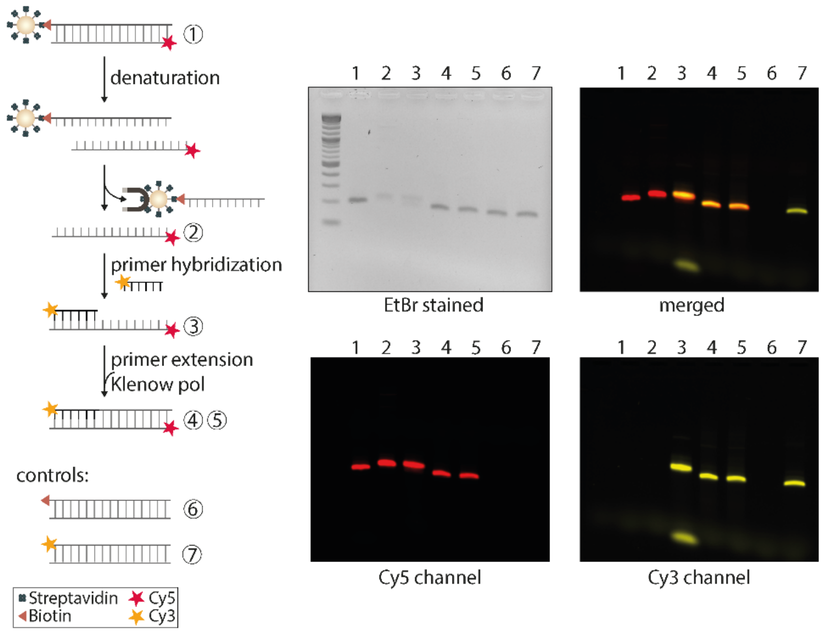


**Figure S6. Analysis of final primer extension step in VEGFA asymmetric probe generation protocol by agarose gel electrophoresis with fluorescently labeled DNA.** Compared to analyses based on ethidium bromide (EtBr) staining (e.g., as shown in **Fig. 2d** of the manuscript), this analysis allows for quantitative and far more sensitive analysis of both DNA strands. Left: Experiment scheme. Right: agarose gels with signaling as indicated below. Note that the ssDNA template (lane 2) runs higher than a perfectly double-stranded DNA of the same length in the EtBr channel (compare with lanes 1, 6, and 7 showing PCR products generated with unmodified dCTP and primers introducing Cy5, biotin, or Cy3, respectively). The hybridization of a primer to the ssDNA template (lane 3, excess unhybridized primer visible in Cy3 channel in lower gel region) results in a high running band, as well. Lanes 4 and 5 contain final primer extension products generated with either unmodified dCTP (4) or modified dhmCTP (5) and do not show a Cy3 signal above the bright and defined bands matching the running height of dsDNA (i.e., lanes 1, 7, which may result from not or only partially extended primer). This data collectively indicates a completed final primer extension step for the asymmetric probe bearing hmC, yielding a fully double-stranded asymmetric probe.

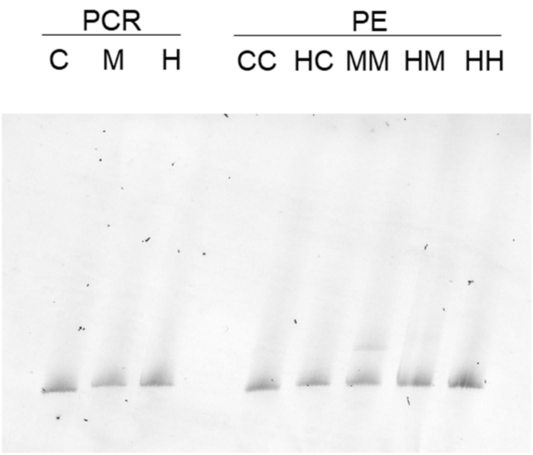


**Figure S7. PAGE analysis of VEGFA enrichment probes.** VEGFA PCR products were generated using dCTP, dmCTP or dhmCTP and a 5´-FAM-labeled primer, and analyzed by 8.3 M urea denaturing sequencing PAGE (8% acrylamide, run length ~45 cm, shown in first three lanes). In addition, symmetric and asymmetric DNA probes were generated by the protocol shown in **Fig. 2c** of the manuscript using ssDNA templates containing C, mC or hmC, a 5´-FAM-labeled primer, and dCTP, dmCTP or dhmCTP in the primer extension step (shown in the last five lanes). The gel shows that all primer-extension generated probes exhibit the same length as the PCR products conducted with nonmodified dCTP and are thus fully double-stranded.

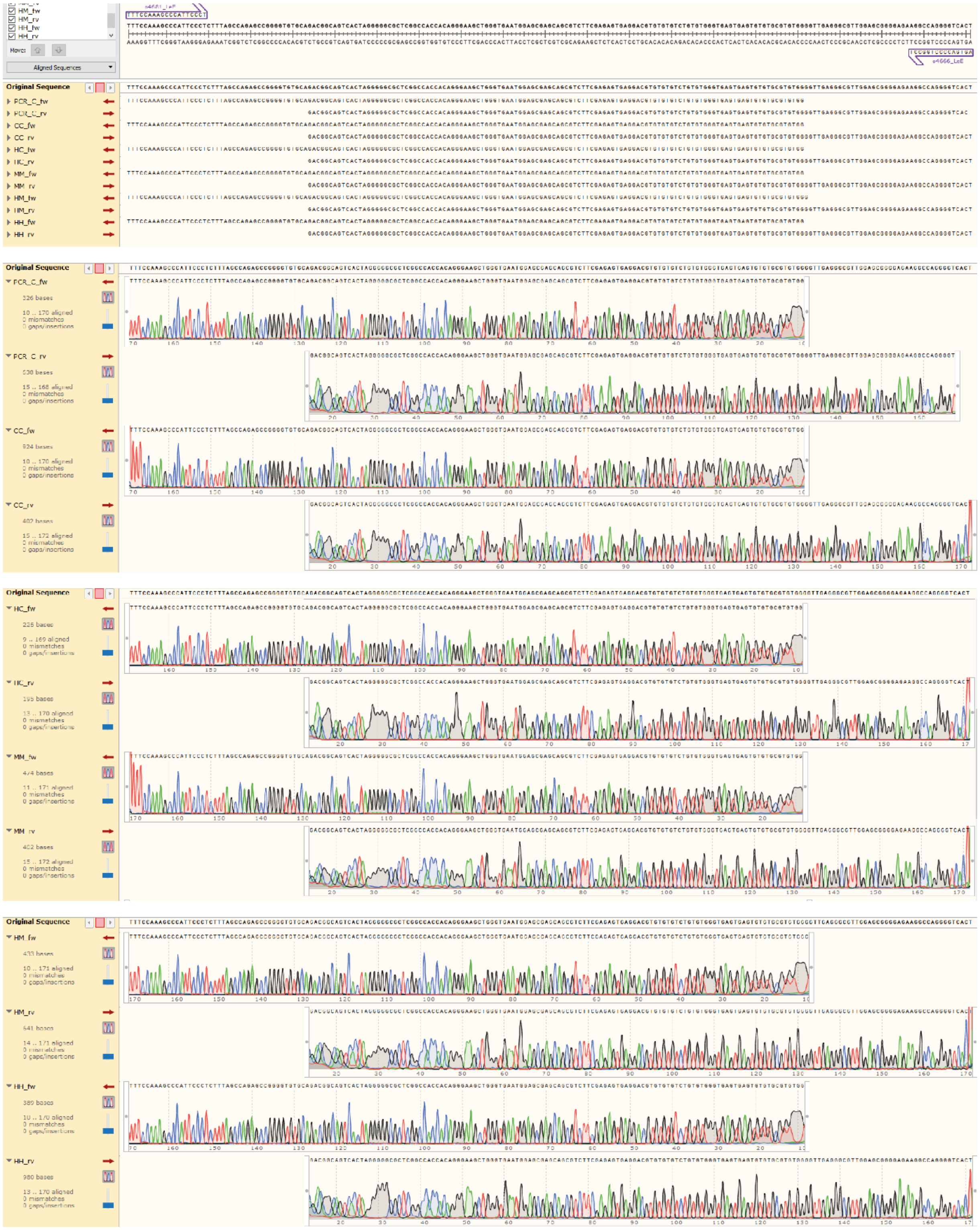
Original Sequence
PCR_C_fw
PCR_C_rv
CC_fw
CC_rv
HC_fw
HC_rv
MM_fw
MM_rv
HM_fw
HM_rv
HH_fw
HH_rv
Aligned Sequences

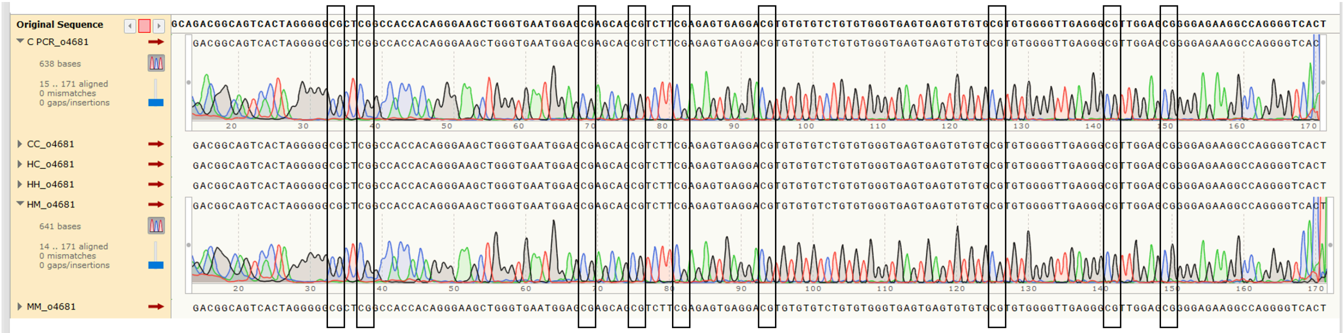


**Figure S8. Sequence verification of final probes.** Both the forward and reverse strand of a regular PCR-generated VEGFA probe without the use of modified dCTPs (PCR_C) was sequenced as reference. In addition, all five asymmetric VEGFA probes generated by the primer-extension-based probe generation protocol (Fig. 2c of the manuscript) were directly subjected to the same forward and reverse Sanger sequencing as the PCR reference (C/C: CC, hmC/C: HC, hmC/hmC: HH, hmC/mC: HM; mC/mC: MM). The sequencing traces do not show differences between the PCR reference and the unmodified or modified probes, and thus no indication for misincorporation steps caused by modified dCTPs. This proves the sequence integrity of the employed probes as well as their quantitative labelling with the indicated modified dCTPs.

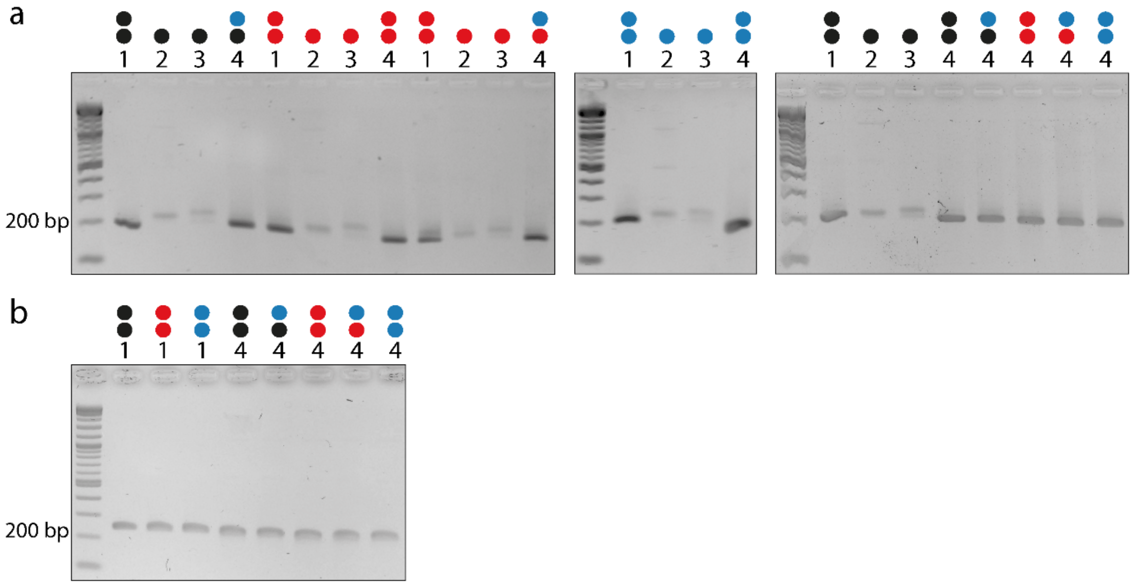


**Figure S9. Agarose gel analyses of all employed probes used in this study.** Samples were collected during the process of generating VEGFA (a) and Sp1 (b) probes and run on a 2 % agarose gel with a 1 kb Plus DNA ladder (New England Biolabs). The numbering corresponds to **Fig. 2c** in the manuscript representing 1: the PCR product, 2: ssDNA, 3: primer hybridized product and 4: (asymmetric) primer extended product. Circles are colored according to the C modifications (C – black, mC – red and hmC – blue) depicting ssDNA (one circle) or dsDNA (two circles).

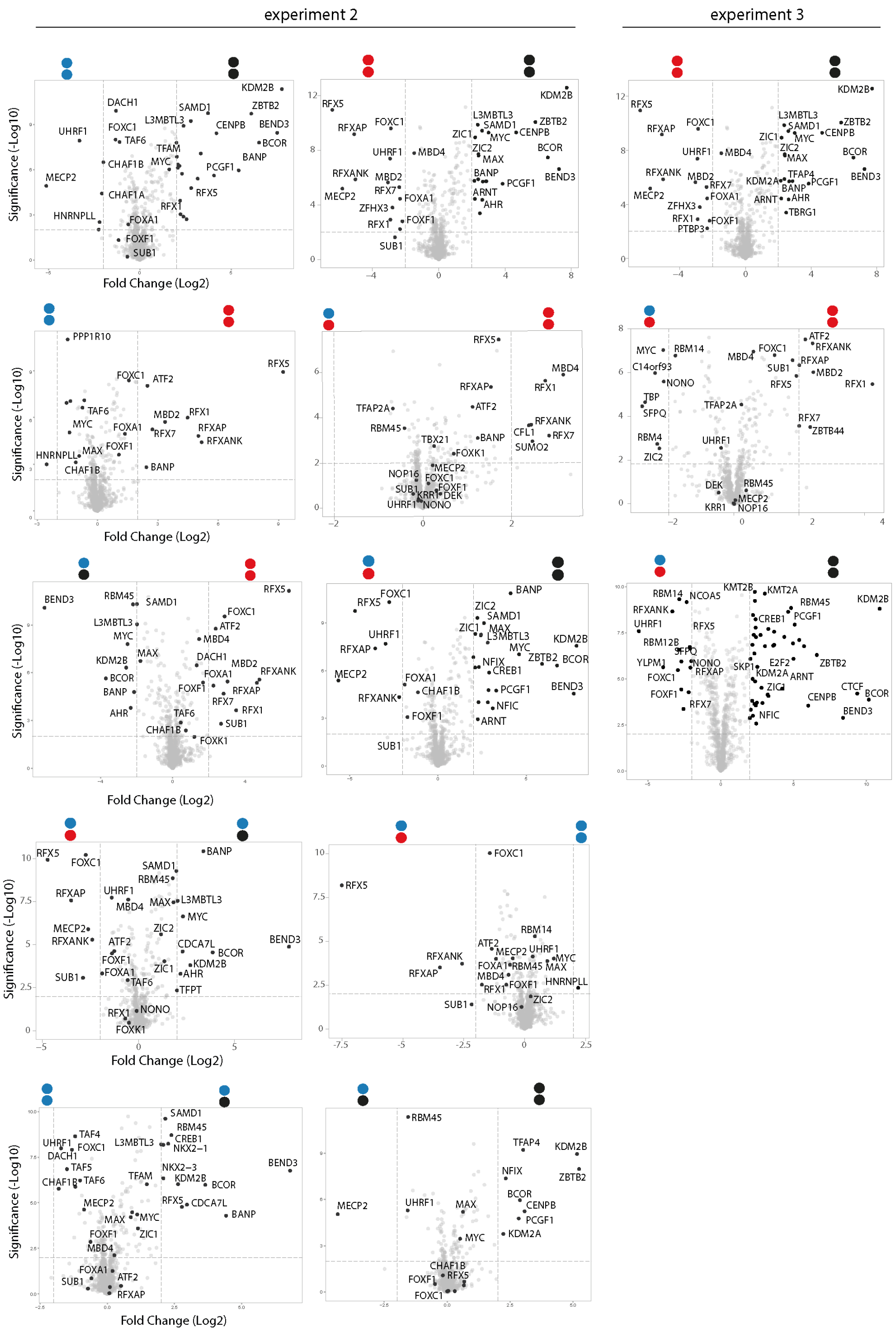


**Figure S10. Supplementary volcano plots from biological experiments 2 and 3 related to Fig. 3.** Each biological experiment contains five technical replicates. Comparison of enriched proteins (p-value < 0.01 and log2 fold change >2) between two indicated CpG dyads.

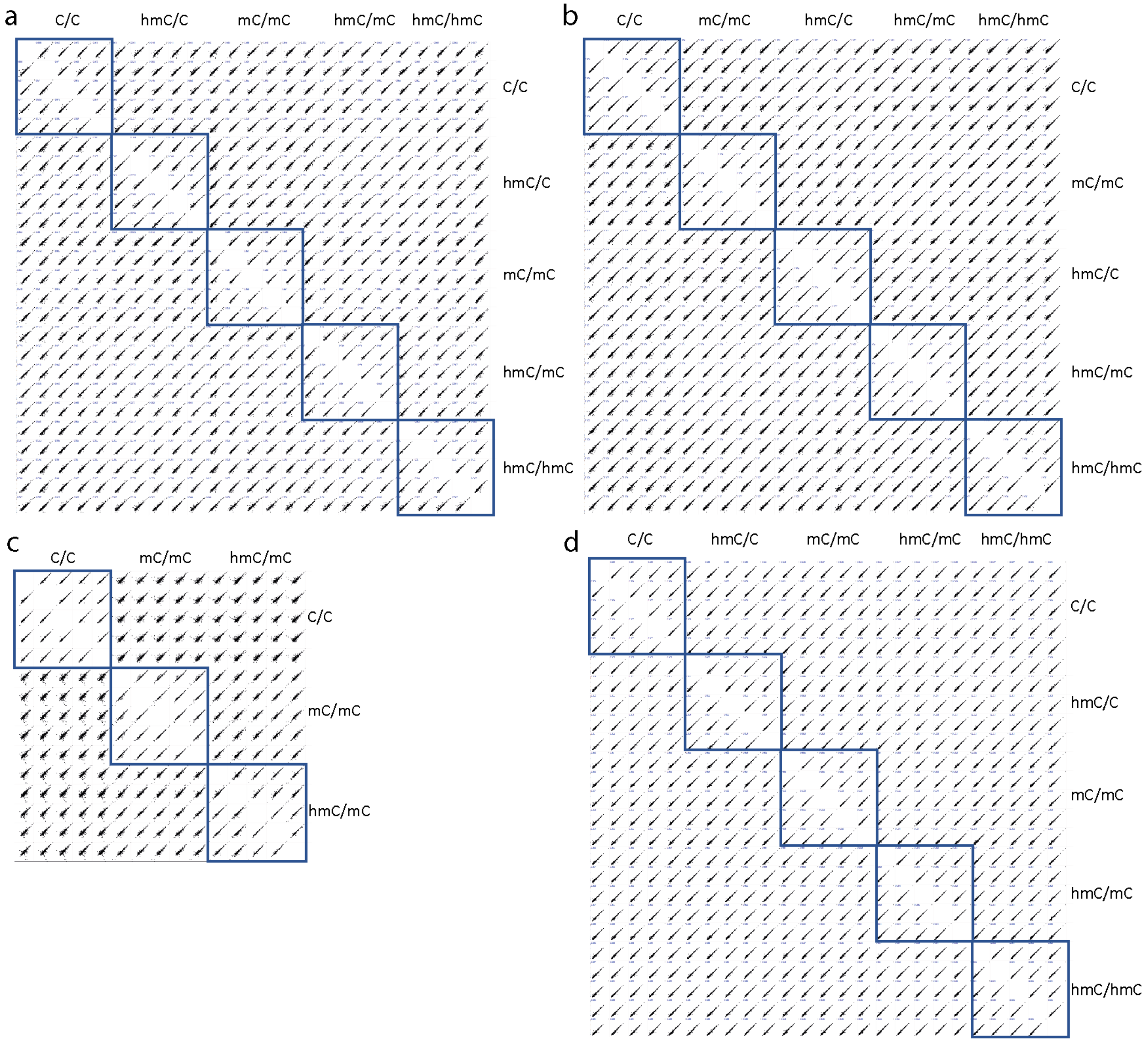


**Figure S11. Reproducibility data for enrichment/MS experiments.** Figures show multiscatter plots for the technical replicates, for each biological experiment separately. (a) HEK293T, biological experiment 1, (b) HEK293T, biological experiment 2, (c) HEK293T, biological experiment 3, (d) mouse brain.

experiment 1:

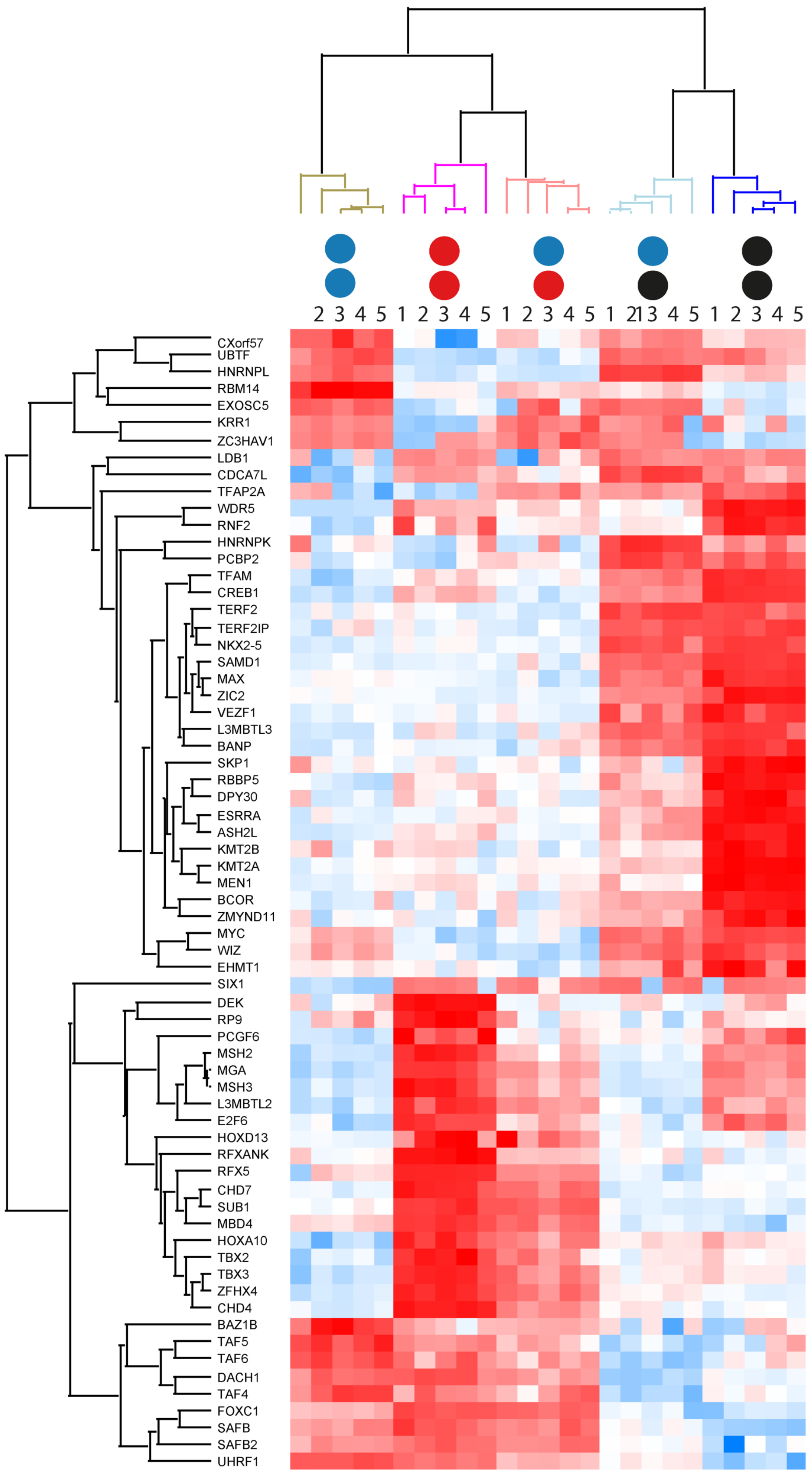

experiment 2:

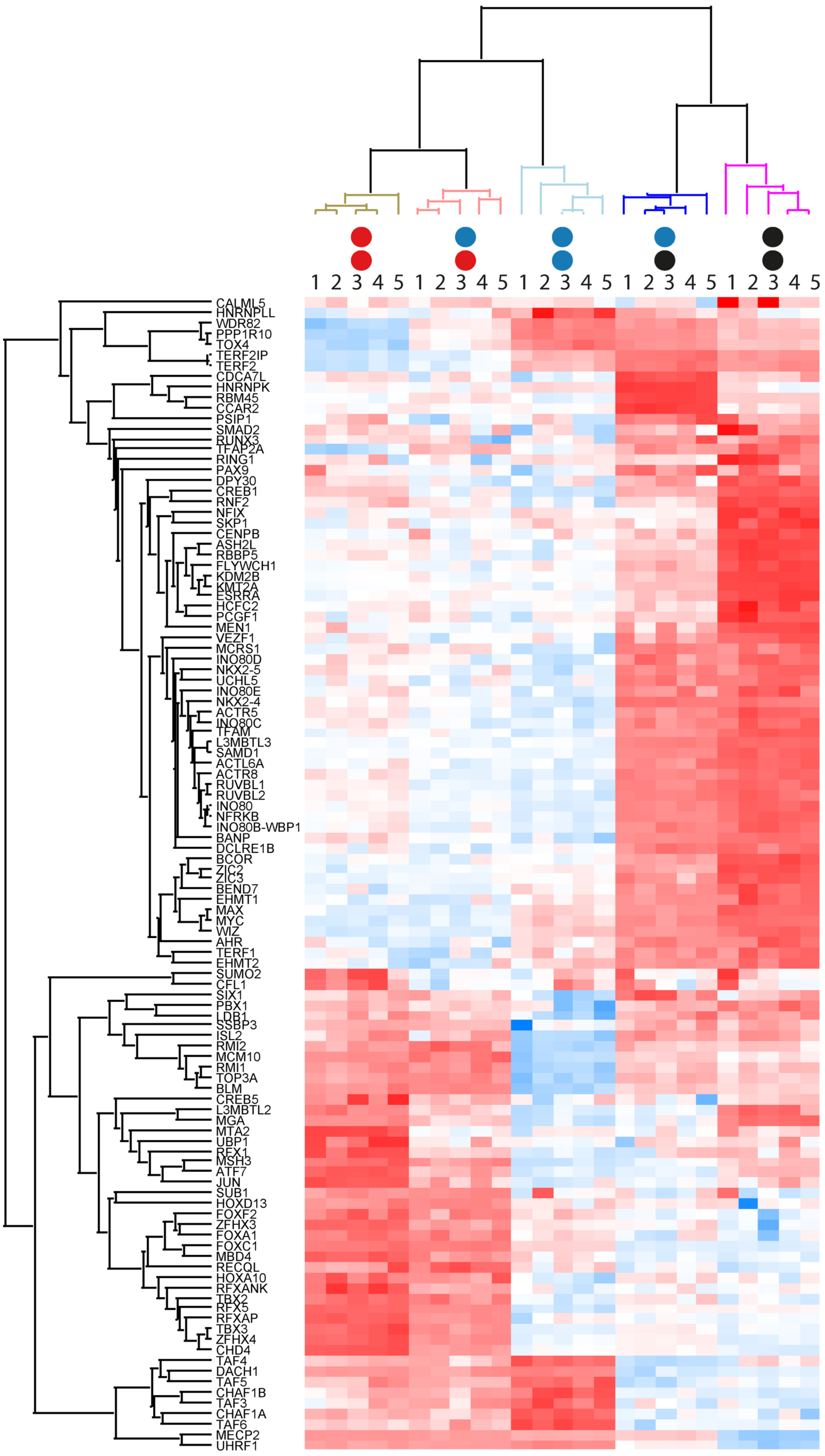

experiment 3:

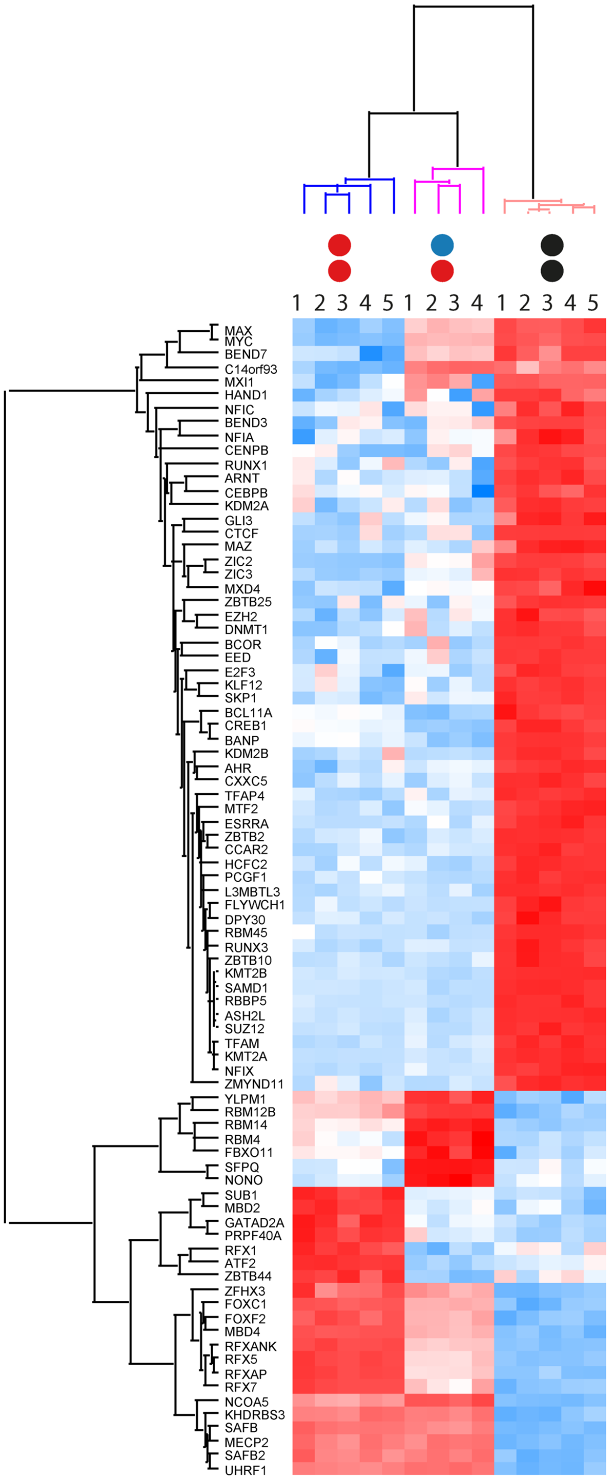


**Figure S12**. **Heatmaps of all three biological experiments corresponding to Fig. 3g** (experiment 1 is shown here again using a different color code for comparison). The heatmaps show correlation-based clustering of the LFQ intensities after log2 transformation and normalization by row mean subtraction. Proteins included in the clustering significantly bind to at least one of the modifications as determined by an ANOVA test. Blue indicates depletion, whereas enrichment is indicated in red.

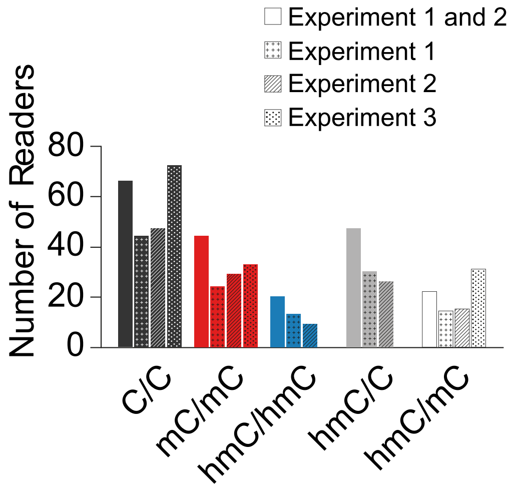


**Figure S13.** Distribution of reader numbers between different indicated VEGFA probe versions for individual HEK293T proteomics experiments. Experiment 1 corresponds to the data shown in **Fig. 3c-g** of the main manuscript. **Fig. 3a** shows that data of Experiments 1 and 2 combined as shown above.

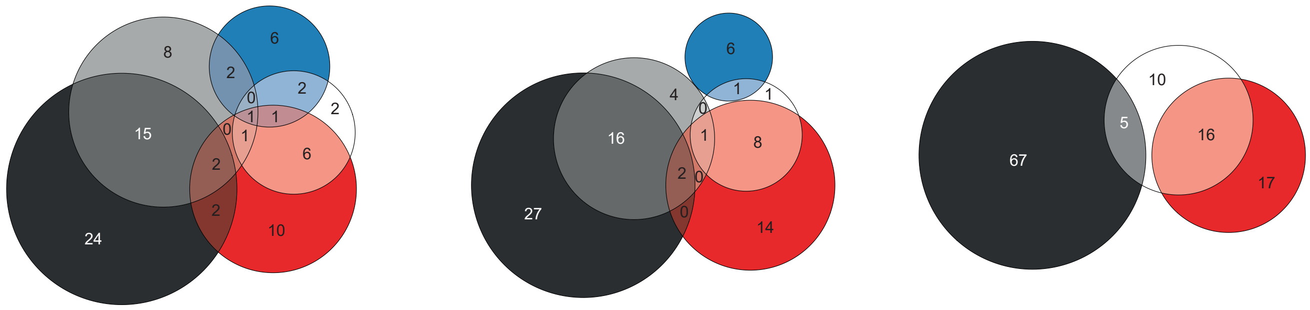

**Figure S14. Venn diagrams of individual biological experiments for proteomics data from HEK293T cells and VEGFA probe enrichments.** Shown are diagrams for experiments 1, 2 and 3 from left to right. Venn diagram in **Fig. 3b** of the manuscript corresponds to combined experiments 1 and 2.

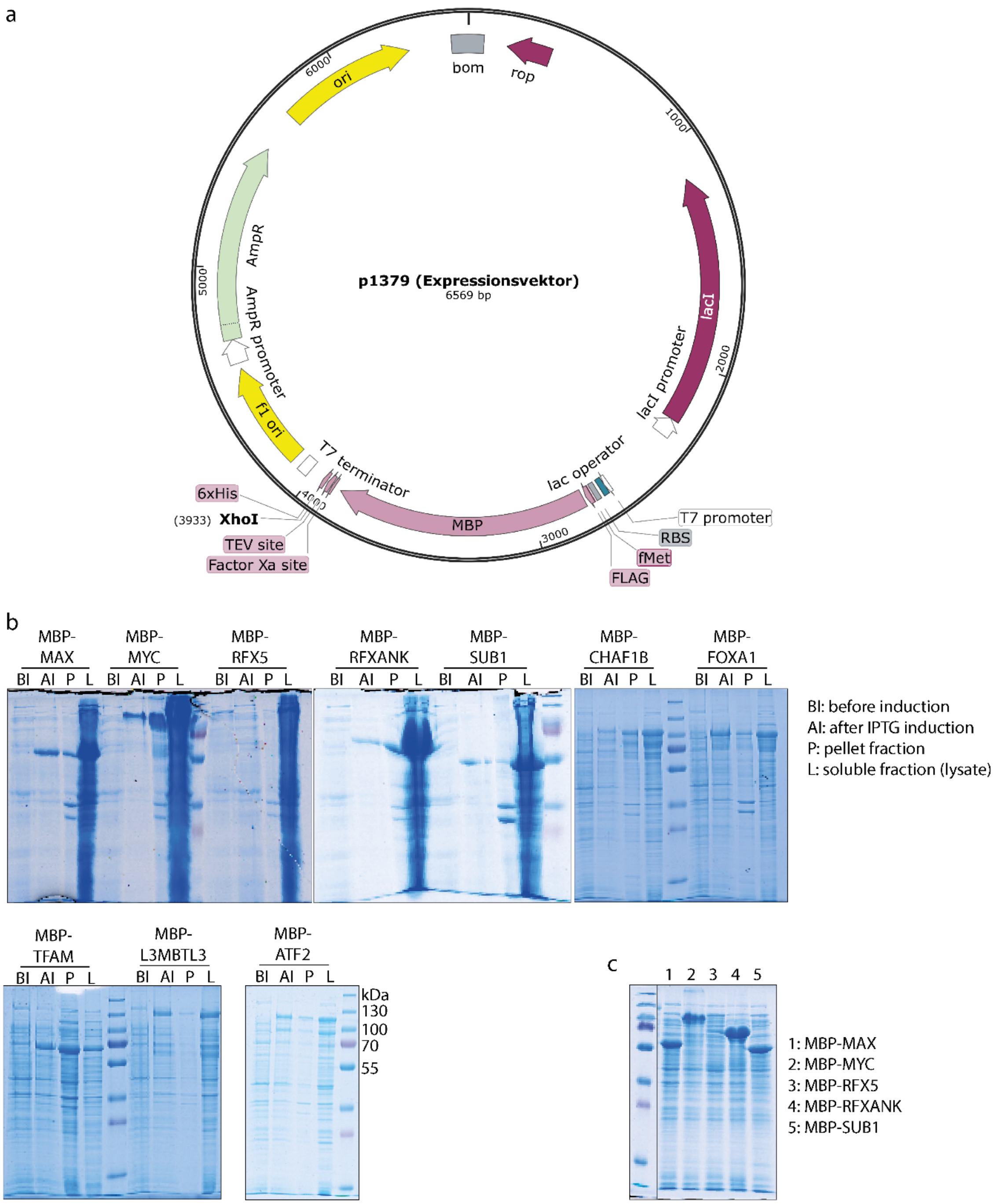


**Figure S15: Expression of candidate proteins.** (a) Plasmid map of the expression vector. The *XhoI* restriction site was used to insert the full-length coding sequences of candidate proteins that were amplified using primers from Table S5. (b) Coomassie stained 12 % SDS PAGES of expression of candidate proteins in *E. coli* BL21 DE(3) Gold showing the following fractions: before induction (BI), after induction with 1 mM IPTG (AI), pellet fraction (P) after sonication and soluble fraction (L) after sonication referred to as lysate. (c) As some soluble fractions were overloaded in (b), they were applied in smaller quantities to a new 12 % SDS PAGE.

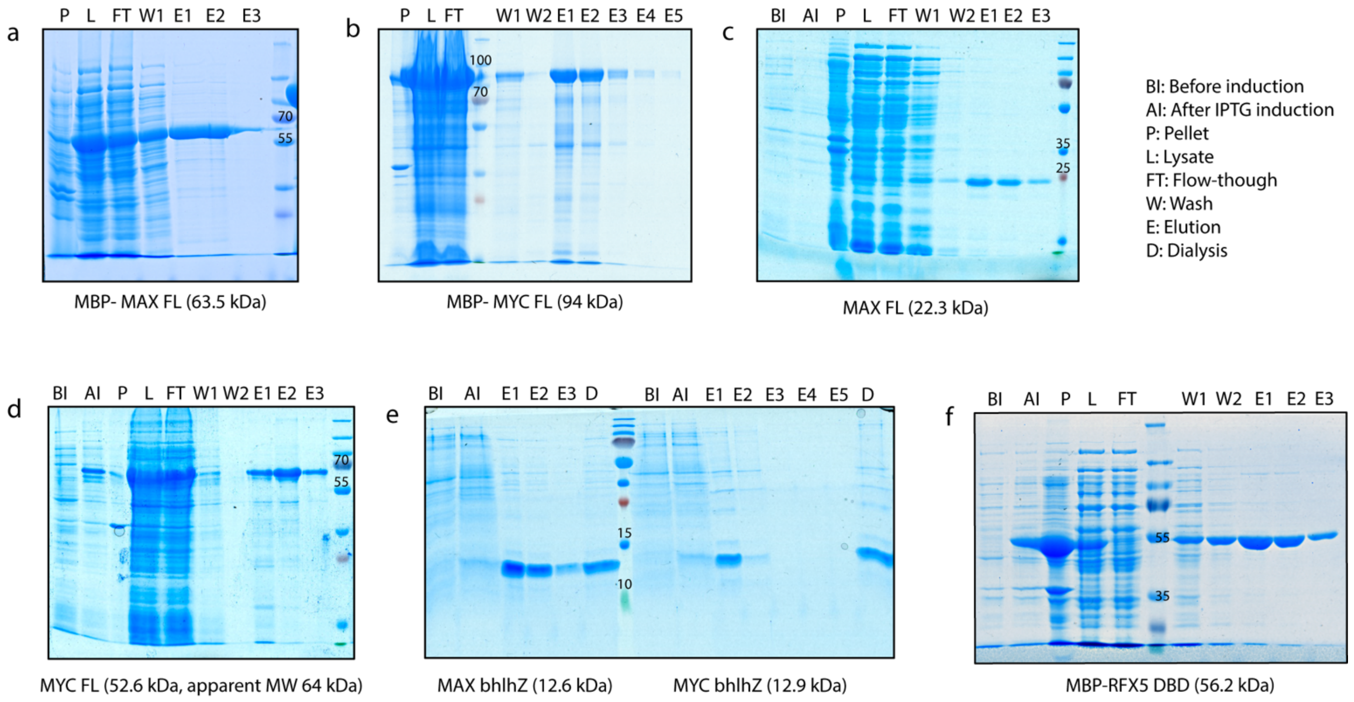


**Figure S16. Expression and purification of candidate proteins.** Coomassie stained 12 % SDS PAGES of purification of candidate proteins (a) MBP-MAX flL, (b) MBP-MYC fl, (c) MAX fl, (d) MYC fl (e) MAX bhlhZ and MYC bhlhZ), and (f) MBP-RFX5 DNA Binding Domain (DBD) in *E. coli* BL21 DE(3) Gold showing the following fractions: before induction (BI), after induction with 1 mM IPTG (AI), pellet fraction (P) after sonication and soluble fraction (L) after sonication referred to as lysate, flow-through (FT) collected after binding to the Ni-NTA resin, as well as the wash (W) and elution (E) fractions, and dialysis product (D).

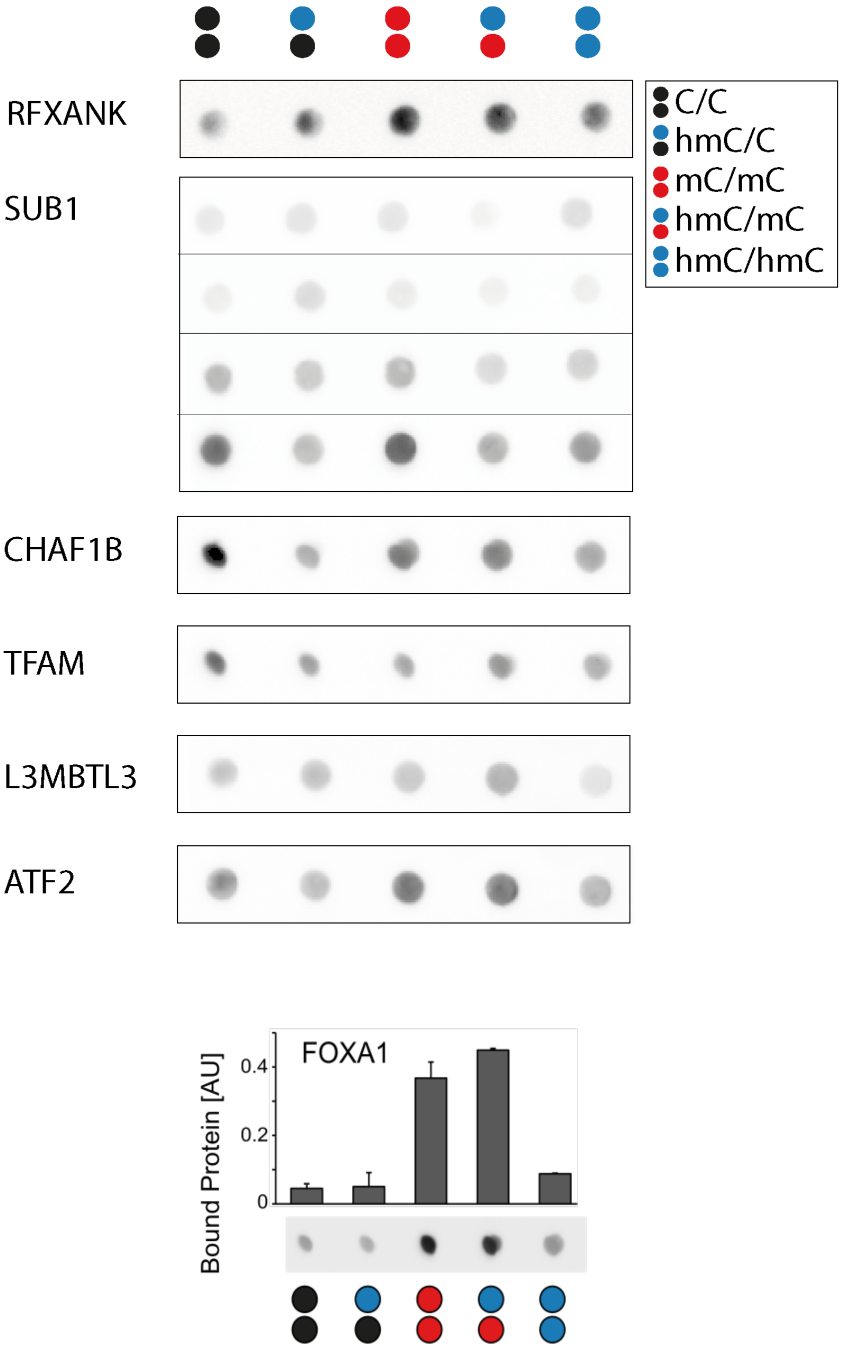


**Figure S17. Representative results from dot blot and EMSA pre-tests with candidate reader proteins.** Anti-MBP dot blot analyses of eluates from enrichments employing VEGFA probes and *E. coli* expression lysates of indicated proteins fused to an N-terminal MBP tag. Bar diagram for FOXA1 from duplicate experiments.

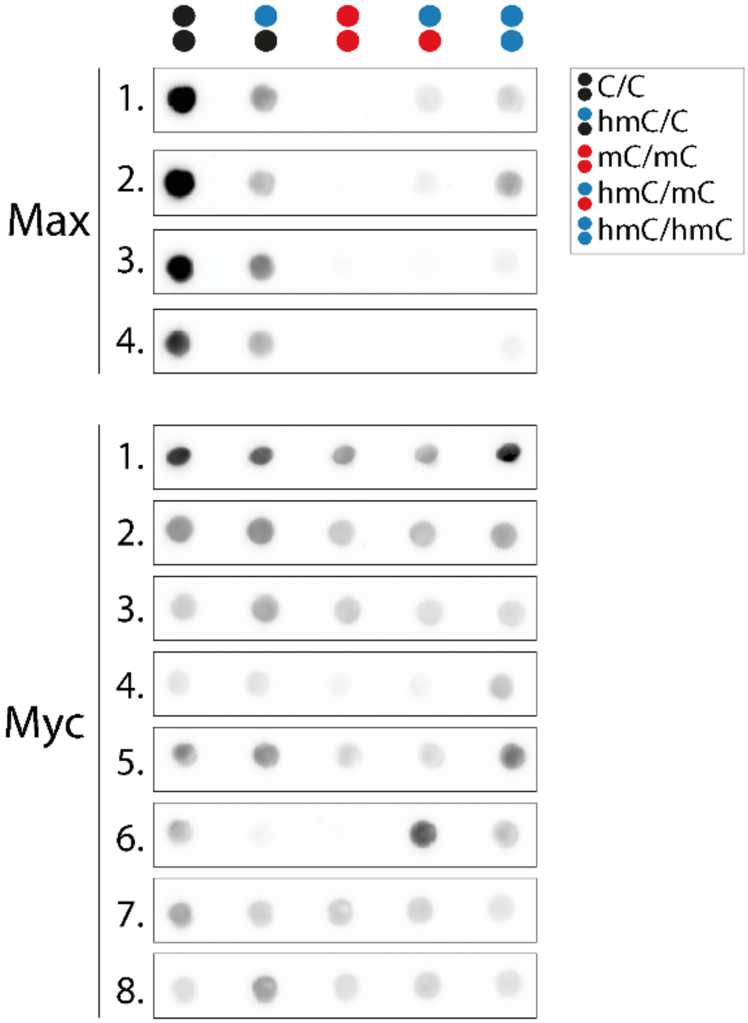


**Figure S18:** Anti-MBP dot blot analyses of eluates from enrichments employing VEGFA probes and *E. coli* expression lysates of Max and Myc fused to an N-terminal MBP tag. Multiple replicates are shown.

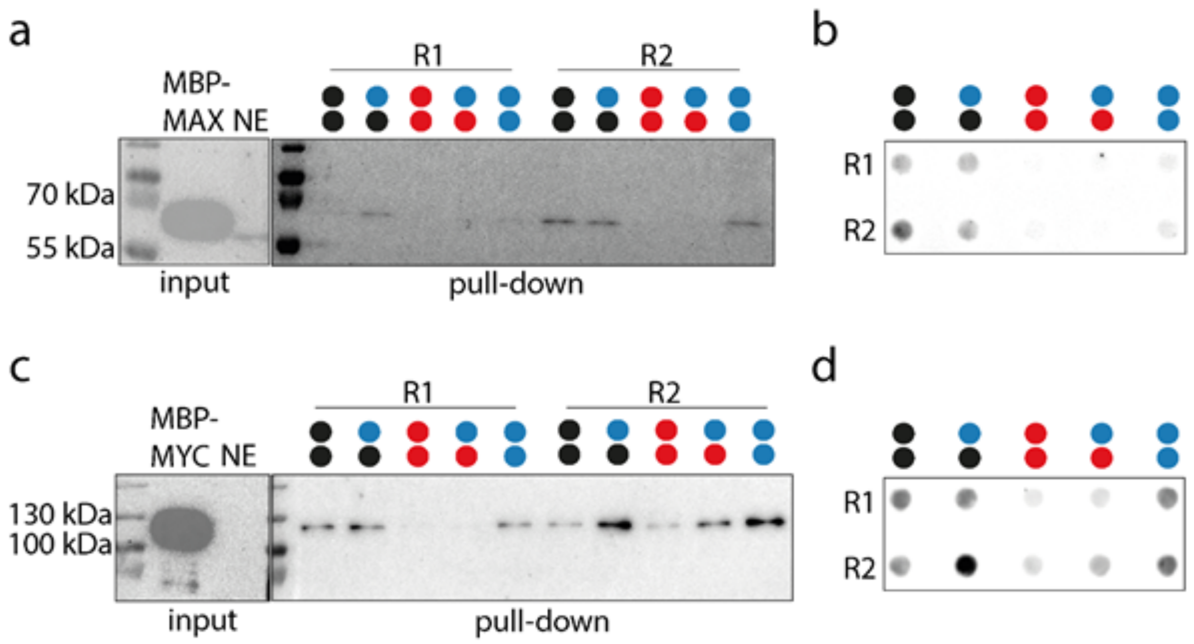


**Figure S19: Evaluation of MBP-MAX and MBP-MYC binding to modified VEGFA probes.** (a) Anti-MBP Western blot analyses of eluates from enrichments employing modified VEGFA probes (color code corresponds to **Fig. S17**) and *E. coli* expression lysate of MAX fused to an N-terminal MBP tag. Multiple replicates are shown. (b) Dot blot analyses with the same samples from (a). (c) Anti-MBP Western blot analyses of eluates from enrichments employing modified VEGFA probes (color code corresponds to **Fig. S17**) and *E. coli* expression lysate of MYC fused to an N-terminal MBP tag. Multiple replicates are shown. (d) Dot blot analyses with the same samples from (c).

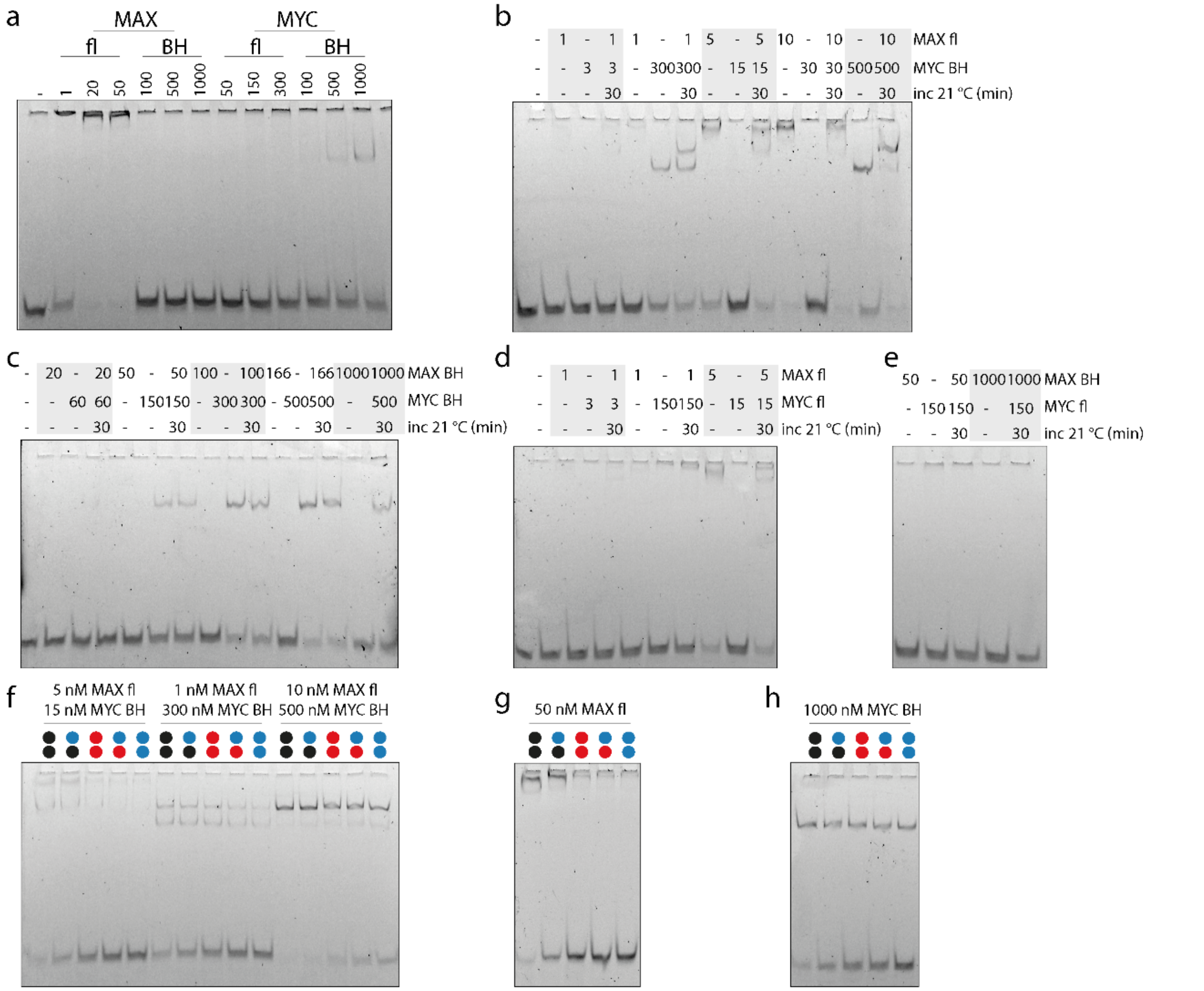


**Figure S20**: **Evaluation of MAX and MYC dimerization.** (a) EMSA analyses of full-length (fl) and bHLHZ (BH) MAX and MYC binding to 2 nM unmodified E-box probe. Protein concentrations are indicated in nM. (b) EMSA analyses of individual MAX fl and MYC BH and potential dimer (30 min incubation of both proteins at 21 °C) binding to 2 nM E-box probe. Protein concentrations are indicated in nM. (c) EMSA analyses of individual MAX BH and MYC BH and potential dimer (30 min incubation of both proteins at 21 °C) binding to 2 nM E-box probe. Protein concentrations are indicated in nM. (d) EMSA analyses of individual MAX fl and MYC fl and potential dimer (30 min incubation of both proteins at 21 °C) binding to 2 nM E-box probe. Protein concentrations are indicated in nM. (e) EMSA analyses of individual MAX BH and MYC BH and potential dimer (30 min incubation of both proteins at 21 °C) binding to 2 nM E-box probe. Protein concentrations are indicated in nM. (f) EMSA analyses of potential MAX fl/ MYC BH dimer after 30 min incubation with indicated protein concentrations to 2 nM modified E-box probe (color code corresponds to **Fig. S17**). (g) EMSA analyses of MAX fl binding to 2 nM modified E-box probe (color code corresponds to **Fig. S17**). (h) EMSA analyses of MYC BH binding to 2 nM modified E-box probe (color code corresponds to **Fig. S17**).

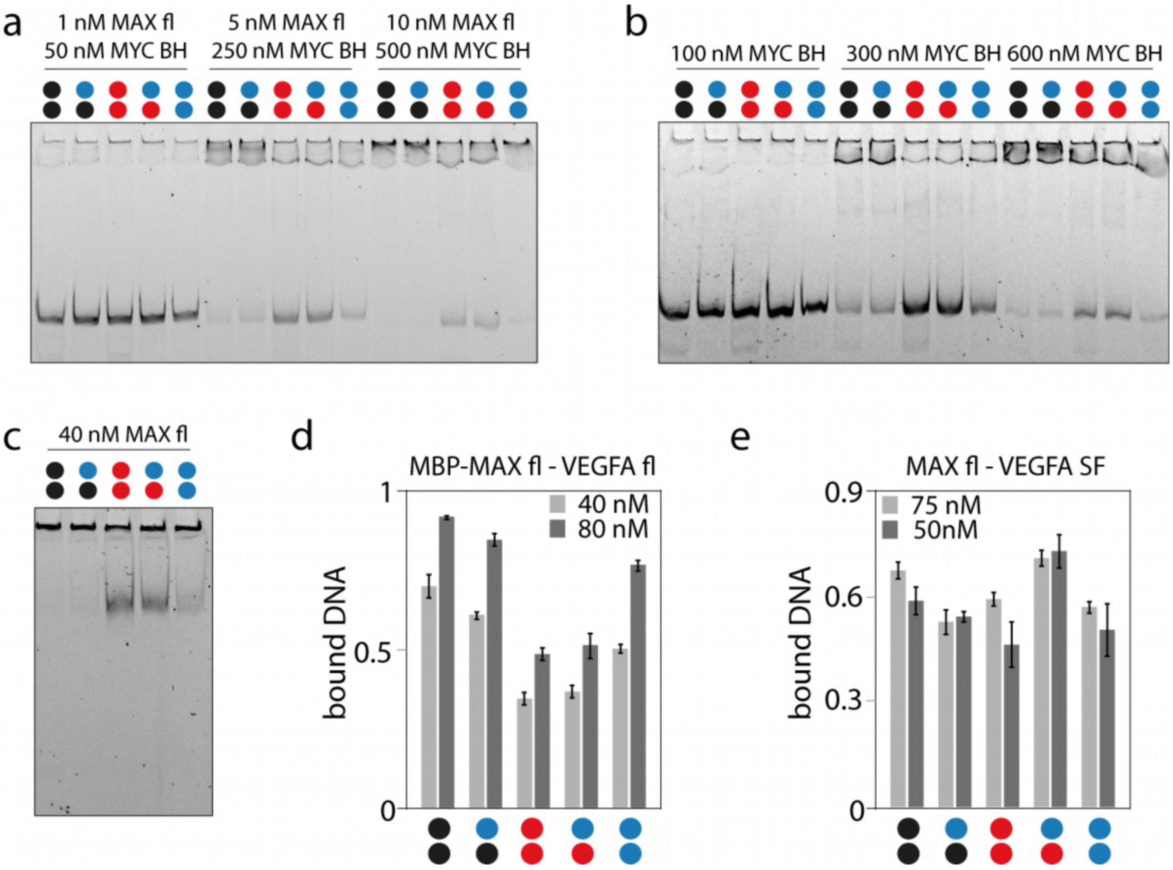


**Figure S21**: **Evaluation of MAX and MYC binding to VEGFA PCR and synthetic probes.** (a) EMSA analyses of full-length (fl) MAX and bHLHZ (BH) MYC binding to 2 nM modified VEGFA probe (color code corresponds to **Fig. S17**). (b) EMSA analyses of MYC BH binding to 2 nM modified VEGFA probe. (c) EMSA analyses of MAX fl binding to 2 nM modified VEGFA probe. (d) Bar diagram from duplicate experiments showing binding preferences of MBP-MAX to differentially modified VEGFA probes. (e) Bar diagram from duplicate experiments showing binding preferences of MBP-MAX to differentially modified VEGFA SF probes (**Table S5** o6107 - o6112).

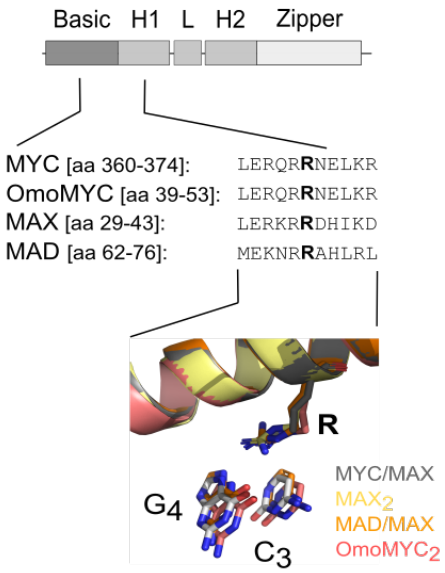


**Figure S22: Superimposed interactions of conserved R367 in the MYC/MAX and related dimers dimer with E-box G4.** Pdb entries (PDB 1NKP, 1AN2, 1NLW, and 5I50).

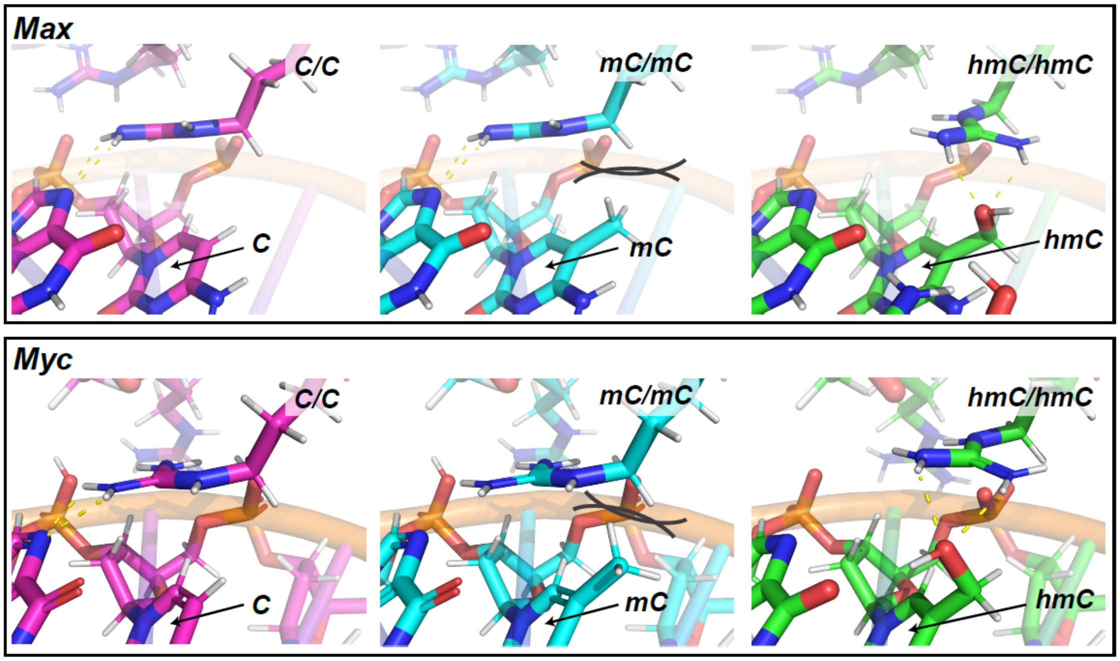


**Figure S23: Visual comparison of the protein-DNA interaction of MAX and MYC with differentially modified CpG dyads.** Shown are structures of the modeled MAX and MYC parts of the MYC/MAX heterodimer in complex with E-box dsDNA (PDB 1NKP) MYC part was modeled exactly as described for MAX in the manuscript. Comparison suggests high structural conservation of hmC binding mode for MAX and MYC.

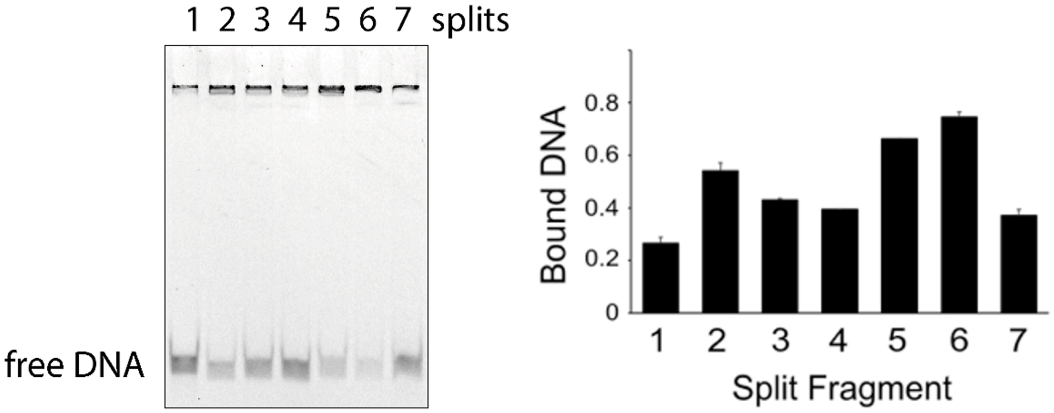


**Figure S24: EMSA-based mapping of $MAX_2$ binding sites in the VEGFA PCR probe.** (left): EMSA gel with purified MBP-MAX fusion protein (500 nM) and unmodified, synthetic VEGFA dsDNA tiling probes covering part of the VEGFA sequence in an overlapping fashion ("splits" (**Table S5** o5688 – o5701), 2 nM). Note that protein-binding lead to later formation of higher MBP-MAX-DNA aggregates in EMSA assays and selectivity can be best judged by inspecting the free DNA bands. (right): bar diagram of duplicate EMSA assays as shown on the left.

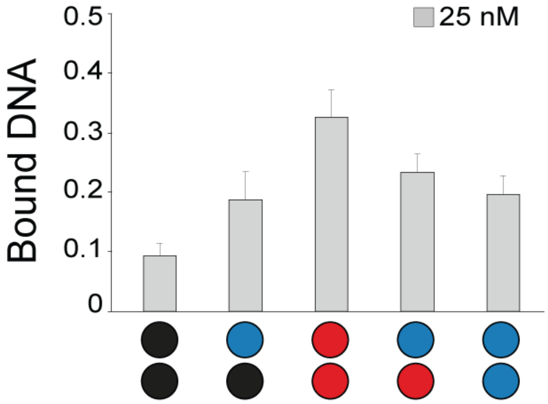


**Figure S25. EMSA analyses of RFX5 binding to X-box 2.** EMSA analyses of 25 nM RFX5 binding to 2 nM modified X-box 2 probes (color code corresponds to **Fig. S17**).

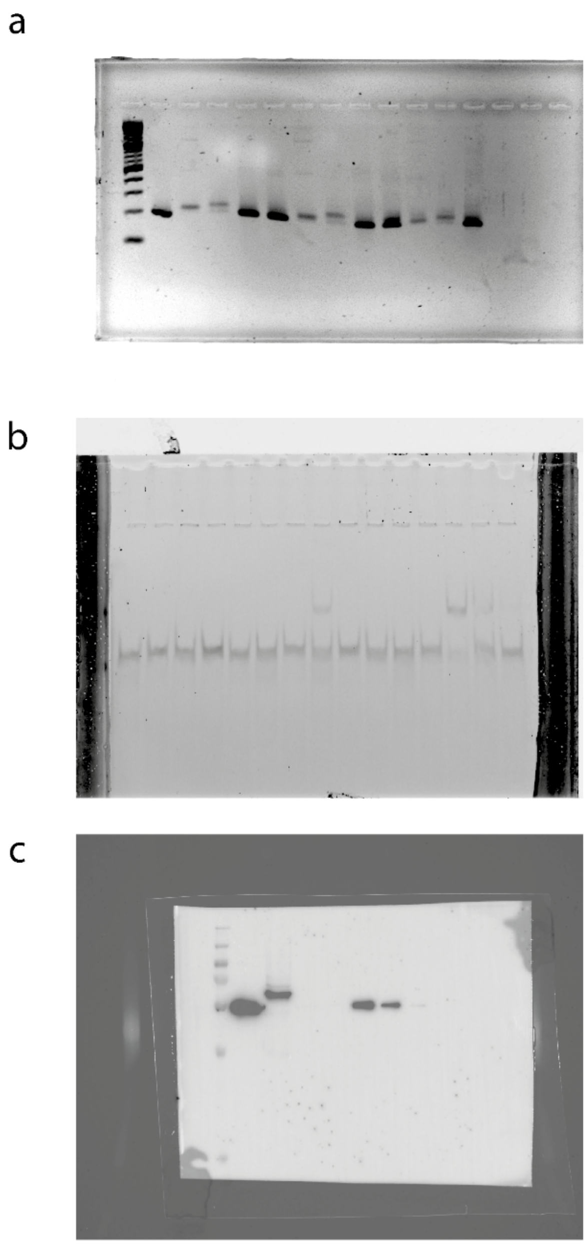

**Figure S26:** Uncropped images related to **Fig. 2** from the manuscript. (a) Agarose gel corresponding to **Fig. 2d**, (b) EMSA gel corresponding to **Fig. 2e**, (c) Western blot corresponding to **Fig. 2f**.

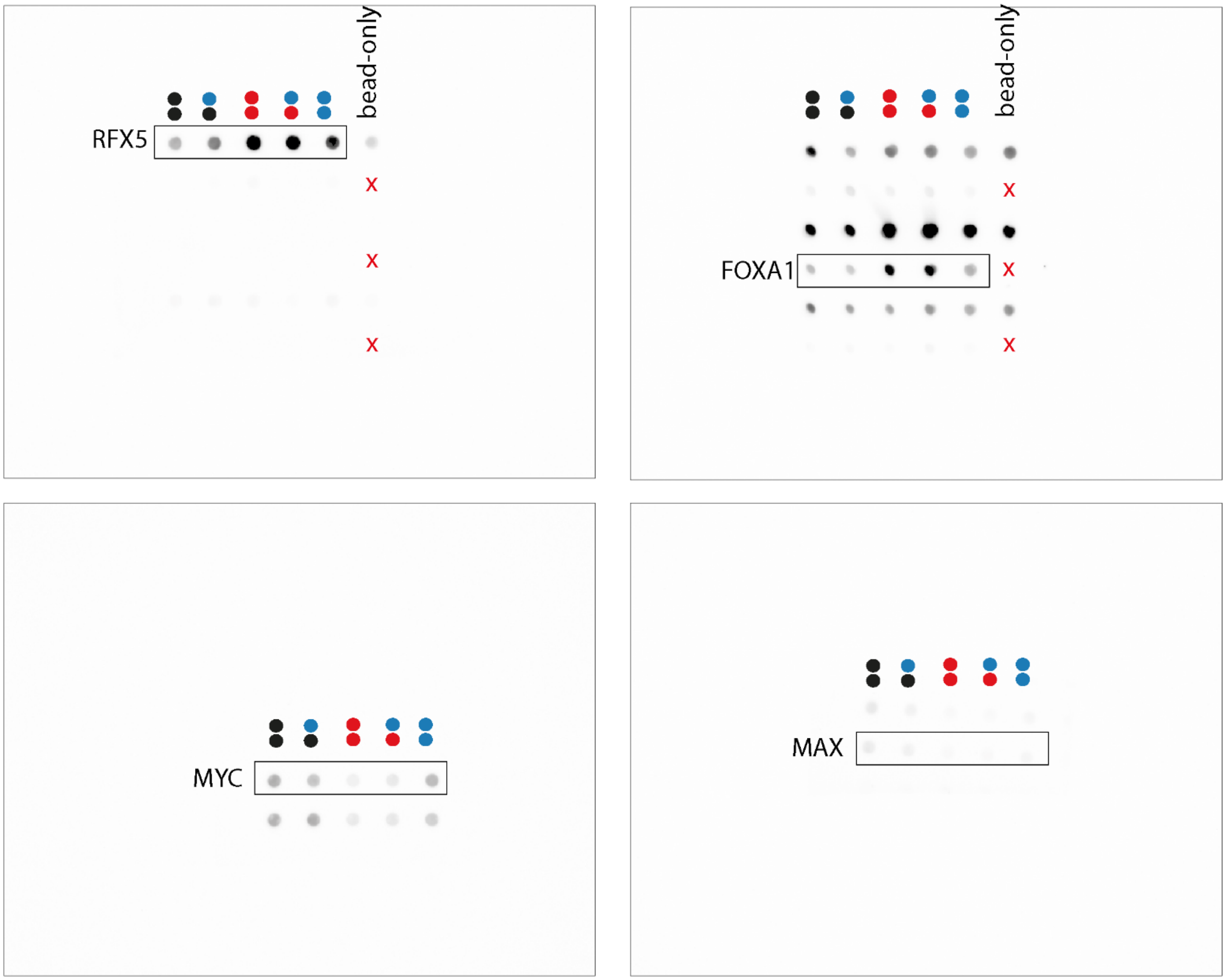


**Figure S27:** Uncropped images related to **Fig. 6** from the manuscript. Dyad modifications and bead-only control (only for RFX5 and FOXA1 blots) are indicated for clarity. Cutouts used for Figure 4 are marked by black boxes, other dot series on same blot may be from other proteins and are not relevant. Red “x“ means that no bead only control was performed for the respective data series.

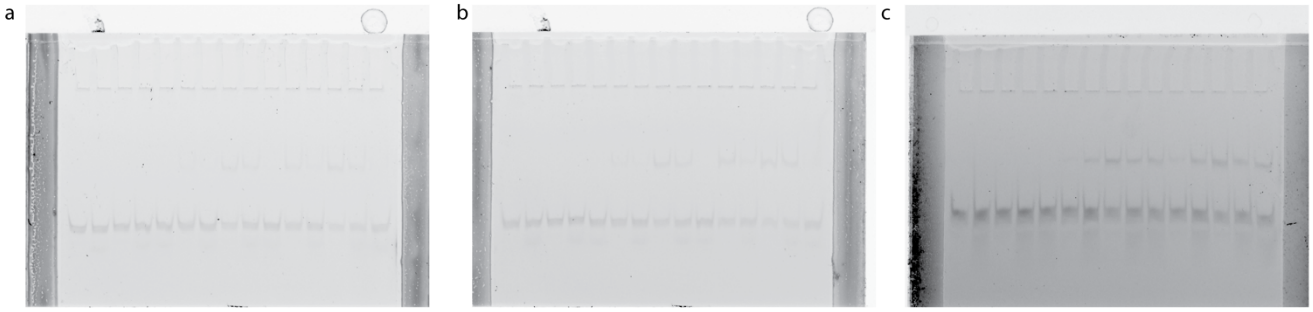


**Figure S28**. Uncropped images related to **Fig. 6j-k**. (a-b) 250, 750 nM RFX5 and X box probe 1 in duplicates, (c) 25 nM RFX5 and X box probe 2 in duplicates used for analysis.

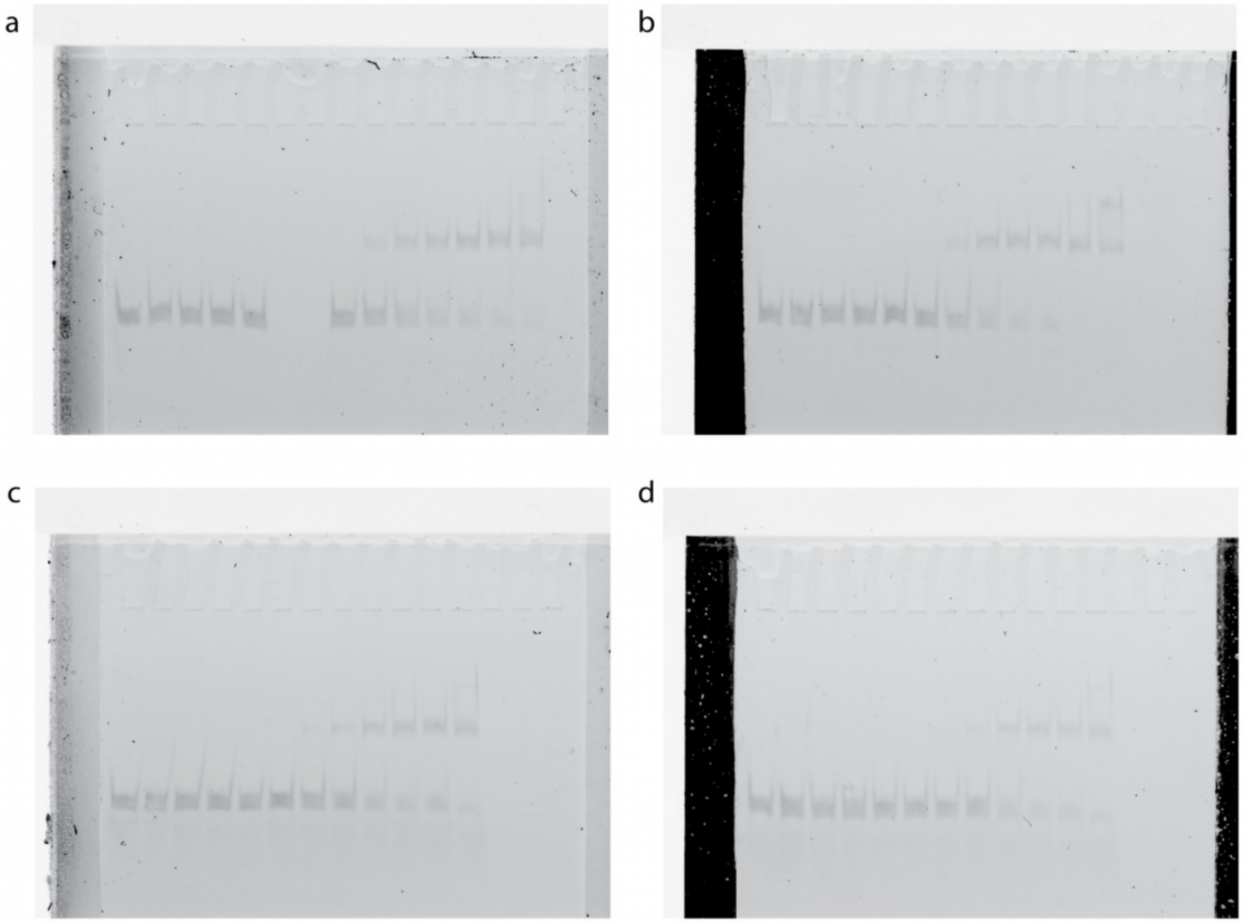


**Figure S29.** Uncropped images related to **Fig. 6l** in the manuscript. (a-b) RFX5 binding to X-box-2 physiological mC/mC and (c-d) hmC/mC. Duplicates shown were used for analysis and $K_D$ calculations.

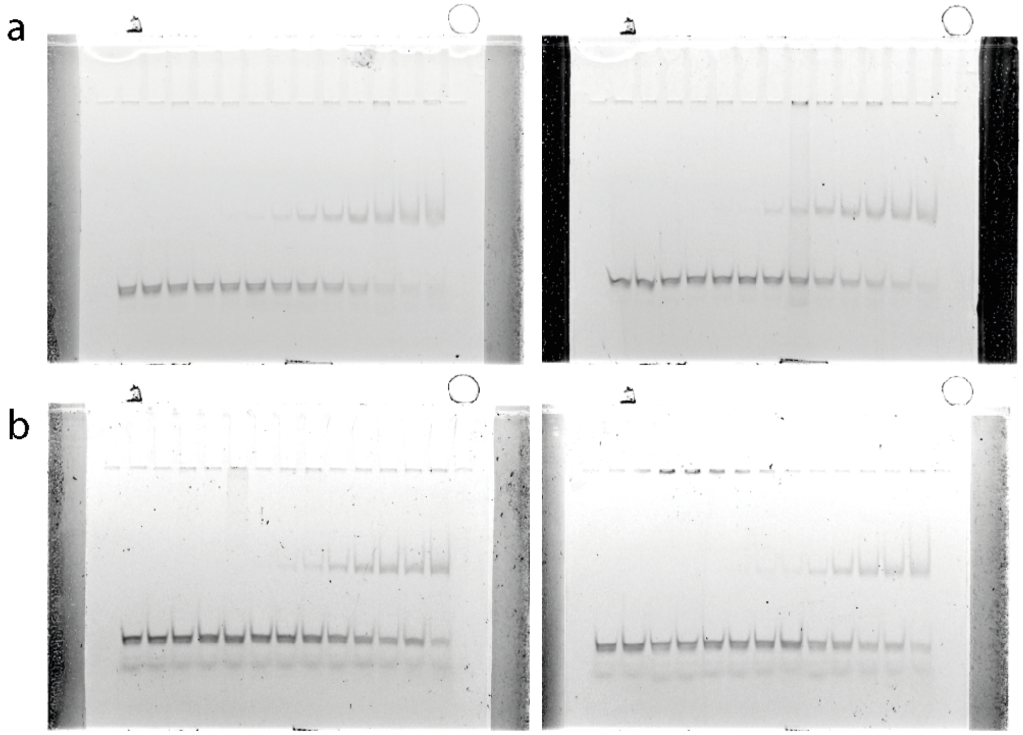

**Figure S30.** Uncropped images related to **Fig. 6l** in the manuscript. (a-b) RFX5 DBD binding to X-box 1 mC/mC (a) and hmC/mC (b). Duplicates shown were used for analysis and $K_D$ calculations.

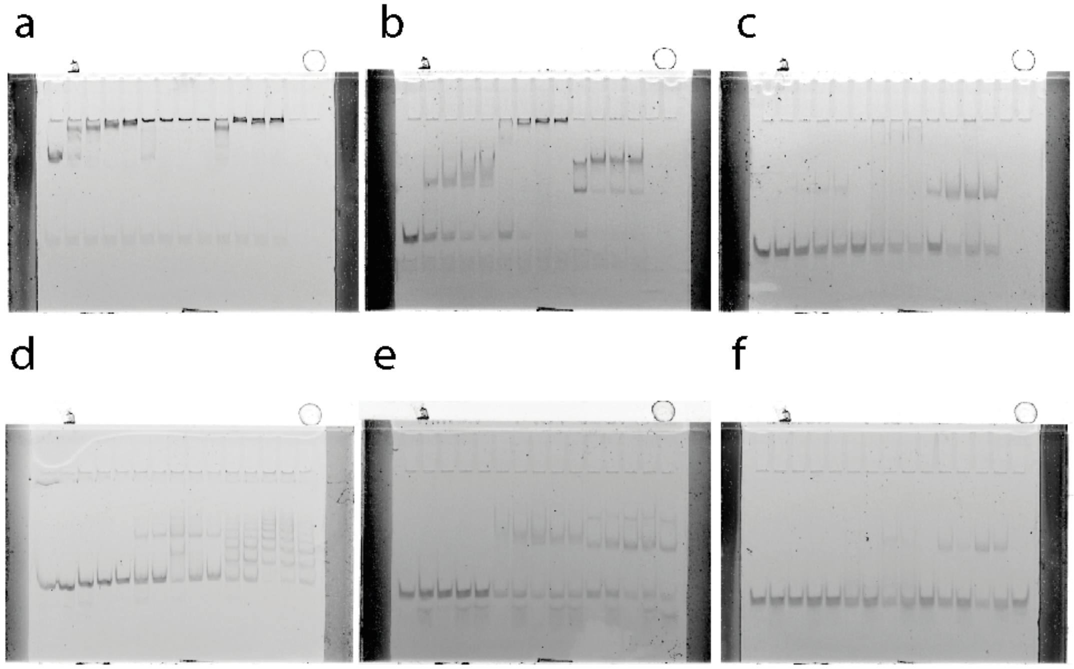

**Figure S31.** Uncropped images related to **Fig. 6** in the manuscript. Concentration series of (a) RFX5 binding to the VEGFA probe, (b) RFX5 binding to the physiological X-box 2 probe and (c) RFX5 binding to the X-box 1 probe. (d) RFX5 binding to diefferentially modified VEGFA probes. (e) RFX5 binding to diefferentially modified X-box 1 probes. (f) RFX5 binding to diefferentially modified X-box 2 probes.

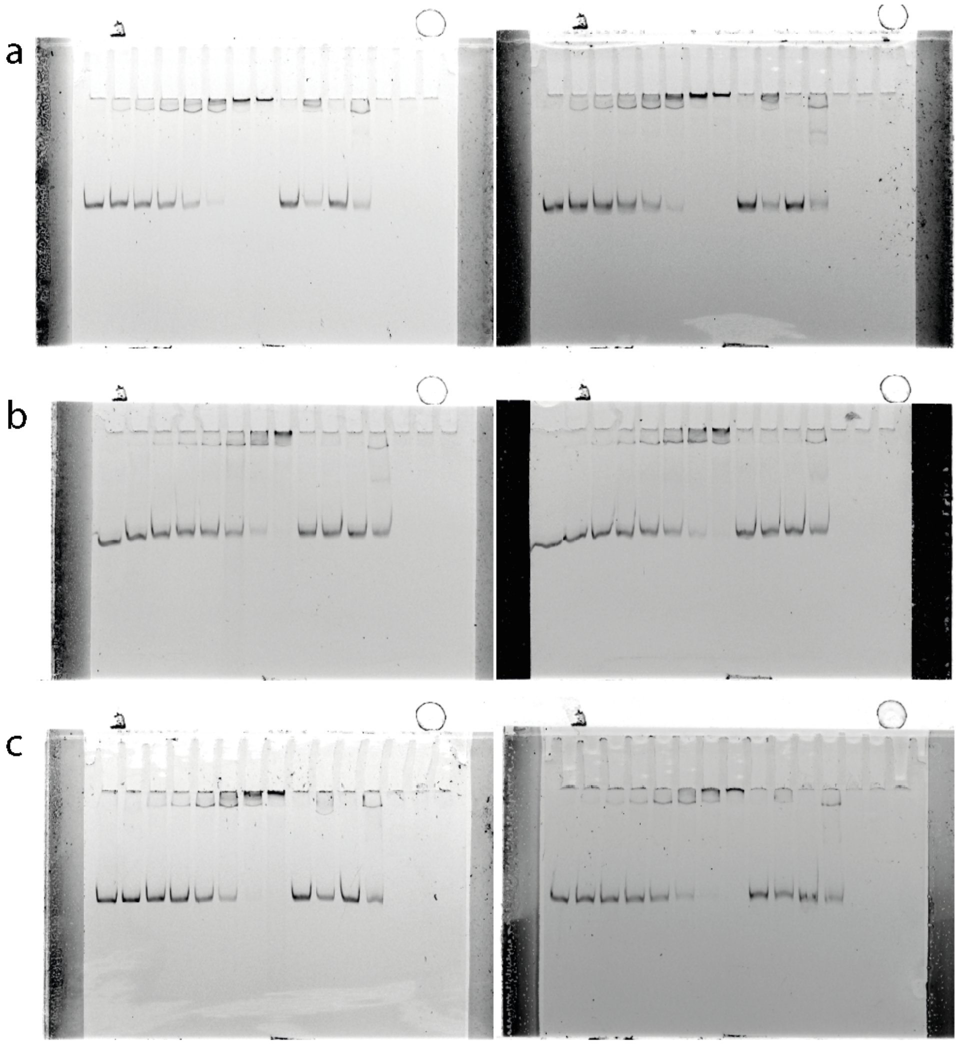


**Figure S32.** Uncropped images related to **Fig 6e** in the manuscript. (a) MAX fl and MYC bHLHZ binding to VEGFA C/C, (b) MAX fl and MYC bHLHZ binding to VEGFA mC/mC, (c) MAX fl and MYC bHLHZ binding to VEGFA hmC/hmC. Duplicates shown were used for analysis and $K_D$ calculations.

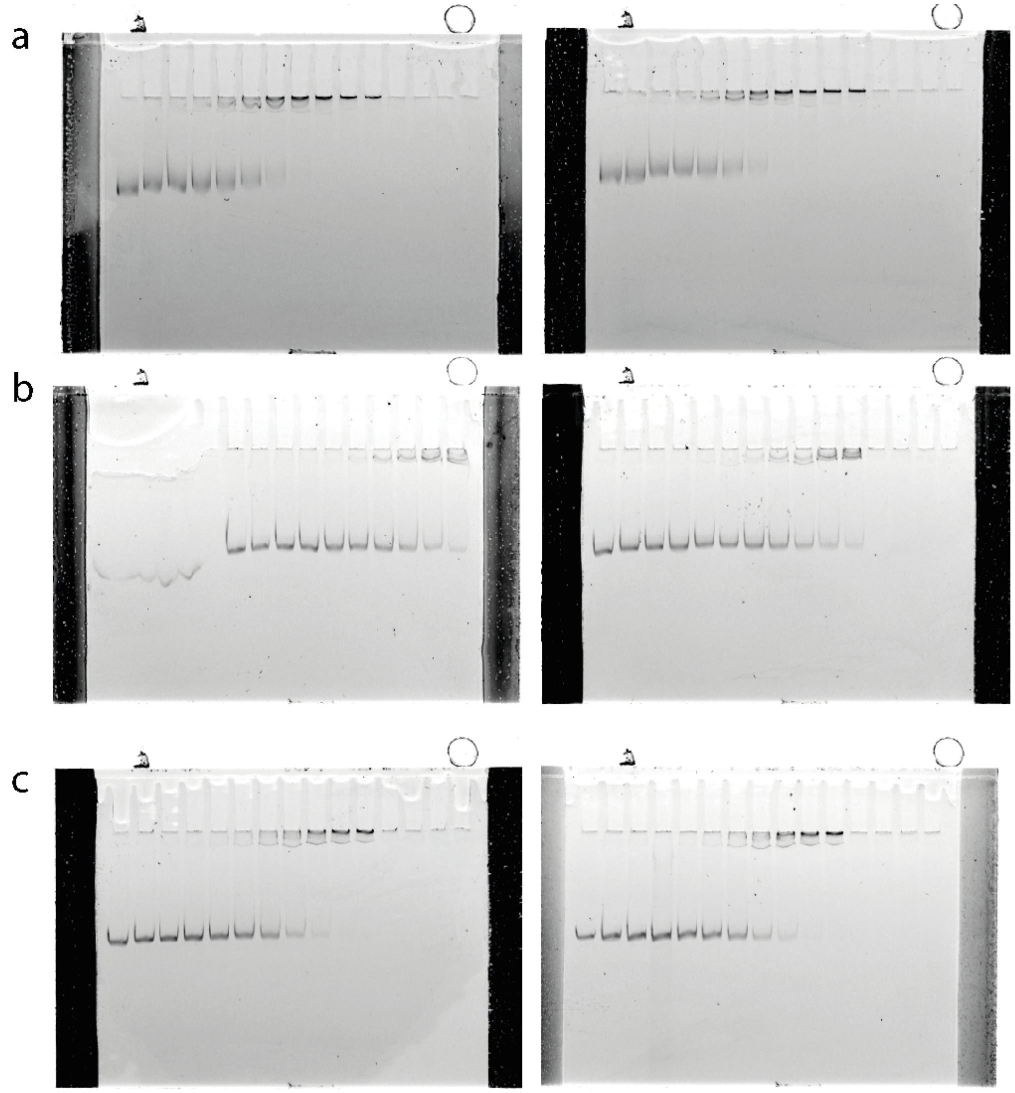

**Figure S33.** Uncropped images related to **Fig. 6e** in the manuscript. (a) MAX fl homodimer binding to VEGFA C/C, (b) MAX fl and MYC bHLHZ binding to VEGFA mC/mC, (c) MAX fl and MYC bHLHZ binding to VEGFA hmC/hmC. Duplicates shown were used for analysis and kD calculations.

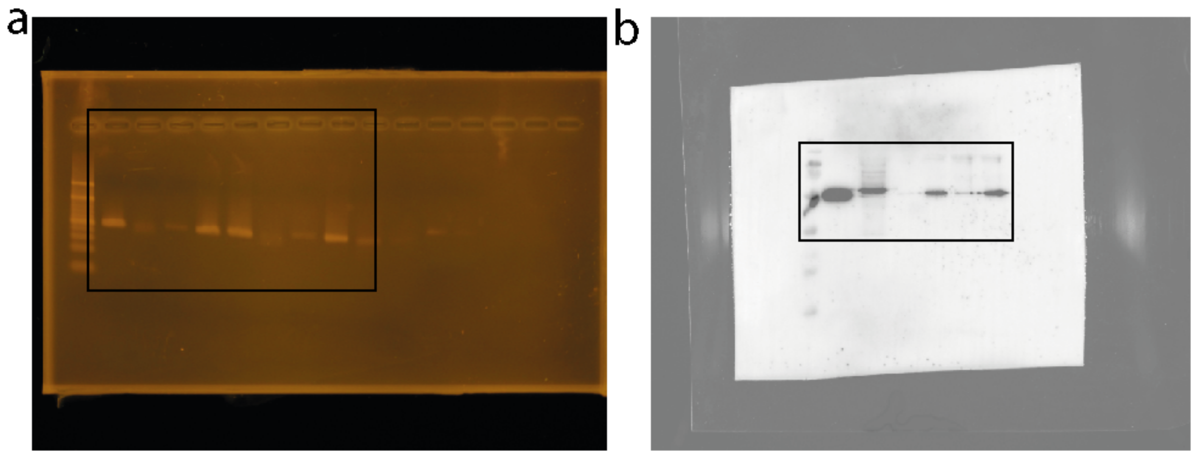

**Figure S34:** Uncropped images related to **Fig. S1.** (a) Agarose gel corresponding to Fig S1a, (b) EMSA gel corresponding to Fig. S1b. Cutouts used for figure are marked by a black box,

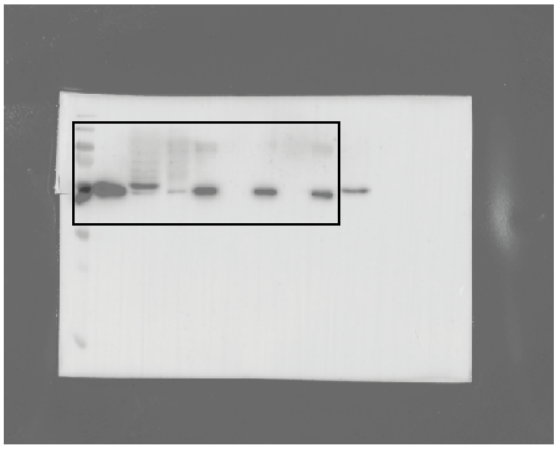

**Figure S35:** Uncropped agarose gel related to **Fig. S2**. Cutout used for figure is marked by a black box,

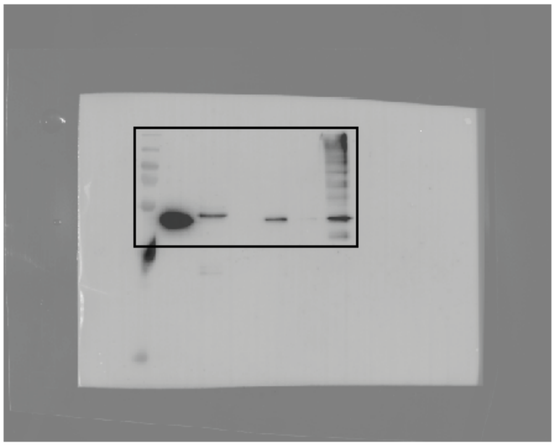

**Figure S36:** Uncropped Western blot related to **Fig. S3**. Cutout used for figure is marked by a black box,

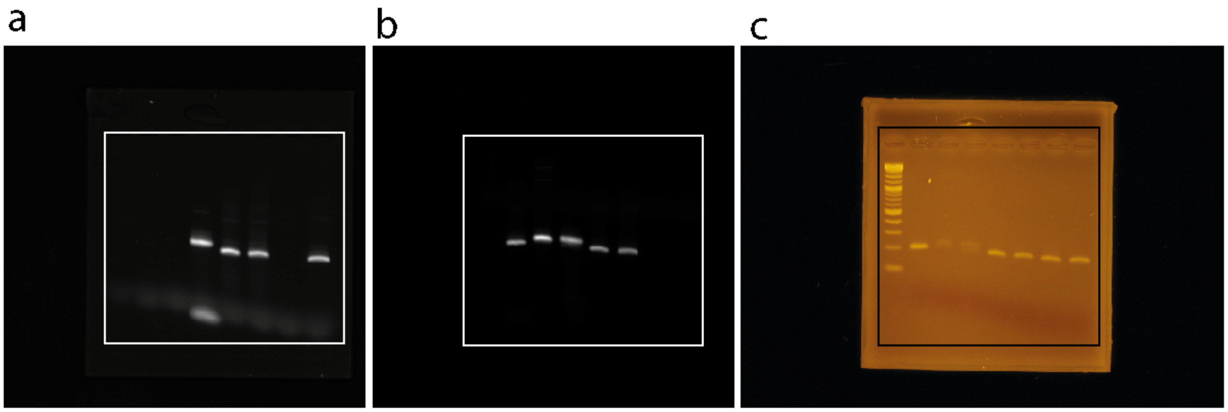


**Figure S37:** Uncropped images related to **Fig. S6**. (a) Agarose gel imaged in Cy3 channel. (b) Agarose gel imaged in Cy5 channel. (c) Agarose gel stained with EtBr. Cutouts used for figure are marked by a box,

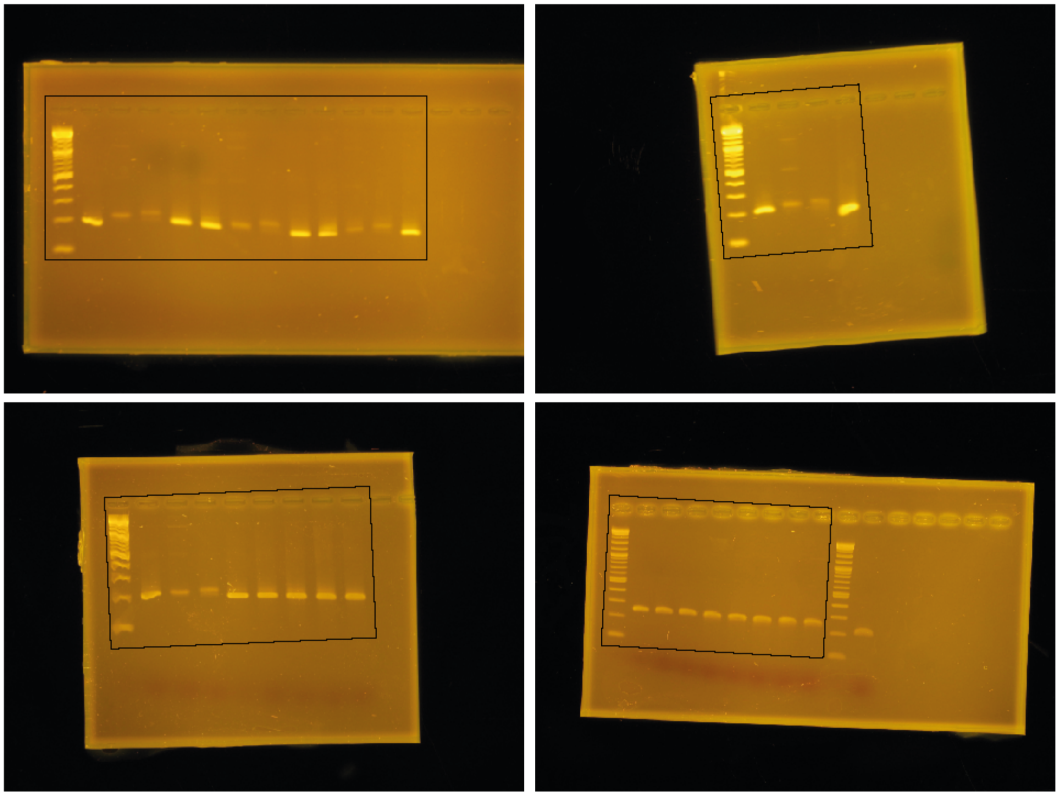

**Figure S38:** Uncropped images related to **Fig. S8**. Cutouts used for figure are marked by a black box, a

b

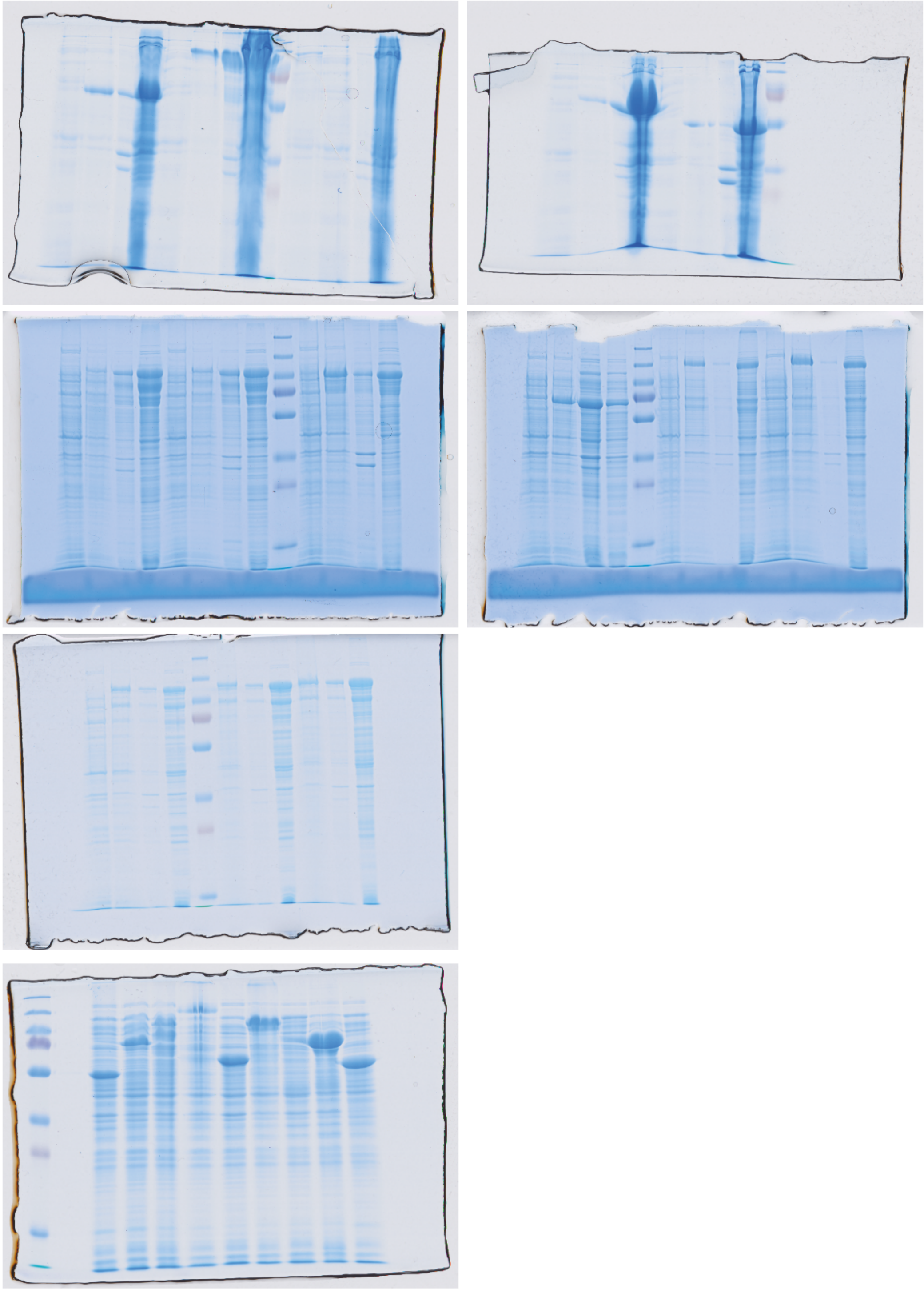

**Figure S39:** Uncropped images related to **Fig. S15**. (a) Coomassie-stained SDS gels corresponding to Fig S15b. (b) Coomassie-stained SDS gel corresponding to Fig. S15c.

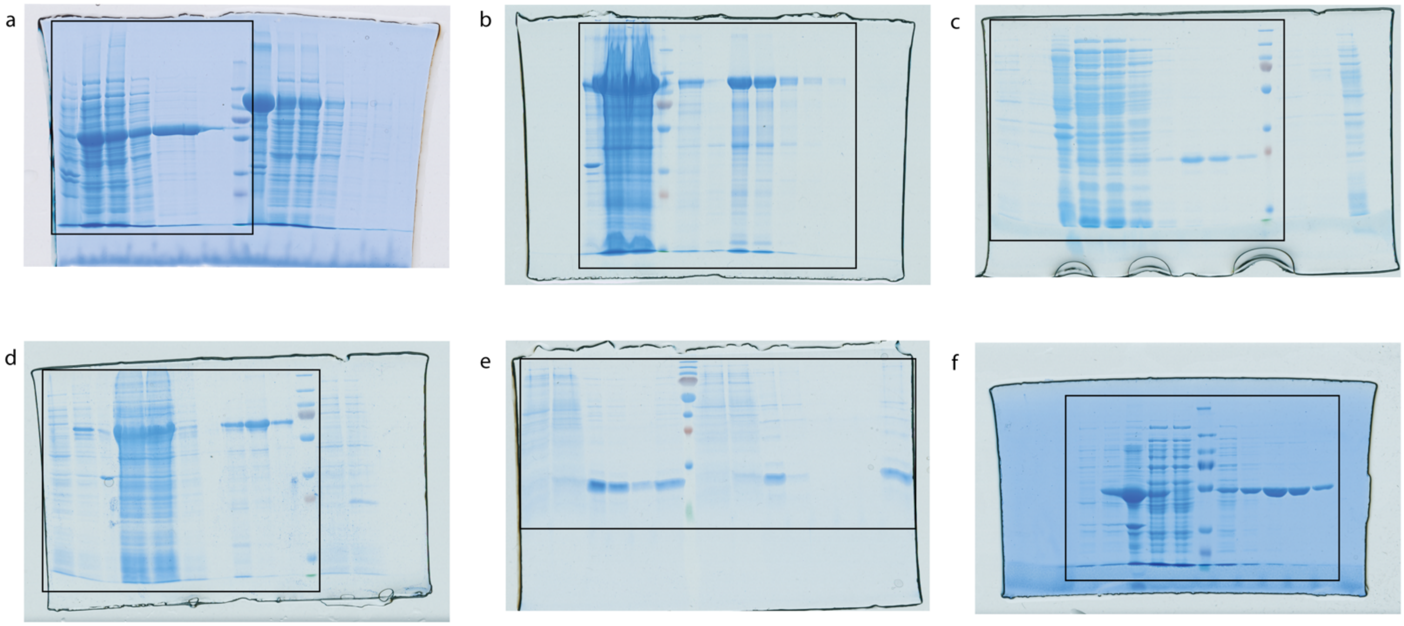


**Figure S40:** Uncropped images related to **Fig. S16**. Coomassie stained SDS gel corresponding to Fig. S16a-g, numbers correspond to numbers in Fig. S16. Cutouts used for Fig. S16 are marked by black boxes,

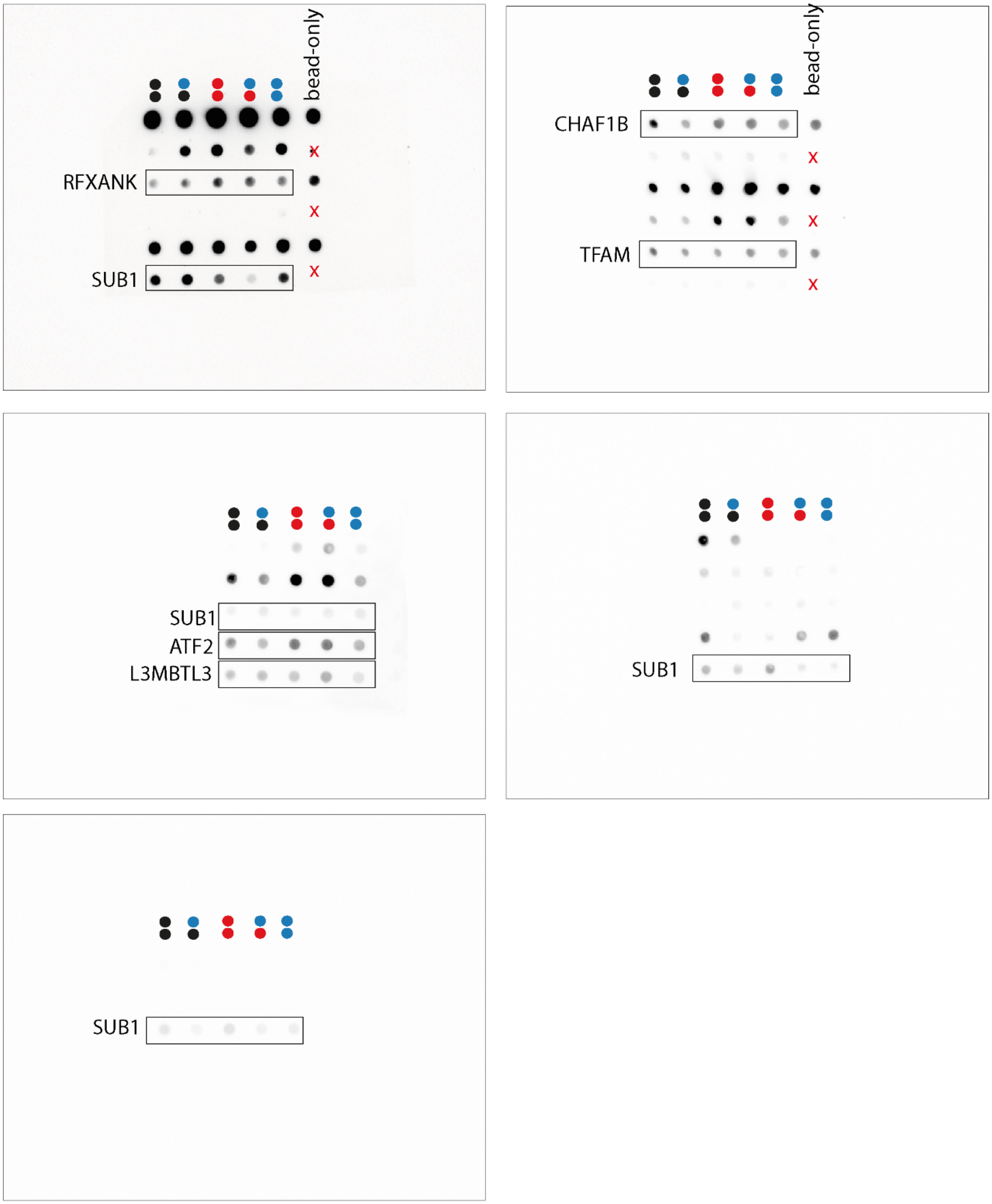


**Figure S41:** Uncropped images related to **Fig. S17**. Dyad modifications and bead-only controls are indicated for clarity. Cutouts used for Fig. S17 are marked by black boxes, other dot series on same blot may be from other proteins and are not relevant. Red "x" means that no bead only control was performed for the respective data series.

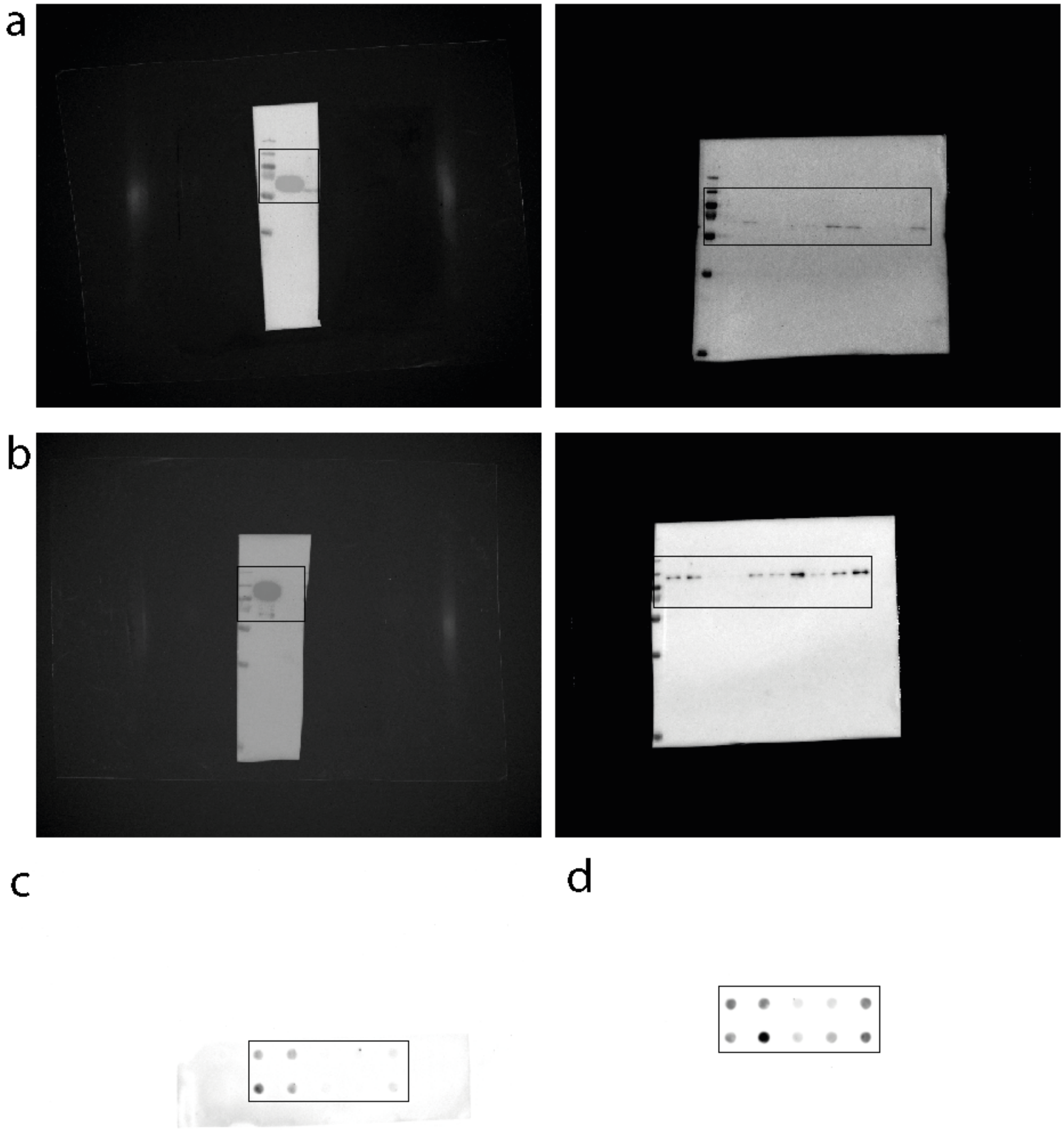

**Figure S42:** Uncropped images related to **Fig. S18**. (a) Western blots corresponding to Fig. S18a, (b) Western blots corresponding to Fig. S18c, (c) Dot blot corresponding to Fig. S18b. (d) Dot blot corresponding to Fig. S18d. Cutouts used for Fig. S18 are marked by black boxes.

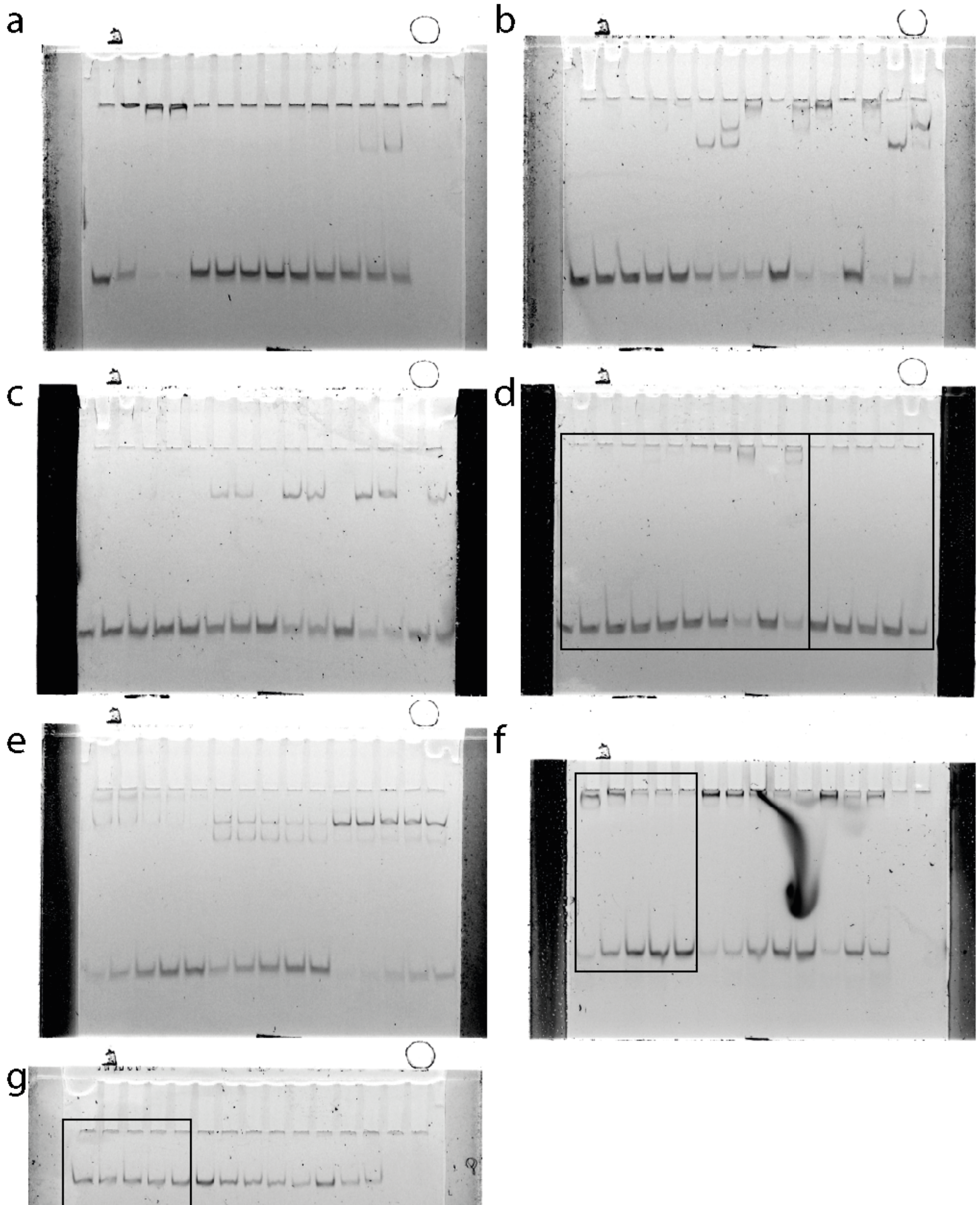


**Figure S43:** Uncropped images related to **Fig. S20**. (a) EMSA gel corresponding to Fig. S20a. (b) EMSA gel corresponding to Fig. S20b. (c) EMSA gel corresponding to Fig. S20c. (d) EMSA gel corresponding to Fig. S20d and e. (e) EMSA gel corresponding to Fig. S20f. (f) EMSA gel corresponding to Fig. S20g. (g) EMSA gel corresponding to Fig. S20h. Cutouts used for Fig. S20 are marked by black boxes.

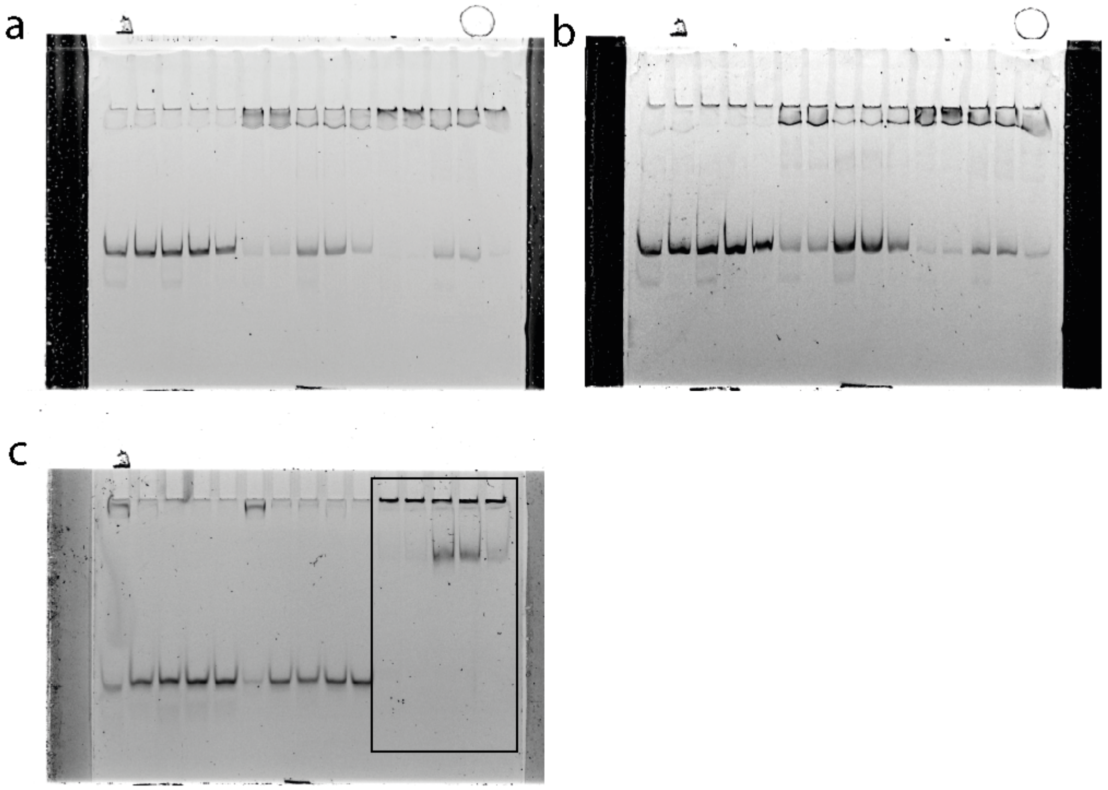


**Figure S44:** Uncropped images related to **Fig. S21**. (a) EMSA gel corresponding to Fig. S20a. (b) EMSA gel corresponding to Fig. S20b. (c) EMSA gel corresponding to Fig. S20c. Cutouts used for Fig. S19 are marked by black boxes.

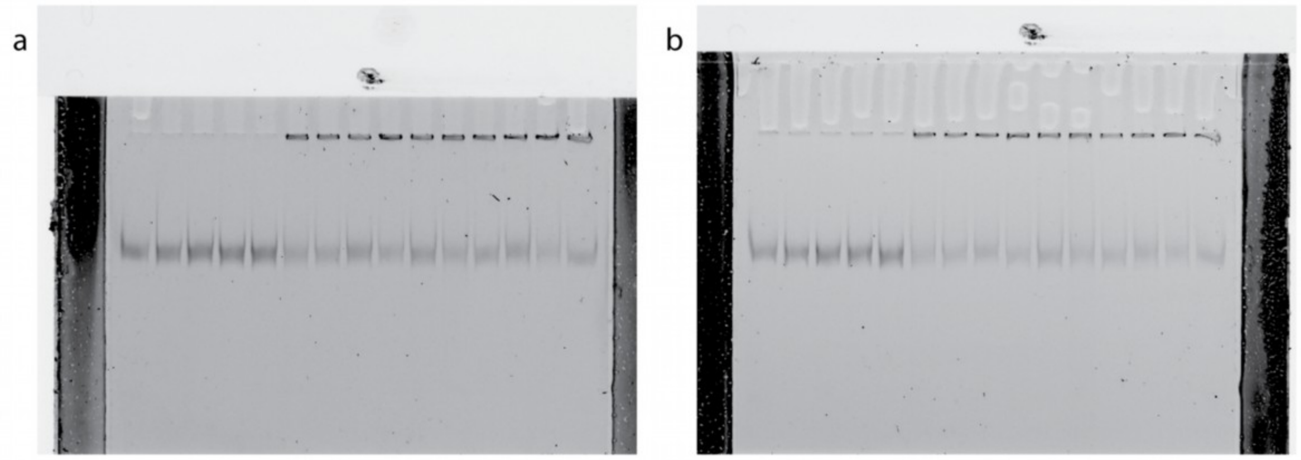


**Figure S45.** Uncropped images related to **Fig. S21e**. Experiments were performed in duplicates, (a) 75 nM and (b) 50 nM MAX and 2 nM VEGFA SF (a and b).
